# Large-scale cerebrospinal fluid proteomics identifies diagnostic biomarker candidates across the clinico-pathological continuum of frontotemporal lobar degeneration

**DOI:** 10.64898/2026.09.29.755483

**Authors:** Rowan Saloner, Louisa Cornelis, Joshua Downer, Connor Dietz, Brandon Chan, Michelle Hughes, Molly Olzinski, Yann Cobigo, Argentina Lario-Lago, Hilary Heuer, Karly Cody, Madeline Wood, Katheryn A.Q. Cousins, David Wolk, Edward B. Lee, David Irwin, Ahmed Abdelhak, Bradley F. Boeve, Gil D. Rabinovici, Renaud La Joie, Lea T. Grinberg, Salvatore Spina, Emily W. Paolillo, Adam Staffaroni, Lawren VandeVrede, Peter A. Ljubenkov, Maria Luisa Gorno-Tempini, Howard J. Rosen, Jennifer S. Yokoyama, Joel H. Kramer, Kaitlin B. Casaletto, Bruce Miller, William W. Seeley, Julio C. Rojas, Adam L. Boxer

## Abstract

Frontotemporal lobar degeneration (FTLD), predominantly comprised of tau and TDP-43 proteinopathies, is a common cause of early-onset dementia. Biomarkers that reliably classify FTLD subtypes are limited, thereby hindering in-vivo diagnosis and clinical trial stratification. We conducted large-scale, multi-platform cerebrospinal fluid proteomics in 566 individuals, including 274 FTLD-spectrum (48 autopsy-confirmed FTLD-TDP, 90 autopsy-confirmed FTLD-tau), 171 Alzheimer’s disease, and 95 controls. In neuropathological subtype contrasts, elevated matrix metalloproteinase MMP10 classified Pick’s disease (PiD) versus other subtypes with near-perfect accuracy across proteomic platforms. We further validated CSF MMP10 as a potential PiD diagnostic biomarker via a targeted immunoassay in an independent, autopsy-confirmed FTLD cohort. Post-synaptic proteins HOMER1 and DLG4 exhibited elevations across FTLD-TDP subtypes (A, B, and C) versus FTLD-tau. Network analysis identified RNA metabolism and pro-inflammatory signatures in FTLD-TDP, and cytoskeletal and heat shock signatures in progressive supranuclear palsy. These findings reveal candidate diagnostic biomarkers and proteomic pathways underlying FTLD clinico-pathological heterogeneity.

## MAIN

Frontotemporal lobar degeneration (FTLD) is a leading pathological cause of early-onset dementia syndromes that encompass a heterogeneous spectrum of behavioral, language, and movement phenotypes. FTLD clinical syndromes include behavioral variant frontotemporal dementia (bvFTD), primary progressive aphasia (PPA), corticobasal syndrome (CBS), progressive supranuclear palsy (PSP), and frontotemporal dementia with motor neuron disease (FTD/MND)^1^. Approximately half of FTLD is associated with deposition of pathogenic tau (FTLD-tau) with the other half associated with pathogenic TDP-43 (FTLD-TDP), and a small fraction due to other rarer FTLD pathologies (e.g., FUS)^2^. FTLD clinico-pathological relationships are complex and frequently overlapping. For example, Pick’s disease (PiD), a distinct 3R-tau subtype linked to bvFTD can also cause nonfluent and semantic variants of primary progressive aphasia (nfvPPA, svPPA), while bvFTD itself can arise from most FTLD-tau and FTLD-TDP subtypes. Consequently, clinical syndrome alone is often insufficient to determine underlying molecular pathology in living patients. The absence of in-vivo biomarkers capable of predicting underlying pathology remains a major barrier to molecular diagnosis and the development and evaluation of pathology-directed therapies^3^.

Recent technological advances in large-scale proteomics enable quantification of thousands of proteins in human biofluids, thereby facilitating diagnostic biomarker discovery in clinical cohorts. Large-scale proteomics have revealed blood and cerebrospinal fluid (CSF) proteomic signatures in late-onset Alzheimer’s disease and familial forms of FTLD^4–8^, in which underlying FTLD-TDP (*C9orf72*, *GRN*) and FTLD-tau (*MAPT*) pathology is known. However, the greatest unmet clinico-diagnostic need exists in sporadic FTLD, where clinical diagnosis remains uncertain and substantial pathological heterogeneity exists even within FTLD-TDP and FTLD-tau molecular classes. This gap has been historically difficult to address because sporadic cases with both antemortem biofluid collection and postmortem autopsy confirmation are scarce, limiting systematic proteomic studies to identify FTLD subtype-specific fluid biomarkers.

Here we apply large-scale, multi-platform CSF proteomics (SomaScan v4.1 and Olink Explore HT) in a large cohort of FTLD cases and age-matched, early-onset AD cases (total N=566). FTLD cases included autopsy-confirmed patients spanning six primary FTLD-TDP and FTLD-tau pathological subtypes. In subtype-specific diagnostic biomarker screening, elevated matrix metalloproteinase protein 10 (MMP10) classified PiD vs. all other pathological subtypes with near-perfect accuracy across proteomic platforms. We further validated CSF MMP10 as a diagnostic biomarker candidate for PiD via a targeted immunoassay across multiple cohorts and identified up to 6-fold higher MMP10 concentrations in PiD vs. other FTLD subtypes. Beyond MMP10, our proteomic screen identified additional proteins with strong classification accuracy for FTLD molecular classes and subtypes, providing additional candidate biomarkers for future validation. In parallel, network analyses revealed coordinated proteomic alterations in RNA metabolism, immune signaling, proteostasis, and synaptic signatures of FTLD and AD. Together, our study identifies robust fluid biomarker candidates for differential diagnosis across the FTLD clinico-pathological continuum and highlights underlying biological pathways linked to neurodegenerative disease heterogeneity.

## RESULTS

### Cohort characteristics and preprocessing

We conducted large-scale CSF proteomics in 566 individuals from the UCSF Memory and Aging Center (see **Fig. 1a** for study design). CSF was assayed via aptamer-based proteomics (SomaScan v4.1) in 498 individuals and proximity-extension assay proteomics (Olink Explore HT) in 286 individuals, with samples from 218 individuals assayed on both platforms. Participants underwent detailed clinico-diagnostic evaluations and were classified along the FTLD clinical spectrum (N=274), AD clinical spectrum with biomarker confirmation (N=171), cognitively unimpaired controls (CTL; N=95), or “Other” (N=26; e.g., Parkinson’s disease, primary psychiatric). FTLD and AD cases were subdivided into autopsy and clinical only cohorts. The autopsy cohort included cases with antemortem CSF and primary neuropathological diagnoses of FTLD-TDP (N=48), FTLD-tau (N=90), or AD (N=50), well-matched by age, sex, and clinical severity at time of CSF collection (**Fig. 1b-d**). FTLD molecular classes were further subdivided into their constituent FTLD-tau (22 PiD, 28 CBD, 33 PSP) or FTLD-TDP (16 TDPa, 12 TDPb, 16 TDPc; **Fig. 1e-f**) subtypes, which were all sporadic with exception of 14 *GRN*-TDPa, 1 *C9orf72*-TDPa, and 4 *C9orf72*-TDPb. AD biomarker-negative CTL (N=75) were included as a reference group in autopsy cohort contrasts. The clinical only cohort consisted of clinically-diagnosed cases with primary FTLD syndromes (N=127) or AD syndromes with in-vivo AD biomarker positivity (N=121). Demographic, clinical, CSF biomarker, and neuropathological characteristics are provided for samples in the full cohort (**Supplementary Table 1)** as well as autopsy-confirmed molecular classes (**Supplementary Table 2)** and FTLD subtypes (**Supplementary Table 3).**

**Figure 1.**
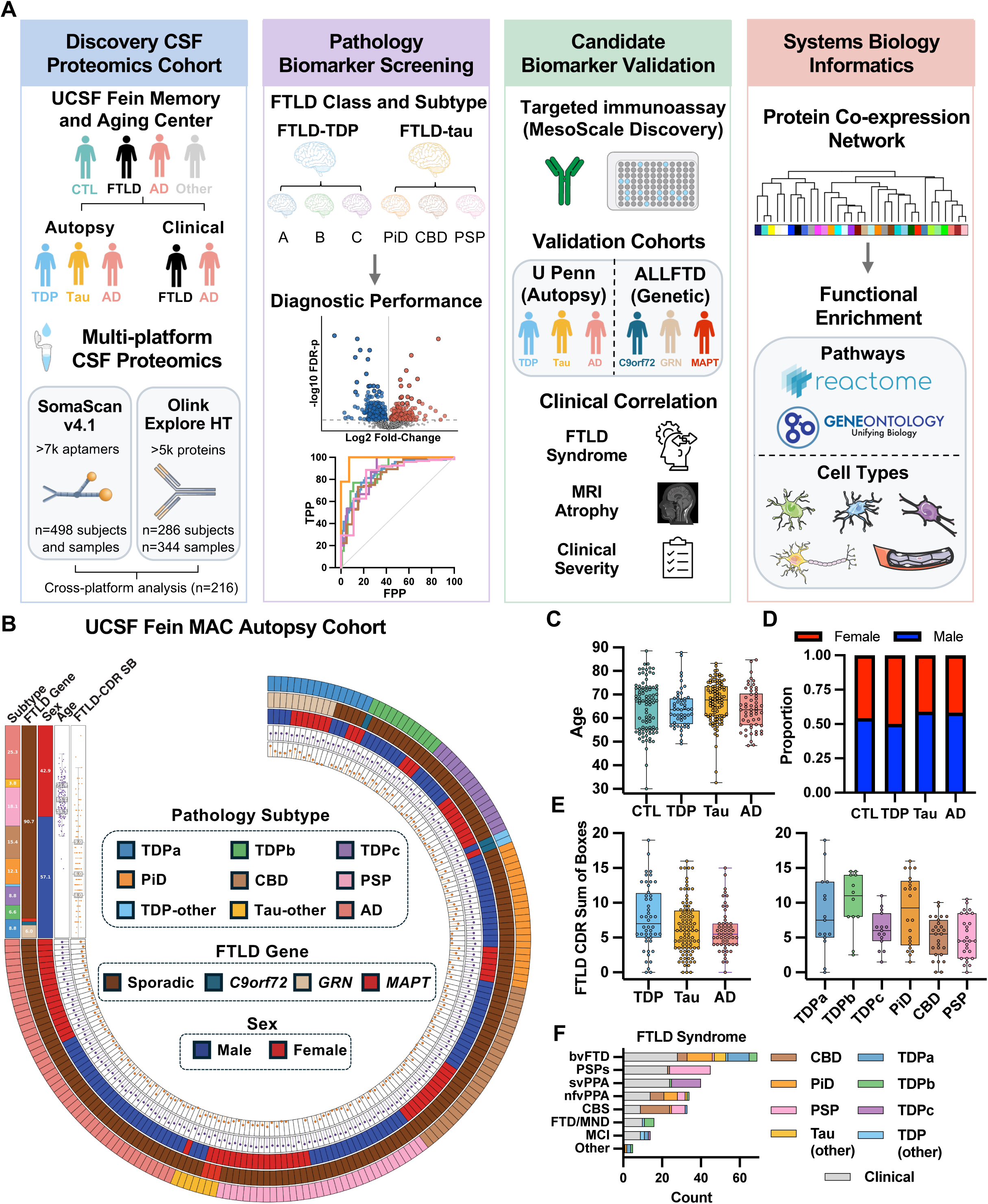
Study design and clinico-pathological characteristics of UCSF Fein Memory and Aging Center discovery cohort. **a**, CSF proteomics were generated in a discovery cohort of frontotemporal lobar degeneration (FTLD), Alzheimer’s disease (AD), and control (CTL) cases from the UCSF Fein Memory and Aging Center. Samples were assayed via SomaScan v4.1 (>7,000 aptamers; n = 498 subjects and samples) and Olink Explore HT (>5,000 proteins; n = 286 subjects and 344 samples), with a subset of samples assayed on both platforms. Differentially abundant analytes distinguishing FTLD molecular class (Tau, TDP) and pathological subtype were assessed for diagnostic performance, then validated by targeted immunoassay (Meso Scale Discovery) and examined in independent autopsy (University of Pennsylvania) and familial FTLD (ALLFTD) cohorts. Weighted protein co-expression network analysis assembled proteins across platforms into modules, which were examined for enrichment of biological pathways (Reactome, Gene Ontology), brain cell types, and associations with pathological subtypes and disease severity outcomes. **b,** Composition of the autopsy cohort. Radial tracks denote, from outer to inner, 1) neuropathological subtype (TDPa, TDPb, TDPc, Pick’s disease [PiD], corticobasal degeneration [CBD], progressive supranuclear palsy [PSP], TDP-other, tau-other, AD); 2) FTLD genetic mutation status (sporadic, *C9orf72*, *GRN*, *MAPT*); 3) sex; 4) age; and 5) CDR®+NACC-FTLD sum of boxes (FTLD-CDR Sum of Boxes) at CSF collection. **c,** Age at CSF collection and **d,** proportion female/male by molecular class (CTL, FTLD-TDP, FTLD-tau, AD). **e,** FTLD-CDR Sum of Boxes at CSF collection by molecular class (left) and neuropathological subtype (right). **f,** Number of participants meeting criteria for each clinical FTLD syndrome (behavioral variant frontotemporal dementia [bvFTD], progressive supranuclear palsy spectrum [PSPs], semantic variant primary progressive aphasia [svPPA], nonfluent variant primary progressive aphasia [nfvPPA], corticobasal syndrome [CBS], frontotemporal dementia with motor neuron disease [FTD/MND], mild cognitive impairment [MCI], other), colored by neuropathological subtype for autopsy-confirmed cases or labeled “Clinical” where diagnosis was based on clinical criteria alone. Box plots (**c**, **e**) show the median (center line), 25th and 75th percentiles (box bounds), and range (whiskers), with individual data points overlaid.

Prior to primary analysis, we applied an established quality control (QC) pipeline to both platform datasets^7,9^. QC included removal of outlier samples (**Supplementary Fig. 1a**) with aberrant network connectivity (SomaScan: N=7; Olink: N=1) and removal of assays (SomaScan aptamers or Olink proteins) with clear bimodal distributions that violated parametric assumptions and high rates (>75%) of values below limit of detection (LOD; >3SD above background; **Supplementary Fig. 1b**). A total of 3,713 SomaScan aptamers and 1,293 Olink proteins were retained after analyte filtering (**Supplementary Tables 4-5**). The majority of assays that failed QC were due to poor detection above background, consistent with prior CSF SomaScan and Olink studies showing reduced detection of proteins in CSF on assays originally optimized for plasma^7,10^. Proteins passing QC were more frequently differentially abundant between all disease cases combined vs. CTL than those failing QC (SomaScan: 37% vs. 5%; Olink: 44% vs. 15%; **Supplementary Fig. 1b**), supporting the effectiveness of QC in retaining disease-relevant signal. Further, median cross-platform correlations were largest for matching SomaScan aptamer and Olink protein pairs that passed QC filtering across both platforms (median bicor=0.57) compared to those that failed QC on one or both platforms (median bicor range: 0.00 to 0.07; **Supplementary Table 6, Supplementary Fig. 1c**). After analyte filtering, nuisance covariance from principal component 1 (PC1), blood contamination, age, and sex was estimated in CTL samples with resulting coefficients used to adjust the full protein abundance matrix for each platform^6^. Variance partition plots confirmed that this approach preserved proteomic variance attributable to diagnostic status while removing nuisance covariance (**Supplementary Fig. 1d)**. A final winsorization step was applied to remove undue influence of individual outlier measurements. Outlier-removed, analyte-filtered, covariate-adjusted, and winsorized SomaScan (log2 RFU) and Olink (NPX) protein abundance matrices were used for downstream association analyses.

### Antemortem CSF proteomics differentiates pathology-confirmed FTLD-TDP, FTLD-tau, and AD

Differential abundance analyses were first performed in the SomaScan dataset across autopsy cohort molecular classes of FTLD-TDP, FTLD-tau, AD, and AD biomarker-negative CTL (**Supplementary Table 7**). A total of 776 aptamers reached FDR-significance in FTLD-TDP vs. CTL (1,141 nominal; **Fig. 2a**), 648 in FTLD-tau vs. CTL (1,056 nominal; **Fig. 2b**), and 28 in FTLD-TDP vs. FTLD-tau (346 nominal; **Fig. 2c**). Targets elevated in FTLD-TDP relative to FTLD-tau included markers of post-synaptic scaffolding (HOMER1, SHANK3, DLG4), mitochondrial function (ATP5PF, COX5A), gene expression regulation (KHSRP, TCEAL5, NUDT21), and neurofilaments (NEFL, NEFH), whereas the most elevated markers in FTLD-tau vs. FTLD-TDP included extracellular matrix remodeling protein MMP10 and synaptic calcium signaling protein calcineurin (PPP3R1). Of the 28 FDR-significant aptamers that differentiated FTLD-TDP vs. FTLD-tau, HOMER1 and ATP5PF exhibited the strongest classification performance in receiver operating characteristic (ROC) analyses with area under the curve (AUC) values of 0.89 (HOMER1) and 0.88 (ATP5PF; **Fig. 2d-f; Supplementary Table 8**).

**Figure 2.**
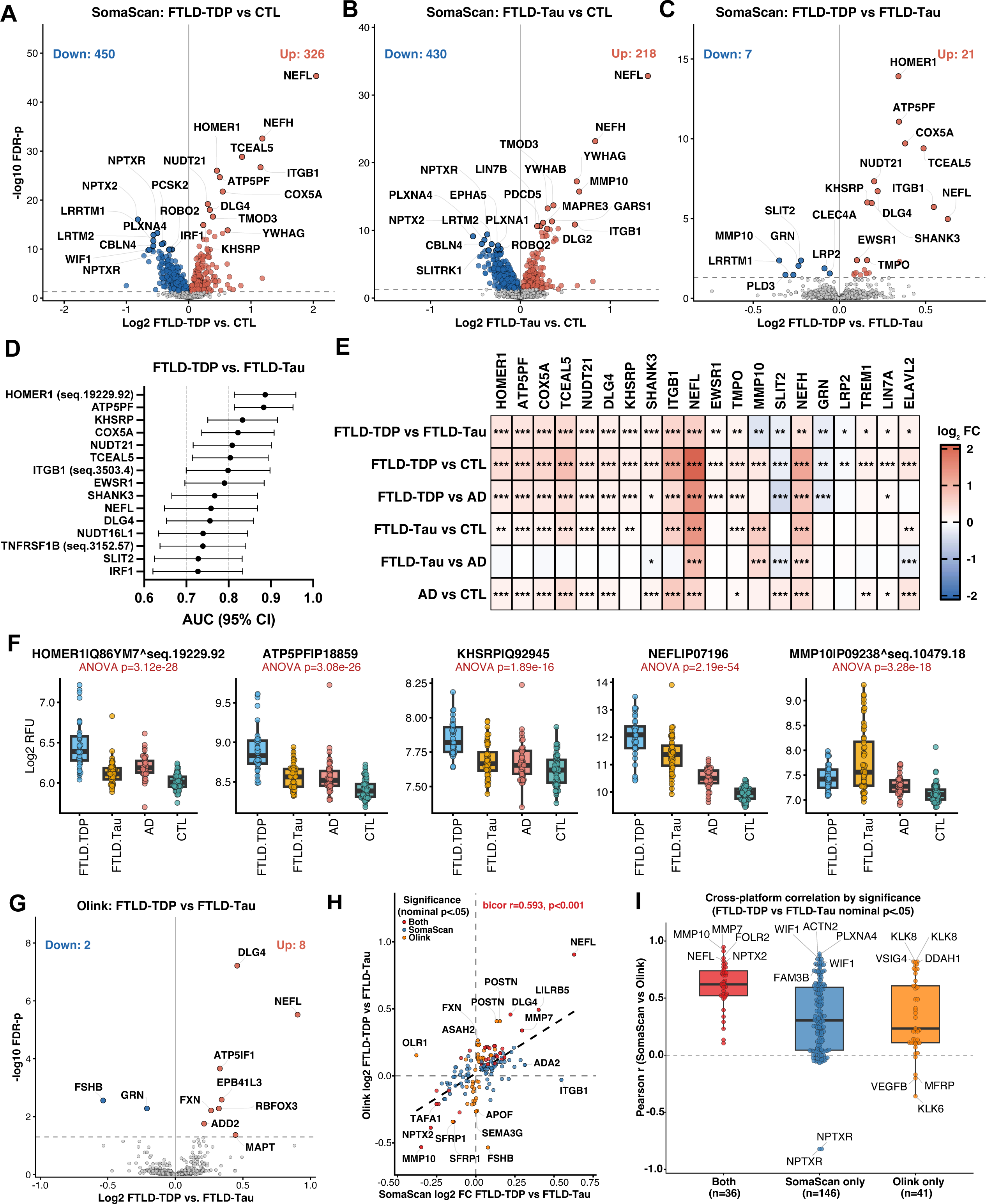
Multi-platform CSF proteomics identify and validate differentiators of FTLD-TDP and FTLD-tau. **a–c**, Volcano plots of SomaScan differential protein abundance for **a,** FTLD-TDP vs. CTL, **b,** FTLD-tau vs. CTL, and **c,** FTLD-TDP vs. FTLD-tau. Dashed line, FDR-P = 0.05; counts denote aptamers reaching FDR significance in each direction (up=red, down=blue). **d,** Area under the receiver operating characteristic curve (AUC, with 95% CI) for the FDR-significant aptamers from **c**, ranked by discriminative performance for FTLD-TDP vs. FTLD-tau. **e,** Heat map of log2 fold change for the top discriminating aptamers from **c** across six pairwise SomaScan molecular-class contrasts. *P < 0.05, **P < 0.01, ***P < 0.001 (FDR-corrected). **f,** Aptamer abundance (log2 relative fluorescence units, RFU) by molecular class for five top discriminating proteins. P values, one-way ANOVA. **g,** Volcano plot of Olink differential protein abundance for FTLD-TDP vs. FTLD-tau. Dashed line, FDR-P = 0.05. **h,** Cross-platform comparison of log2 fold change (FTLD-TDP vs. FTLD-tau; biweight midcorrelation [bicor]) for matched SomaScan–Olink analyte pairs with nominal P < 0.05 on either platform, colored by platform significance. **i,** Per-analyte Pearson correlation between SomaScan and Olink abundance for matched pairs, grouped by significance category from **h**. Box plots (**d**, **f**, **i**) show median, 25th–75th percentiles (box) and range (whiskers), with individual data points overlaid.

FTLD molecular class differential abundance contrasts were conducted in the Olink dataset to validate SomaScan findings and identify additional hits not measured on the SomaScan platform (**Supplementary Table 9**). The Olink dataset included 115 autopsy cases (34 AD, 48 FTLD-tau, 33 FTLD-TDP) and 55 CTL from the SomaScan dataset as well as 33 new FTLD-tau and 10 new FTLD-TDP cases. Ten Olink proteins reached FDR-significance in FTLD-TDP vs. FTLD-tau comparisons (101 nominal; **Fig. 2g**). Top hits included elevated DLG4, NEFL, and ATP5IF1 in FTLD-TDP, which replicated SomaScan results with somewhat attenuated classification performance (AUC range 0.75 – 0.78; **Supplementary Table 10**).

Comparison of FTLD-TDP vs. FTLD-tau log-fold changes for hits with nominal significance on either platform indicated strong cross-platform concordance in effect sizes (bicor=0.54) with NEFL, DLG4, and MMP10 among the largest log-fold changes across platforms (**Fig. 2h)**. Median cross-platform correlations for matching SomaScan aptamer and Olink protein pairs (**Fig. 2i**) were highest for targets with nominal significance on both platforms (median *r*=0.62) compared to SomaScan (median *r*=0.30) or Olink only (median *r*=0.28). Among proteins with significance on both platforms, MMP10 and NEFL exhibited strong cross-platform correlations (MMP10: *r*=0.92; NEFL: *r*=0.86).

We further validated FTLD molecular class-related protein alterations across and within platforms by correlating effect sizes with those reported in a recent Olink proteomics study by Hok-A-Hin et al.^5^ (n=666 proteins). We observed good effect size concordance of differentially abundant proteins for FTLD-TDP and FTLD-tau vs. CTL contrasts across and within platforms (bicor range: 0.48 to 0.72; **Supplementary Fig. 2**). MMP10 emerged as a consistent hit for FTLD-TDP vs. FTLD-tau contrasts across cohorts. However, there was poor concordance for the broader collection of differentially abundant FTLD-TDP vs. FTLD-tau proteins vs. Hok-A-Hin et al. (bicor range: 0.05 to 0.08), which may reflect the lower depth of coverage, including absence of key hits (e.g., HOMER1, DLG4), and inclusion of clinically-defined cases in FTLD-TDP (16% svPPA) and FTLD-tau groupings (62% PSP-RS or CBS) in the Hok-A-Hin et al. dataset.

### AD CSF proteomic signatures are detectable in FTLD with AD co-pathology yet distinct from top FTLD proteomic markers

AD vs. CTL SomaScan comparisons indicated robust elevations in markers previously linked to AD, including SMOC1, YWHAG, and CNRIP1 (**Extended Data Fig. 1a**). NEFL exhibited the largest elevations in FTLD-TDP vs. AD and FTLD-tau vs. AD contrasts (**Extended Data Fig. 1b-c).** The majority of FDR-significant aptamers that differed between FTLD-TDP and FTLD-tau also differed between these groups and AD, including higher HOMER1 in FTLD-TDP and higher MMP10 in FTLD-tau (**Fig. 2e-f**). Similar results were observed in Olink comparisons (**Extended Data Fig. 1d-f)**, with NEFL again exhibiting the largest elevations in FTLD-TDP and FTLD-tau vs. AD.

To determine the contribution of AD co-pathology to proteomic profiles in FTLD, we examined differential CSF SomaScan protein abundance in all FTLD-spectrum cases with CSF pTau181/Aβ42 ratios (n=18 positive vs. n=88 negative) as well as autopsy-confirmed FTLD with ADNC staging (n=17 high/intermediate ADNC vs. n=40 low/none ADNC), adjusting for *APOE* ε4 positivity (**Extended Data Fig. 2a**). The CSF SomaScan proteomic signature of pTau181/Aβ42 positivity within FTLD exhibited concordance with primary AD pathology vs. CTL effects, including elevations in PEA15, SMOC1 and YWHAG (**Extended Data Fig. 2b-c**). In contrast, core markers of neurodegeneration including NPTX2 and NEFL exhibited small, non-significant associations with AD co-pathology despite differential abundance in primary AD vs. CTL, reinforcing FTLD as the principal source of variance in these proteins when FTLD is the primary etiology. SomaScan pTau181/Aβ42 positivity proteomic effect sizes in FTLD were concordant with Olink CSF pTau181/Aβ42 proteomic effect sizes in FTLD (**Extended Data Fig. 2d**) and SomaScan ADNC-defined (high/intermediate vs. low/none) proteomic effect sizes in autopsy-confirmed FTLD (**Extended Data Fig. 2e**). SMOC1 levels in autopsy-confirmed FTLD cases with high ADNC approached levels observed in primary AD, whereas top FTLD-TDP vs. FTLD-tau hits MMP10 and HOMER1 did not track with ADNC stage in FTLD (**Extended Data Fig. 2f**), confirming specificity to FTLD rather than AD co-pathology.

### FTLD pathological subtype screening reveals subtype-specific diagnostic CSF biomarker candidates

Although TDP and tau represent major molecular classes of FTLD, there is clinically relevant pathological heterogeneity between subtypes within each class. We therefore performed differential abundance analysis across FTLD-TDP and FTLD-tau subtypes (**Supplementary Tables 12-13**) and one-vs-rest ROC curves to evaluate diagnostic performance of candidate proteins in discriminating a given FTLD subtype vs. other diseases combined (i.e. other FTLD subtypes and AD) and vs. CTL (**Supplementary Tables 14-15**). A heatmap of normalized abundance values for top performing aptamers (AUCs>0.80) are plotted across individuals and subtypes in **Fig. 3a**. Elevated MMP10 separated PiD from all other cases with near-perfect accuracy (AUC=0.98, 100% specificity, 93% sensitivity, **Fig. 3b-d**), indicating that prior signals in the broader FTLD-tau molecular class were driven predominantly by PiD. Top aptamers for other sporadic pathologies included elevated levels of calcium/calmodulin-regulated neuronal phosphodiesterase PDE1B in CBD (AUC=0.83, 73% specificity, 85% sensitivity), interferon-stimulated ubiquitin-like modifier ISG15 in TDP-C (AUC=0.87, 89% specificity, 77% sensitivity), and neural proliferation differentiation and control protein 1 (NPDC1) in TDPb (AUC=0.83, 78% sensitivity, 88% specificity). Consistent with prior work in genetic FTLD, *GRN*-predominant TDP-A cases exhibited highly sensitive and specific depletions of GRN and C1QTNF1 protein levels^6^. DLG4 and HOMER1 were notably elevated in each individual FTLD-TDP subtype relative to other subtypes, suggesting broader generalizability to FTLD-TDP as a molecular class than any given subtype.

**Figure 3.**
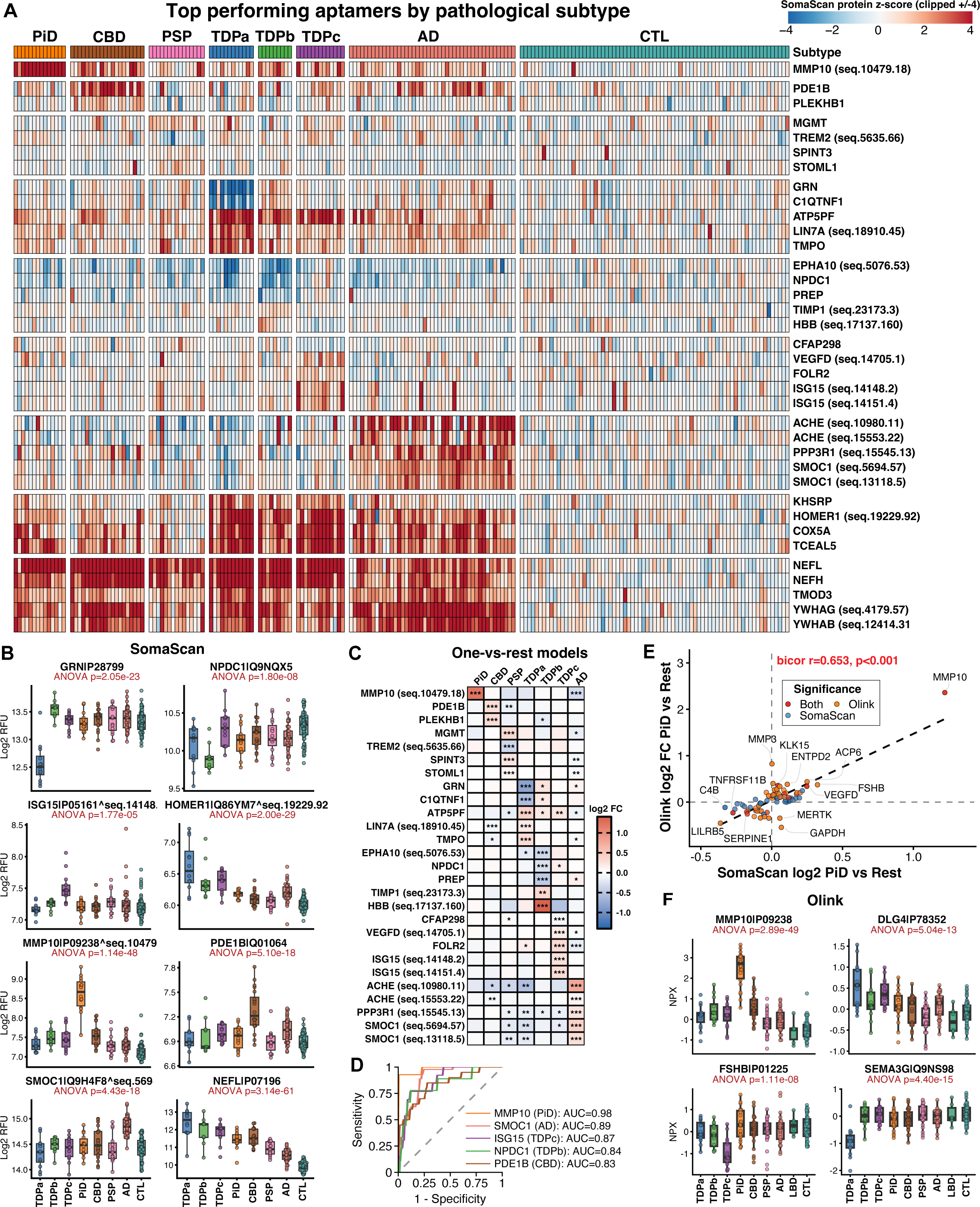
Pathological subtype–specific CSF biomarker candidates in FTLD. **a**, Heat map of SomaScan aptamer abundance (z-scored, clipped ±4) for aptamers achieving one-vs-rest AUC > 0.80 for at least one pathological subtype, plotted across individuals grouped by subtype (PiD, CBD, PSP, TDPa, TDPb, TDPc, AD, CTL). **b,** Aptamer abundance (log2 RFU) by subtype for top aptamers from **a**. P values, one-way ANOVA. **c,** Heat map of log2 fold change for one-vs-rest differential abundance models (each subtype vs. all other diseases combined) for the aptamer set in **a**. *P < 0.05, **P < 0.01, ***P < 0.001 (FDR-corrected). **d,** Receiver operating characteristic curves for the top one-vs-rest classifier proteins, with AUC. **e,** Cross-platform comparison of SomaScan and Olink log2 fold change for PiD vs. rest (biweight midcorrelation), colored by platform significance. **f,** Olink abundance (NPX) by subtype for four representative proteins, including LBD as an additional Olink-only comparison group. P values, one-way ANOVA. Box plots (**b**, **f**) show median, 25th–75th percentiles (box) and 1.5× interquartile range (whiskers), with individual data points overlaid.

Olink analyses, which included the 14 original PiD cases from the SomaScan dataset and 8 new PiD cases, validated elevated MMP10 as highly sensitive and specific to PiD (AUC=0.99, 91% sensitivity, 98% specificity; **Fig. 3e-f**). Olink MMP3 was also elevated in PiD vs. rest (AUC=0.82), although MMP3 cross-platform correlation was poor (*r*=-0.06) and SomaScan MMP3 did not show a PiD signal. Despite similar molecular class performance, Olink HOMER1 exhibited a distinct subtype pattern from SomaScan HOMER1 with significant elevations in TDPa, but not TDPb or TDPc, and decreased abundance in PiD and PSP, but not CBD. In addition to MMP10 and MMP3, the only other Olink protein to surpass a one-vs-rest AUC>0.80 for classifying a sporadic FTLD subtype was follicle stimulating hormone subunit beta (FSHB), which was decreased in TDPc cases vs. rest. Alongside progranulin protein (“GRN”), axon guidance protein semaphorin 3G (SEMA3G) exhibited marked decreases in TDPa cases, consistent with a recent report of decreased SEMA3G in *GRN* mutation carriers^11^. DOPA decarboxylase (DDC), recently proposed as a diagnostic biomarker candidate for LBD^12^, was elevated across LBD, PSP, and CBD but did not differ across these groups (**Supplementary Table 13**), supporting the hypothesis that elevated DDC is sensitive to dopaminergic dysfunction across parkinsonian disorder-causing neuropathologies but not adequate as a stand-alone biomarker for 4R-tauopathy vs. α-synuclein differential diagnosis^13^.

Olink HOMER1 failed QC due to low detectability and was not included in primary molecular class or pathology subtype models. However, we examined its distribution across cases given the large signal observed on SomaScan. Notably, Olink HOMER1 also exhibited strong classification performance with higher abundance in FTLD-TDP vs. FTLD-tau (AUC=0.75). However, this difference was driven by lower levels in FTLD-tau vs. CTL rather than higher levels in FTLD-TDP vs. CTL (**Supplementary Fig. 3a**). Pathological subtype contrasts showed lower Olink HOMER1 in both PiD and PSP compared to other groups (**Supplementary Fig. 3b)**. HOMER1 cross-platform concordance was also modest (*r*=0.31; **Supplementary Fig. 3c**), raising the possibility that the SomaScan HOMER1 aptamer (seqID: 19229.92) may capture an alternative and more reliably detectable proteoform than the Olink HOMER1 monoclonal antibody with greater sensitivity to FTLD-TDP.

### Protein co-expression network analysis identifies RNA metabolism, post-translational modification, immune, and synaptic pathway alterations in FTLD

To identify large-scale proteomic pathways linked to FTLD subtypes and AD, we constructed a harmonized, multi-platform protein co-expression network among samples assayed on both SomaScan and Olink platforms. Inspection of multidimensional scaling (MDS) plots confirmed removal of the platform effect and expected clustering of matched analyte pairs with high cross-platform concordance following harmonization (**Supplementary Fig. 4**). Network analysis revealed 60 protein co-expression modules (**Supplementary Table 16**), which were functionally annotated based on Gene Ontology (GO), pathway and cell type enrichment analyses (**Supplementary Table 17; Supplementary Fig. 5**). Module eigenproteins were then examined in relation to pathological subtype and disease severity outcomes (restricted to FTLD spectrum cases), including CDR®+NACC-FTLD sum of boxes (clinical severity), frontotemporal gray matter volume, and CSF NfL elecsys (**Fig. 4a; Supplementary Table 18**). Correlations of individual proteins with FTLD disease severity outcomes are also reported for SomaScan (**Supplementary Table 19)** and Olink (**Supplementary Table 20)**.

**Figure 4.**
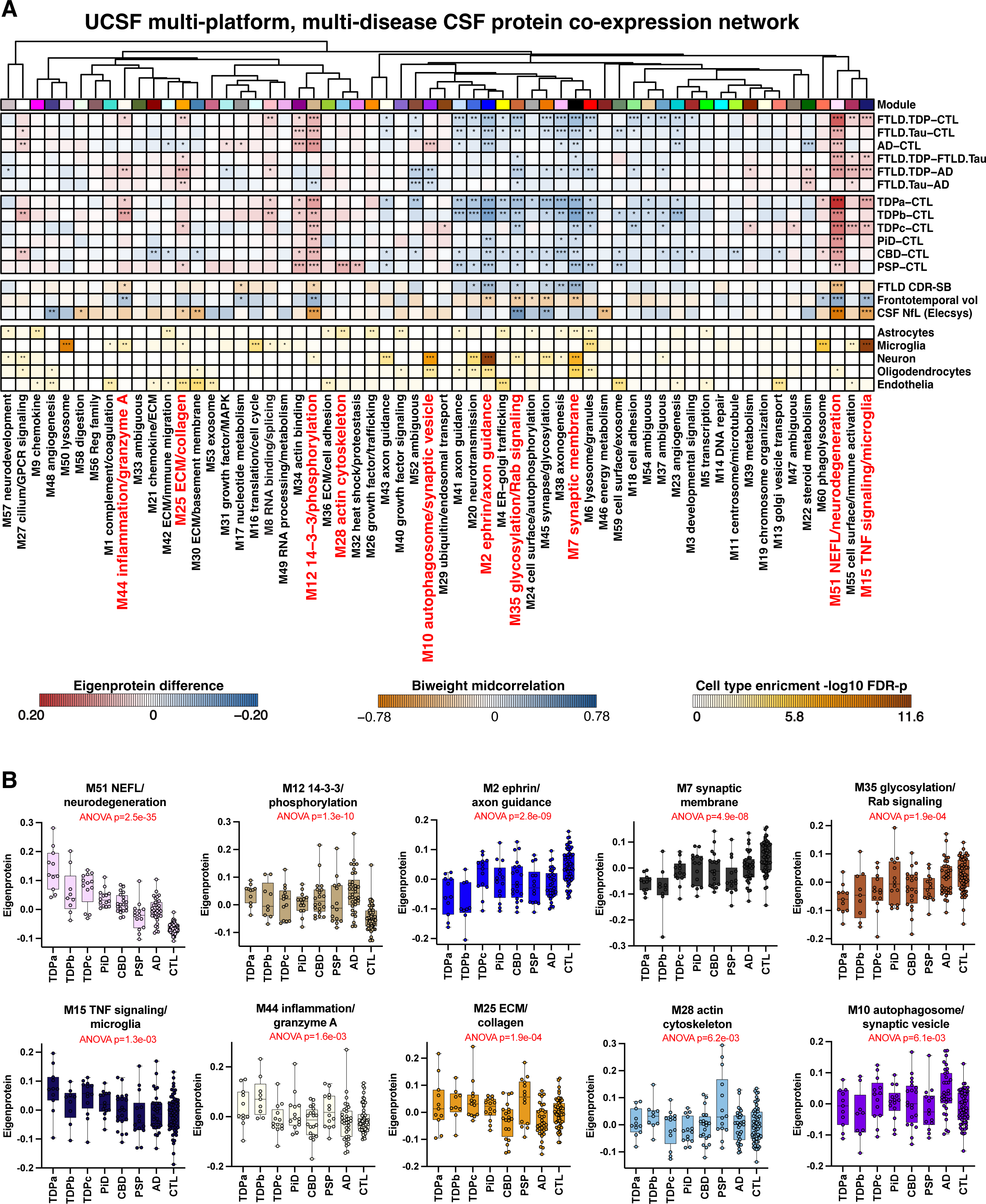
Multi-platform CSF protein co-expression network reveals pathway alterations across FTLD and AD. **a**, Relationship between 60 weighted co-expression network modules assembled from n = 5,006 harmonized SomaScan/Olink analytes across n = 216 samples assayed on both platforms. Modules are grouped by hierarchical clustering and functionally annotated. Heatmaps depict pairwise molecular class and pathological subtype contrasts (eigenprotein difference), biweight midcorrelations with disease severity outcomes (CDR®+NACC-FTLD sum of boxes, frontotemporal gray matter volume, and CSF NfL [Elecsys]) and cell-type marker enrichment (one-tailed Fisher’s exact test) for astrocyte, microglia, neuron, oligodendrocyte, and endothelial marker gene sets. *P < 0.05, **P < 0.01, ***P < 0.001 (nominal). **b,** Module eigenprotein values by pathological subtype for ten representative modules. P values, one-way ANOVA. Box plots (**b**) show median, 25th–75th percentiles (box) and range (whiskers), with individual data points overlaid.

Several modules exhibited alterations across multiple molecular classes (**Fig. 4b**). M51 NEFL/neurodegeneration (hubs: NEFL, TCEAL5, ITGB1, NEFH) was robustly elevated in all diseases vs. CTL. Across all FTLD cases, M51 exhibited the strongest positive correlation with clinical severity (bicor: *r*=0.38, *p*=4.2e-05) and negative correlation with frontotemporal volume (bicor: *r*=-0.37, *p*=9.8e-05). M51 also included top differentiators of FTLD-TDP vs. FTLD-tau (e.g., HOMER1, DLG4, ATP5IF1, ATP5PF). SomaScan and Olink MMP10 were also members in M51 but were not the strongest drivers of overall module abundance (kME <0.69). Similar to M51, M12 14-3-3/phosphorylation (hubs: 14-3-3 family proteins, PDCD5, PTPA) was strongly elevated across multiple diseases and harbored markers of neurodegeneration and post-translational modification pathways. A large cluster of modules related to synaptic signaling, axon guidance, and neurodevelopment exhibited robust decreases across multiple diseases, with M2 ephrin/axon guidance (hubs: NRG3, CADM2, EPHB6, NRXN1) and M7 synaptic membrane (hubs: ADGRB3, PCDH9, PLXNA4) exhibiting some of the strongest negative correlations with clinical severity and positive correlations with frontotemporal volume. These modules also harbored NPTX2 (SomaScan NPTX2: M2 kME=0.83; Olink NPTX2: M7 kME=0.70), which was the strongest individual protein correlate of clinical severity across platforms (**Supplementary Tables 19-20**).

Several modules were selectively elevated within a molecular class or subtype. M8 RNA binding (hubs: KARS1, HNRNPF, HNRNPA2B), M44 inflammation/granzyme A (hubs: GZMA, CD48, FASLG), and M15 TNF signaling/microglia (TNFRSF1B, TNFRSF1A, TNFSF8) were predominantly elevated in FTLD-TDP subtypes. Two related modules linked to cytoskeletal and heat shock pathways (M28 actin cytoskeleton [hubs: HARS1, YAP1, STAT3]; M54 heat shock/proteostasis [hubs: HSPA1A, HSPA1B, DNASE1L2]) were predominantly elevated in PSP. M10 autophagosome/synaptic vesicle (hubs: YWHAG, GAP43, UBE2V2) was strikingly elevated in AD but not FTLD and included SMOC1 alongside other proteins associated with AD co-pathology within FTLD.

Overall, our multi-disease, multi-platform proteomic network reveals synaptic and post-translational modification signatures common across pathologies, RNA metabolism and immune signatures disproportionately altered in FTLD-TDP, cytoskeletal and heat shock signatures disproportionately altered in PSP, and synaptic vesicle signaling and autophagy signatures disproportionately altered in AD.

### Targeted immunoassay validation confirms PiD-specific elevations of CSF MMP10

MMP10 exhibited the strongest subtype-specific diagnostic performance in our large-scale proteomics screen. We therefore performed a series of follow-up analyses to validate diagnostic specificity to PiD. We first measured CSF MMP10 on a high-sensitivity Meso Scale Discovery (MSD) immunoassay, allowing for absolute quantification of MMP10 concentration on a targeted platform. MSD MMP10 was quantified in the same 344 samples from the UCSF Olink cohort (286 unique subjects [including 22 PiD]) and an independent validation cohort of 82 samples (including 10 PiD) from the Penn FTD Center. MSD MMP10 exhibited high intra-assay precision in both cohorts (UCSF median coefficient of variation [CV]=3.9%, Penn median CV=2.7%) with only two samples with CV>20% that were excluded from analysis (**Fig. 5a**). MSD MMP10 exhibited large cross-platform correlations with Olink (*r*=0.98) and SomaScan (*r*=0.91; **Fig. 5b-c**). In the UCSF autopsy cohort, baseline MMP10 was markedly elevated in PiD (median 1,395 pg/mL) relative to non-PiD pathologies (median 230 pg/mL) and CTL (median 143 pg/mL; **Supplementary Table 21; Fig. 5d**), separating PiD from non-PiD cases with high accuracy (AUC=0.97, sensitivity 91%, specificity 98%). MSD MMP10 exhibited positive correlations with CSF NfL (Elecsys) across PiD (*r*=0.65), non-PiD (*r*=0.56), and CTL (*r*=0.35; **Fig. 5e**), but the vast majority of non-PiD cases with the highest NfL concentrations did not reach MMP10 concentrations at the lower end of the PiD range.

**Figure 5.**
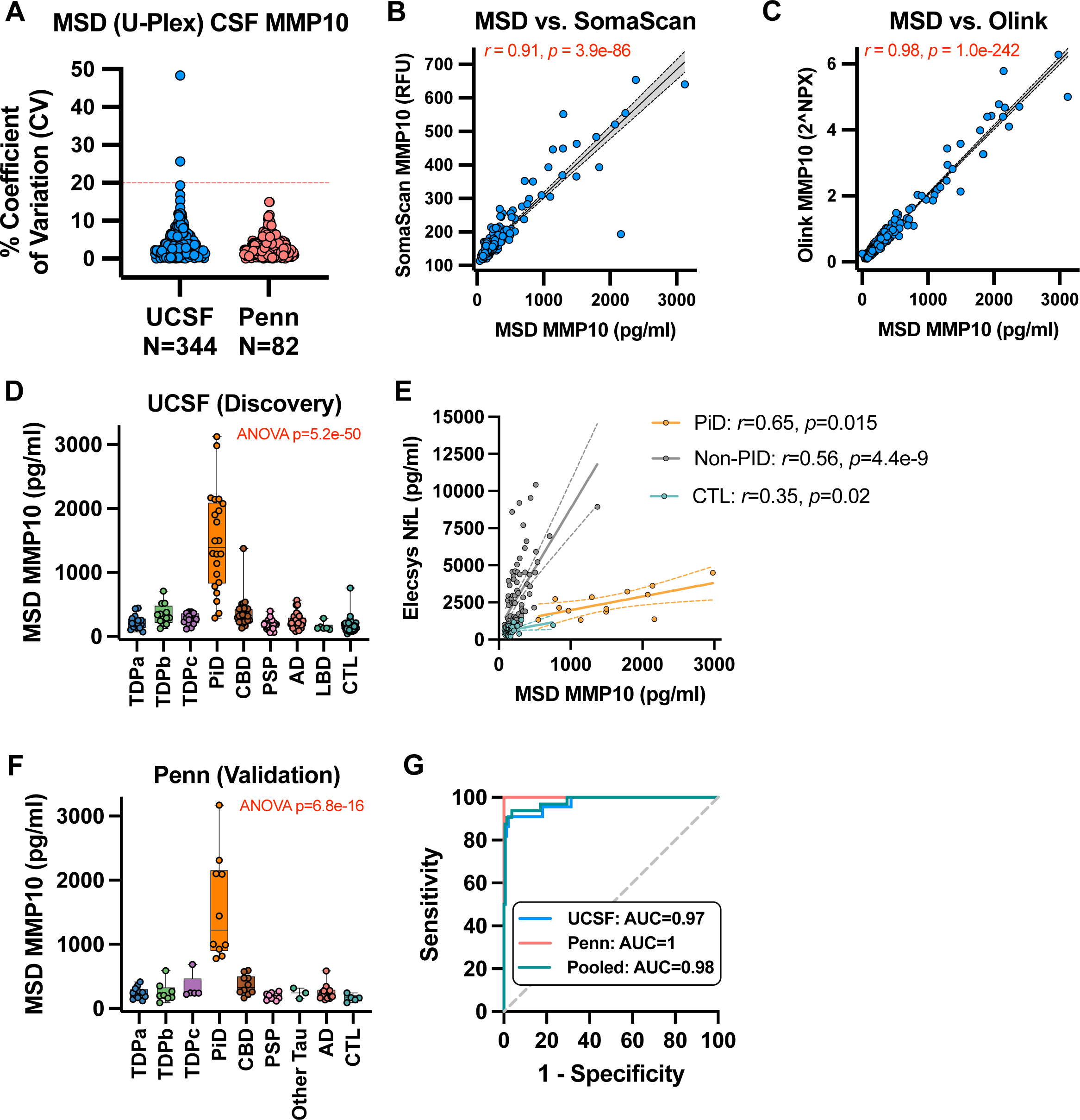
Targeted immunoassay validation supports CSF MMP10 as a diagnostic biomarker for PiD. **a**, Intra-assay precision (% coefficient of variation) for the MSD U-Plex CSF MMP10 assay in the UCSF (n = 344 samples) and Penn (n = 82 samples) cohorts. **b, c,** Pearson correlation of MSD (Meso Scale Discovery) CSF MMP10 concentration with **b,** SomaScan MMP10 (RFU) and **c,** Olink MMP10 (2^NPX). **d,** MSD MMP10 concentration by pathological subtype in the UCSF discovery cohort. **e,** Correlation of MSD MMP10 with CSF neurofilament light chain (Elecsys immunoassay), stratified by CTL, PiD, and non-PiD. **f,** MSD MMP10 concentration by pathological subtype in the Penn validation cohort. **g,** Receiver operating characteristic curves for MSD MMP10 classification of PiD vs. non-PiD cases in the UCSF, Penn, and pooled cohorts, with AUC. Box plots (**d, f**) show median, 25th–75th percentiles (box) and range (whiskers), with individual data points overlaid.

Penn validation analyses revealed highly similar median MMP10 concentrations to those observed in UCSF (PiD: 1,223 pg/mL, non-PiD pathologies: 239 pg/mL, CTL: 164 pg/mL; **Fig. 5f**), with a remarkable non-overlapping PiD vs. rest distribution resulting in perfect classification accuracy (AUC=1, sensitivity 100%, specificity 100%). In a pooled cohort analysis (**Fig. 5g)**, an MSD MMP10 cut-off >549 pg/mL accurately classified 30 out of 32 PiD cases and misclassified only 8 out of 216 non-PiD pathologies (AUC=0.98, sensitivity 94%, specificity 96%). These results support elevated CSF MMP10 as a robust diagnostic biomarker for PiD, independent of neurodegeneration, that can be reliably measured with absolute quantification on a targeted assay platform suitable for clinical translation.

#### CSF MMP10 is elevated in a subset of 4R-tau-predominant *MAPT* mutation carriers

MMP10 elevations in PiD raised the possibility of specificity to 3R-tau. We and others have previously reported gene group-level elevations in CSF MMP10 in familial FTLD-tau (*MAPT*) vs. familial FTLD-TDP (*GRN*, *C9orf72*)^6,11^. To determine how CSF MMP10 differed across individuals carrying *MAPT* mutations with varying predilections for 3R-and 4R-tau predominant aggregation, we examined CSF SomaScan MMP10 in the subset of UCSF cases with familial FTLD and a larger set of familial FTLD cases from the ALLFTD Consortium. In UCSF, MMP10 levels were above the cut-off for classifying PiD in 2 out of 11 *MAPT* carriers (both symptomatic) but below the cut-off in all 53 familial FTLD-TDP cases (29 *C9orf72*, 21 *GRN*, 3 *TARDBP*; **Extended Data Fig. 3a**). MMP10 was also elevated in the only individual carrying a TSC1 mutation, a known cause of tauopathy^14^. In ALLFTD, MMP10 levels were above the PiD cut-off in 4 of 54 *MAPT* carriers, including 3 symptomatic cases and 1 presymptomatic case but below the cut-off in all 112 familial FTLD-TDP cases (78 *C9orf72*, 34 *GRN*; **Extended Data Fig. 3b**). Of the 6 *MAPT* carriers with elevated MMP10 across cohorts, 5 had P301L mutations (out of 15 P301L cases) and one had an S305I mutation (out of 3 S305I cases; **Extended Data Fig. 3c-d**). Given that P301L and S305I mutations give rise to predominantly 4R-tau aggregates^15,16^, elevations in CSF MMP10 are unlikely to be exclusive to 3R-tau aggregation and may instead be sensitive to other pathological features that are shared across PiD and a subset of familial tauopathy cases.

### CSF MMP10 tracks with clinico-anatomical profiles suggestive of underlying PiD in clinically-diagnosed bvFTD and PPA

We next examined CSF MMP10 levels in the cohort of sporadic clinically-diagnosed FTLD cases with SomaScan data. Ten of 69 sporadic FTLD clinical cases exhibited CSF SomaScan MMP10 levels above the cut-off for classifying PiD, whereas none of the in-vivo biomarker-confirmed AD or CTL cases exhibited elevated CSF MMP10 (**Fig. 6a**). MMP10+ FTLD cases were exclusively comprised of syndromes with known associations to PiD, specifically 4 bvFTD, 5 svPPA, and one PPA unspecified case (**Fig. 6b**)^17^. To increase confidence that PiD was the most likely underlying etiology for clinically-diagnosed SomaScan MMP10+ FTLD cases, we examined the prevalence of clinician-predicted pathological diagnoses of PiD, determined via multidisciplinary case reviews of clinical and neuroimaging data at the time of study participation. PiD was documented as the primary predicted pathology for 7 of 10 MMP10+ cases (3 of 4 bvFTD, 3 of 5 svPPA, 1 of 1 PPA unspecified; **Fig. 6c**) and second most likely pathology for two of the three remaining MMP10+ cases (**Supplementary Table 22**). Knife-edge atrophy patterns, common in PiD^18^, were noted for the majority of MMP10+ cases. In contrast, PiD was not the primary predicted pathology for any of the 59 SomaScan MMP10-cases.

**Figure 6.**
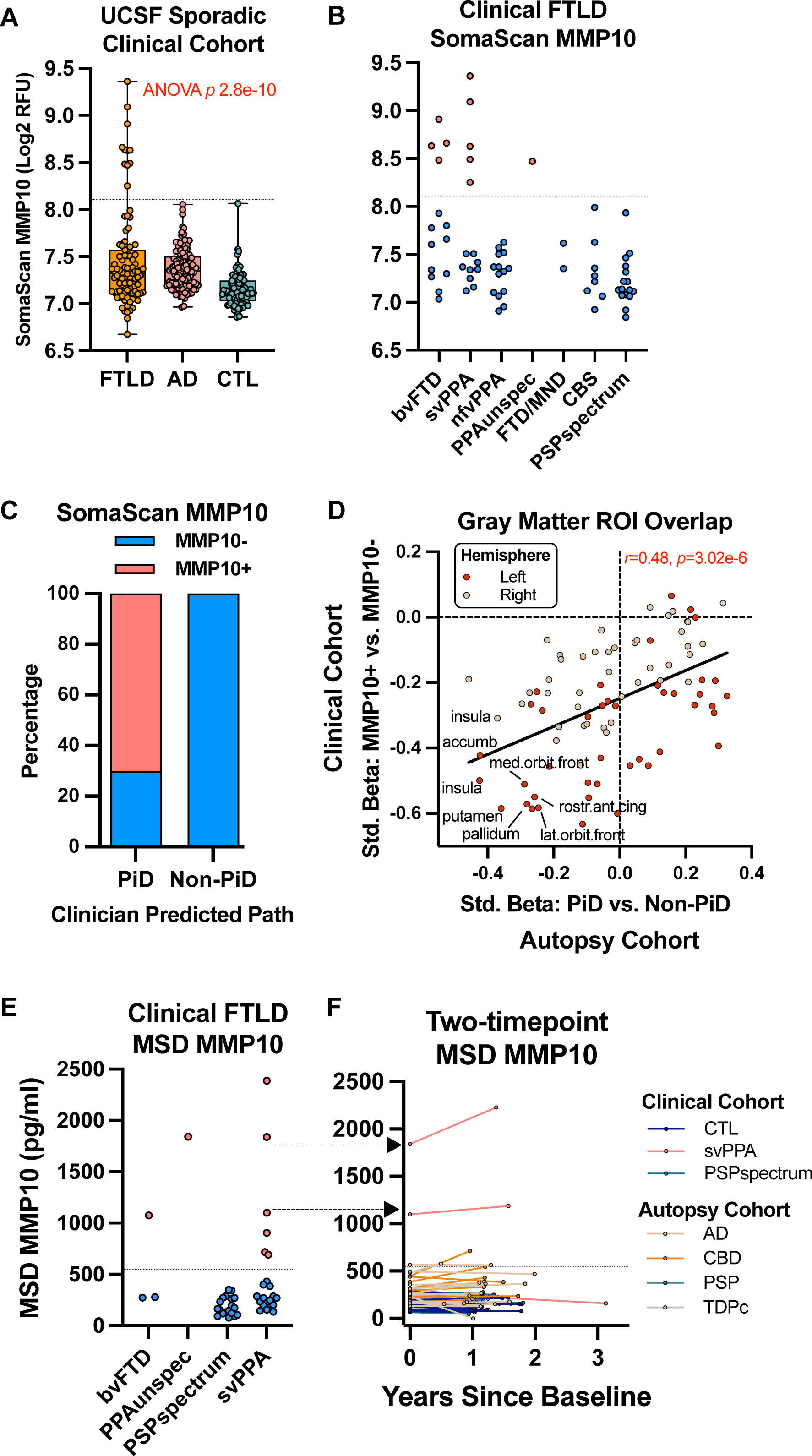
CSF MMP10 tracks with clinico-anatomical profiles suggestive of underlying PiD in clinically-diagnosed FTLD. **a**, SomaScan MMP10 abundance in the UCSF sporadic clinical cohort by diagnostic group (FTLD, AD, CTL). Dashed line, PiD classification cut-off (from Fig. 3d). P value, one-way ANOVA. **b,** SomaScan MMP10 abundance among clinically-diagnosed FTLD cases by clinical syndrome (bvFTD, svPPA, nfvPPA, PPA unspecified, FTD/MND, CBS, PSP-spectrum). **c,** Percentage of SomaScan MMP10+ and MMP10− clinical FTLD cases with clinician-predicted primary pathology of PiD versus non-PiD, based on multidisciplinary case review of clinical and neuroimaging data. **d,** Pearson correlation between standardized regional gray matter atrophy effect sizes (107 Desikan-Killiany ROIs, models covaried for clinical phenotype, age, sex, and intracranial volume) for PiD vs. non-PiD in the autopsy cohort and MMP10+ vs. MMP10− in the clinical cohort (bvFTD/PPA-spectrum only), colored by cohort and hemisphere. **e,** MSD MMP10 concentration in the clinical FTLD cohort by clinical syndrome. Dashed line, PiD classification cut-off (549 pg/mL). **f,** Longitudinal MSD MMP10 concentration across two timepoints, colored by diagnosis. Box plots (**a, b, e**) show median, 25th–75th percentiles (box) and range (whiskers), with individual data points overlaid.

In addition to clinical record review, we quantified the overlap in brain volumetric atrophy patterns across 107 gray matter ROIs from the Desikan-Killiany atlas for autopsy-confirmed PiD (vs. non-PiD subtypes) and MMP10+ clinical cases without autopsy (vs. MMP10-cases). Analyses were restricted to bvFTD or PPA-spectrum FTLD phenotypes (i.e., nfvPPA, svPPA, PPA unspecified). Models covaried for clinical phenotype, age, sex, and intracranial volume. Standardized effect sizes showed a robust positive correlation between the two contrasts (*r*=0.48, p=3.02e-08; **Fig. 6d; Supplementary Table 23**), indicating that MMP10-related atrophy in-vivo aligns with the anatomic pattern observed in autopsy-confirmed PiD. Ten ROIs exhibited significant atrophy effects in both autopsy-confirmed PiD and clinically-diagnosed CSF MMP10+ groups, encompassing frontoinsular, orbitofrontal, temporal, and subcortical structures. With the exception of the insula, which exhibited bilateral atrophy across groups, the atrophied ROIs were left-hemisphere regions, likely reflecting the enrichment of language-predominant phenotypes in the analytic sample.

Of the 45 clinically-diagnosed FTLD cases with MSD MMP10 data, 8 exhibited baseline MMP10 levels above the 549 pg/mL cut-off for PiD, including 1 bvFTD, 6 svPPA, and 1 PPA unspecified (**Fig. 6e**). None of the 19 PSP-spectrum clinical cases exhibited elevated MSD MMP10. MSD MMP10 was also measured at an additional timepoint in 14 CTL and 43 individuals with clinical (svPPA, PSP-spectrum) or autopsy-confirmed diagnoses (AD, CBD, PSP, TDPc), including 2 svPPA cases with elevated baseline MMP10. Notably, the MMP10+ svPPA case with the highest baseline MMP10 concentration also exhibited the largest annualized increase in MMP10 at follow-up (280 pg/mL per year; **Fig. 6f**). Only two of the 54 MMP10-cases at baseline had levels that were slightly above the cut-off at follow-up (1 CBD, 1 AD), but were still appreciably lower than baseline concentrations in the MMP10+ svPPA cases.

Collectively these cross-sectional and longitudinal findings in clinically-diagnosed FTLD indicate that elevated CSF MMP10 identifies cases with clinical and neuroanatomical features consistent with underlying PiD across behavioral and language-predominant phenotypes, supporting broad clinical relevance.

## DISCUSSION

Our data demonstrate that large-scale CSF proteomics can identify molecular class and subtype-specific diagnostic biomarker candidates and proteomic signatures across the FTLD clinico-pathological continuum. Most notably, we identified and validated CSF MMP10 as having high potential to be a robust diagnostic biomarker for PiD across three independent methods of protein quantification and two independent autopsy cohorts. In parallel, network analysis revealed coordinated changes in RNA metabolism, immune signaling, proteostasis, and synaptic pathways across FTLD and AD, underscoring both shared and disease-enriched biology.

CSF MMP10 (also known as stromelysin-2) was notable for its strength and consistency of validation in PiD, emerging across SomaScan and Olink proteomic platforms and a targeted MSD assay that was replicated in an independent autopsy cohort. In clinically-diagnosed FTLD, MMP10 elevations only occurred among syndromes with known ties to PiD and also tracked with brain atrophy patterns seen in PiD, supporting its face validity in living patients. Although MMP10 is mildly elevated in other FTLD subtypes and AD cases compared to controls^19^, the PiD signal was up to 6-fold higher than non-PiD signal, preserving strong subtype-specific diagnostic value. The near-perfect classification performance (AUC 0.97-1.00) for PiD places MMP10 among the strongest reported CSF biomarker signals for any autopsy-confirmed neurodegenerative etiology to date, rivaling CSF pTau and Aβ isoforms in AD. Clinically, elevated CSF MMP10 could rule-in PiD for tau-targeted therapies in bvFTD, help rule out PiD for TDPc-targeted therapies in svPPA (e.g., NCT05184569^20^), and could help stratify PiD vs. 4R-tau etiologies in tau-targeted trials in nfvPPA^21^.

Mechanistically, the link between MMP10, tau, and neuroinflammation is unclear. Prior work has implicated MMP10 in inflammatory cell migration and glial activation. For example, apoptosis-associated speck-like protein containing a C-terminal caspase recruitment domain (ASC) specks are elevated in PiD and can induce microglial release of MMP10 and MMP3^22^, a closely-related stromelysin (stromelysin-1) that was also elevated in PiD on the Olink platform. MMP10 and MMP3 can also be secreted by senescent astrocytes^23^, resulting in a cytotoxic pro-inflammatory environment that may perpetuate neurodegeneration and extracellular matrix degradation. Given milder CSF MMP10 elevations in non-PiD cases, it is plausible that MMP10 captures a response to tissue injury and neuroinflammation that is not limited to PiD but sufficiently pronounced in PiD to result in a signal with diagnostic utility. Our familial *MAPT* results argue against specificity to 3R-tau given observed elevations in a subset of P301L and S305I mutations that are associated with predominantly 4R-tau pathology. Future work should aim to resolve the cellular source(s) of origin and tauopathy-specific mechanisms driving CSF MMP10 alterations in PiD and other proteinopathies.

Several additional subtype biomarker candidates merit follow-up. SomaScan HOMER1 was the strongest pan-TDP candidate that was elevated across TDP subtypes A, B, and C alongside other postsynaptic proteins SHANK3 and DLG4. HOMER1 and SHANK3 participate in an activity-regulated postsynaptic scaffold linking mGluR5 to glutamate receptor signaling^24^, suggesting that this pattern may reflect altered mGluR5-linked synaptic signaling or enrichment of these synaptic constituents in neuronal populations selectively vulnerable to TDP-43 pathology^25^. ISG15, which regulates innate immune cGAS–STING pathways previously implicated in TDP^26^, was selectively elevated in TDPc cases. SomaScan PDE1B, a regulator of glutamatergic synaptic plasticity on striatal spiny projection neurons^27^, emerged as a candidate biomarker that distinguished CBD from other subtypes, including PSP. Additional validation efforts including targeted immunoassays in independent cohorts, similar to the MMP10 work presented here, are needed to move these candidate biomarkers toward clinical applications.

Our multi-platform, multi-disease CSF proteomic network analysis allowed us to directly compare proteomic pathway alterations across early-onset AD and sporadic FTLD, thereby expanding prior work focused exclusively on AD or familial FTLD. Here we show concordant changes in RNA metabolism and immune modules in sporadic FTLD-TDP that we previously observed in familial FTLD-TDP^6^. By contrast, an autophagosome/synaptic vesicle module was selectively elevated in AD and harbored proteins associated with AD co-pathology within FTLD (e.g., SMOC1), consistent with prior findings implicating this pathway in late-onset AD^7^. Across diseases, synaptic/axonal and protein modification modules were the strongest correlates of clinical severity stage and frontotemporal atrophy. These network-based findings help disentangle shared vs. distinct neural injury pathways across proteinopathies, thereby providing important context to prior work restricted to single conditions.

Several limitations should be noted. Autopsy-confirmed FTLD cohorts remain rare and sample sizes with antemortem CSF are smaller than AD-focused studies, particularly for some FTLD subtypes (e.g., TDPb). Nevertheless, we present to our knowledge the largest CSF proteomic cohort with autopsy-confirmed FTLD, with our strongest signal reproducing across orthogonal assays and independent cohorts. Second, SomaScan and Olink are affinity-based platforms that still require additional validation with targeted immunoassays prior to clinical translation. This is particularly relevant for hits such as HOMER1, where differences in detectability and disease associations across platforms suggest possible proteoform effects that may be resolved with additional approaches such as immunoprecipitation mass spectrometry. Third, our study was intentionally CSF-focused because of its proximity to brain and likelihood for producing larger disease signals than blood; however, future work will be needed to determine which targets can be translated into robust antibodies for blood-based biomarkers to support clinical scalability.

Overall, our study interrogating large-scale antemortem CSF proteomics in autopsy-confirmed FTLD provides a framework for biomarker discovery and validation in a population with substantial clinico-pathological heterogeneity and high diagnostic uncertainty. More broadly, the human biofluid proteomic data generated here provide a resource for future biomarker, mechanistic, and therapeutic target studies across FTLD and AD.

## METHODS

### UCSF Fein Memory and Aging Center (MAC) discovery cohort

Participants included 566 individuals enrolled in observational cohort studies at the UCSF Fein MAC. Participants included 274 FTLD spectrum cases, 171 AD spectrum cases, and 95 cognitively unimpaired controls. All participants underwent lumbar puncture for collection of CSF samples, genetic screening, and comprehensive clinico-diagnostic procedures, including neurological examination and informant interview. Clinical diagnoses were made via multidisciplinary consensus case conference with board-certified neurologists and neuropsychologists. Clinical stage was determined with the Clinical Dementia Rating Scale (CDR®) plus Behavioral and Language Domains from the National Alzheimer’s Coordinating Center (NACC) FTLD module (CDR®+NACC-FTLD)^28–30^. CSF samples (n=624) were assayed for large-scale proteomics via the SomaScan v4.1 platform (>7k proteins; 498 subjects and samples) and/or Olink Explore HT platform (>5k proteins; 286 subjects and 344 samples). For Olink, 58 subjects (27 FTLD, 17 AD, 14 CTL) had an additional timepoint of CSF data collection ranging from 0.5 to 3.1 years since baseline (median 1.2 years since baseline). 341 (55%) samples were assayed for pTau181, Aβ42, Aβ40, and NfL via fully automated Elecsys immunoassays (Roche Diagnostics International Ltd, Rotkreuz, Switzerland), as described previously^31^. The UCSF Institutional Review Board approved study procedures and all participants provided written informed consent or assent with proxy consent.

### UCSF MAC clinico-pathological characterization

UCSF MAC FTLD cases included 202 sporadic, symptomatic FTLD cases and 72 cases with an autosomal dominant FTLD-causing mutation (52 symptomatic, 20 asymptomatic). Symptomatic FTLD cases met criteria for a primary FTLD clinical syndrome, specifically behavioral variant FTD (bvFTD)^32^ with or without motor neuron disease (MND)^33^, semantic variant primary progressive aphasia (svPPA)^34^, nonfluent variant PPA (nfvPPA)^34^, corticobasal syndrome with negative AD biomarkers (CBS-FTD)^35^, or progressive supranuclear palsy spectrum syndromes (PSPspectrum)^36^.

AD spectrum individuals had at least one positive AD biomarker based on available CSF pTau181/Aβ42 ratios (>0.022^31^), amyloid PET visual read, postmortem neuropathology, or combinations thereof. 159 AD cases met criteria for cognitive impairment^37,38^ due to AD and 12 were asymptomatic. Symptomatic AD cases were enriched for early-onset clinical variants with a comparable age distribution to FTLD cases, including early-onset amnestic AD (onset before age 65 years; EOAD)^39^, posterior cortical atrophy (PCA)^40^, logopenic variant primary progressive aphasia (lvPPA)^34^, behavioral variant AD (bvAD)^41^, CBS due to AD (CBS-AD)^35^, mild cognitive impairment due to AD (MCI-AD)^38^, or late-onset amnestic AD (onset after age 65 years; LOAD)^37^.

Cognitively unimpaired (CU) control participants were clinically normal, functionally independent (CDR Global = 0), community-dwelling older adults. Of the 95 CU participants, 75 were confirmed AD biomarker negative, while the remaining 20 did not have available AD biomarker data. All participants were free of major neuromedical confounds (e.g., HIV, brain tumor).

A subset of cases went to autopsy with full postmortem neuropathological examinations, in accordance with 2012 NIA-AA guidelines that include characterization of AD, FTLD, and Lewy body disease (LBD) pathology^42,43^. Primary FTLD spectrum neuropathologic diagnoses were made according to 2009 Mackenzie consensus recommendations^44,45^. Autopsy-confirmed FTLD cases included 90 FTLD-tau and 48 FTLD-TDP. The six most common FTLD subtypes were represented, specifically 22 PiD, 28 CBD, 33 PSP, 16 TDPa (14 *GRN* carriers, 1 *C9orf72*), 12 TDPb (4 *C9orf72*), and 16 TDPc. AD cases included 50 individuals with intermediate or high AD neuropathological change (ADNC) and a primary AD clinical syndrome. The Olink cohort also included six cases with a primary pathological diagnosis of LBD that were included as an additional diagnostic group in Olink-specific differential abundance analyses.

### CSF collection and processing

CSF was collected at baseline in polypropylene tubes via lumbar puncture in lateral recumbent or sitting positions. CSF samples were centrifuged at 2000 g for 10 minutes at room temperature. Supernatant was aliquoted in 500 microliter polypropylene tubes and stored at −80°C until further analyses. All CSF samples underwent only one freeze/thaw cycle before analysis.

### UCSF MAC SomaScan proteomics

498 CSF samples across 498 subjects were assayed via the SomaScan v4.1 platform^46^, which measured 7,289 targets across 6,406 unique proteins. The SomaScan assay leverages slow off-rate modified aptamers (SOMAmers), which are short, single stranded deoxynucleotides that bind to protein targets with high specificity^47^. A volume of 65 microliters of CSF was used to create SOMAmer-protein reactions in 96-well plates. Tagged SOMAmer-protein complexes were captured in a bead-based assay, and levels of SOMAmer bound to sample were quantified through a fluorescent signal in DNA hybridization microarrays^46^. The reaction signal was detected digitally and expressed as aggregated Agilent relative fluorescent units (RFU), which were normalized to scale and subsequently log_2_-transformed.

### UCSF MAC Olink proteomics

344 CSF samples across 286 subjects were assayed via the proximity extension assay (PEA) Olink Explore HT platform ^46^, which measured 5,401 unique human proteins. All samples were centrifuged at 10000g for 5 minutes, and 50 μl were aliquoted on 96 well plates and measured alongside QC samples, plate, and negative controls. Following PEA, sequencing was conducted on an Illumina NovaSeqX instrument. Measured protein abundance is reported as intensity-normalized protein expression (NPX). Intra-and inter-run CVs were calculated using three control samples on each plate. Test sample QC was assessed using three internal controls (incubation, extension, and amplification controls) spiked into each sample. Only data from baseline timepoints were used for cross-sectional association analyses.

### Proteomics quality control

Quality control was performed on UCSF MAC SomaScan and Olink datasets using a previously published pipeline^7,9^. Examination of multidimensional scaling (MDS) plots confirmed expected clustering of experimental CSF samples vs. background or control samples. Experimental sample outliers were detected (7 SomaScan, 1 Olink) and removed from analysis using the WGCNA sample network connectivity outlier algorithm (>3 standard deviations [SD] below mean sample connectivity)^48^. Cross-platform correlations for all analytes with matching UniProt accession IDs were conducted using biweight midcorrelations (bicor) and Pearson’s correlations between SomaScan RFU and unlogged Olink NPX (2^NPX^) values. Prior to disease association analyses, we removed analytes with clear bimodal abundance distributions that would violate parametric model assumptions by applying Hartigan’s dip test for unimodality^49^ together with a Gaussian mixture model fit with one or two components. Analytes were classified as bimodal and filtered when the dip test was significant (p<0.05) and the Gaussian mixture model supported two components. We also removed analytes with >75% of values below the limit of detection (LOD), calculated as the median log_2_ signal plus 3 SD of the assay’s buffer (SomaScan) or negative control (Olink) measurements. After bimodality and LOD filtering, 3,713 SomaScan and 1,293 Olink analytes were retained for downstream analysis.

Nonparametric bootstrap regression in control subjects^6^ was performed to estimate ‘typical’ covariance attributable to age, sex, platform-wide principal component 1, and blood contamination, calculated via a composite of blood protein measurements^7^ (SomaScan: HBA1, HBB, HBG1, FGA, FGB, FGG, HP, CAT, CA1; Olink: FGA, FGG, CAT, CA1). Coefficients estimated in controls were then applied to the full protein abundance matrices of cases and controls to derive covariate-adjusted abundance values while preserving disease-related variance. Finally, the adjusted matrices were winsorized at 5 standard deviations to reduce the influence of extreme outlier measurements while preserving the abundance rank order of samples. The resulting outlier-removed, analyte-filtered, covariate-adjusted, and winsorized CSF protein abundance matrices were used for downstream analysis.

### UCSF neuroimaging

UCSF participants with available structural magnetic resonance imaging (MRI) imaging data were included in the analysis of frontotemporal atrophy (n=459). MRI was performed in the UCSF Fein MAC using either a Siemens Trio Tim or Prisma Fit 3T scanner. T1-weighted acquisition parameters were consistent across the Trio Tim and Prisma Fit scanners, with minimal variation in repetition time (TR) and echo time (TE) (sagittal slice orientation; slice thickness = 1.0 mm; slices per slab = 160; in-plane resolution = 1.0 x 1.0 mm; matrix = 240 x 240 x 256; TR = 2,300 ms; Trio Tim TE = 2.98 ms; Prisma Fit TE = 2.9 ms; TI = 900 ms; flip angle = 9°). All T1-weighted images were visually inspected for quality control and images with excessive motion or artifacts were excluded. Magnetic field bias was corrected using N3 algorithm^50^. A T2-weighted FLAIR was acquired with slice thickness = 1.00mm; slices per slab = 176; in-plane resolution = 1.0×1.0mm; matrix = 256×256; TR = 5000 ms, TE = 397 ms, TI = 1800 ms. Tissue volumes were segmented using SPM12^51^. Regional gray matter volume estimates were quantified by translating a standard parcellation atlas into ICBM space and summing the gray matter within each region using the Desikan-Killiany atlas^52^. In addition to individual ROIs, a frontotemporal gray matter volume composite^12^ was constructed by combining frontal and temporal regions of the Desikan-Killiany atlas and then correcting for total intracranial volume and scanner.

### Differential abundance and diagnostic performance

Differential protein abundance for molecular class and pathological subtype models were performed by one-way ANOVA via the parANOVA script, followed by FDR correction for pairwise contrasts. ROC curves for pairwise contrasts were performed via the pROC package. Area under the curve (AUC) estimates and 95% confidence intervals were computed using the DeLong method. Optimal classification thresholds were determined by maximizing the Youden index (sensitivity + specificity − 1), with sensitivity and specificity reported at this threshold. For one-vs-rest subtype classifications, ROC analyses compared each pathological subtype against all other FTLD subtypes plus AD combined. Controls were excluded from rest groups to ensure top AUC estimates captured proteins with differential diagnostic potential across pathological subtypes rather than non-specific neurodegeneration markers that discriminate controls from disease.

### Cross-platform normalization and co-expression network analysis

Olink and SomaScan analytes that passed platform-specific analyte filtering (n=5,006 analytes across platforms) were assembled for 216 participants with CSF measured on both platforms after sample-wise outlier removal. Consistent with prior work, data were transposed prior to cross-platform normalization using the TAMPOR algorithm^53^ (available at https://github.com/edammer/TAMPOR), with proteins treated as samples (columns) and samples as rows for two-way table median polish of ratio. Protein pairs with matching UniProt IDs across platforms (n=1,143) were used as the global internal standard (GIS) to calculate the central tendency within and across platforms for the TAMPOR denominators, as previously described^7,53^. In the case of multiple aptamers with the same UniProt ID, the aptamer with the largest, positive correlation to a matching Olink protein was selected as the GIS. Post-TAMPOR normalized data used in subsequent network analyses were of the form log₂(abundance/central tendency) across the common protein set.

WGCNA^48,54^ was used to generate a CSF protein network from the post-TAMPOR *n*=5,006 log_2_ protein abundance by *n*=216 samples matrix. The WGCNA blockwiseModules function was run with the following parameters: power=13, deepSplit=4, minModuleSize=10, mergeCutHeight=0.07, TOMdenom=“mean”, bicor correlation, signed network type, PAM staging and PAM respects dendro as TRUE, with clustering completed within a single block. Module memberships were then iteratively reassigned to enforce kME table consistency, as previously described^48^. Module eigenproteins, representing the first principal component of abundance for all proteins within a module, were examined in relation to molecular class, FTLD subtype, and disease progression outcomes.

### Gene ontology (GO) and cell type marker enrichment

GO enrichment of differential abundance lists and network modules was calculated as a Fisher exact test p value transformed to z score and visualized using the publicly-available GOparallel code (https://www.github.com/edammer/GOparallel), which pulls curated gene sets from the Bader Lab’s monthly update^55^. As previously published^48^, enrichment of network modules for cell type-specific marker gene symbols was performed using one-tailed Fisher’s exact test (https://github.com/edammer/CellTypeFET).

### Validation Cohorts

### ARTFL/LEFFTDS Longitudinal Frontotemporal Lobar Degeneration (ALLFTD) Consortium

Participants included 166 carriers of pathogenic mutations in the *C9orf72* (n=78)*, GRN* (n=34) or *MAPT* (n=54) genes^56^. Participants were enrolled in the ALLFTD consortium (<u>NCT04363684</u>)^57,58^, which includes over 30 collaborating centers across the US and Canada. The ALLFTD study was approved through the Trial Innovation Network at Johns Hopkins University. Local ethics committees at each of the sites approved the study, and all participants provided written informed consent or assent with proxy consent. Inclusion in the present study required completion of baseline lumbar puncture for CSF collection. Mutation carriers were classified as presymptomatic or symptomatic based on a CDR®+NACC-FTLD Global score of 0 (presymptomatic) or ≥0.5 (symptomatic). All CSF samples were assayed via SomaScan v4.1 and followed the same pre-processing pipeline as UCSF MAC samples.

#### University of Pennsylvania

Participants included 77 individuals with a postmortem neuropathological diagnosis of FTLD (n=57) or AD (n=20) as well as 5 cognitively unimpaired controls with antemortem CSF collected via observational studies at the University of Pennsylvania FTD and AD research centers. FTLD cases included 10 PiD, 10 CBD, 9 PSP, 3 FTLD-tau unclassifiable, 12 TDPa, 8 TDPb, and 5 TDPc. FTLD and AD patients were autopsied at the University of Pennsylvania Center for Neurodegenerative Disease Research. Written informed consent was obtained according to the Declaration of Helsinki and approved by the University of Pennsylvania Institutional Review Board.

#### MMP10 Mesoscale Discovery (MSD) Assay

CSF MMP10 concentrations were measured in the UCSF MAC discovery cohort (N=344 [same samples as Olink experiment]) and University of Pennsylvania validation cohort (N=82). CSF MMP10 was measured by electrochemiluminescence using the commercially available single-analyte U-PLEX Human MMP10 assay (MSD, Rockville, MD, USA). CSF samples were thawed and prepared according to the manufacturer’s instructions, then loaded onto the assay plate in duplicate wells. Samples were measured in duplicate with a CSF dilution factor of 1:4. Duplicate measurements were averaged for downstream analyses. The lower limit of detection (LLOD) was 0.37 pg/mL.

#### Statistical Analysis

Statistical analyses were performed in R (v4.6.1). Correlations were performed using biweight midcorrelations (bicor) or Pearson’s correlations where indicated. Comparisons between two groups were performed using *t* tests. Comparisons among three or more groups were performed with ANOVA and pairwise comparisons of significance. *P* values were adjusted for multiple comparisons by FDR correction according to the Benjamini-Hochberg method where indicated.

## Data Availability

Summary statistics and visualizations for limit of detection, cross-platform correlation, differential abundance, disease severity correlations, and module membership are available for all analytes, irrespective of QC status, via a ShinyApp: https://salonerlab.shinyapps.io/MACWideShinyApp/. Raw data are available upon reasonable request with formal applications to each respective study. UCSF Fein MAC data can be requested at https://memory.ucsf.edu/research-trials/professional/open-science. ALLFTD data can be requested at https://www.allftd.org/data. University of Pennsylvania data can be requested at https://www.pennbindlab.com/data-sharing. Certain data elements may be restricted due to the potential for identifiability in the context of the sensitive nature of genetic data.

## Code Availability

Algorithms used for protein data processing and analysis are available in existing R packages and via GitHub at https://github.com/edammer, as described in the Methods.

## Author Contributions

R.S., W.W.S., J.C.R. and A.L.B. conceptualized the study. R.S. led the analyses and produced the figures. J.D., L.C., C.D., and M.O. provided additional analytic support. R.S. has full access to the data in the study and takes responsibility for the integrity of the data and the accuracy of the data analysis. B.C. and J.C.R. performed the MMP10 immunoassay. M.H. and A.A. performed Olink proteomics. R.S. drafted the manuscript. R.S., K.A.Q.C., D.W., E.L., D.I., G.D.R., J.H.K., K.B.C., B.F.B., H.J.R., J.S.Y., B.L.M., W.W.S., J.C.R., and A.L.B. obtained funding. All authors critically revised the paper for intellectual content. All authors read and approved the final version of the manuscript.

## Acknowledgments

We are deeply grateful to the study participants and their families for participating in the affiliated research programs. We additionally thank the teams of coordinators and administrative staffs who have provided integral support for the ongoing execution of research at our centers. We also thank Dr. Eric Dammer for consultation on proteomic preprocessing and bioinformatic pipelines.

## Funding

Data collection and dissemination of the data presented in this paper were supported by funding support to the UCSF MAC, including National Institutes of Health (NIH) National Institute on Aging (NIA) for Frontotemporal Dementia: Genes, Images, and Emotions: P01 AG019724 (B.L.M.), UCSF Alzheimer’s Disease Research Center: P30 AG062422 (G.D.R.), and UCSF Brain Aging Network for Cognitive Health: R01AG032289 (J.H.K.), R01AG048234 (J.H.K.). This work is supported by additional NIH funding (grant nos. K23AG090757 (R.S.), K23AG084883 (E.W.P.), R01AG038791 (A.L.B.), R01AG032306 (H.J.R.), K23AG059888 (J.C.R.), K23AG073514 (L.V.), P01AG019724 (B.L.M.), P30AG062422 (W.W.S.), R01AG072475 (K.B.C.), K23AG061253 (A.M.S.), K24AG045333 (H.J.R.)). The ALLFTD Consortium (U19: AG063911 [A.L.B., H.J.R. and B.F.B.]) is funded by the NIA and the National Institute of Neurological Diseases and Stroke (NINDS); the former ARTFL and LEFFTDS Consortia (ARTFL: U54 NS092089 [A.L.B., H.J.R. and B.F.B.]) were funded by NINDS and National Center for Advancing Translational Sciences; and LEFFTDS (U01 AG045390 [H.J.R. and B.F.B.]) was funded by the NIA and NINDS. ALLFTD samples are stored at the National Centralized Repository for Alzheimer’s Disease and Related Dementias (NCRAD), which receives government support under a cooperative agreement grant (U24 AG21886) awarded by the NIA. The UPenn ADRC is supported by NIA grant P01AG066597 and FTD Center is supported by NIA grant P30AG072979. The UPenn Brain Bank is supported by NIA grants P30AG072979, P01AG066597 and P01AG084497.

Additional support comes from the Alzheimer’s Association (grant nos. AARF-23-1145318 (R.S.) and AARG-20-683875 (K.B.C.)), New Vision Research (grant no. CCAD 2024-001-1 (R.S.)), an American Academy of Neurology, American Brain Foundation, and Association for Frontotemporal Degeneration Fellowship Award (R.S.), the Bluefield Project to Cure FTD, CurePSP, Larry L. Hillblom Foundation (grant nos. 2018-A-006-NET (J.H.K.) and 2018-A-025-FEL (A.M.S.)), and the Rainwater Charitable Foundation.

**Supplementary Figure 1. Proteomic quality control (QC) pipeline for SomaScan and Olink CSF datasets. a**, Sample filtering. Multidimensional scaling (MDS) plots of leading log-fold-change dimensions for SomaScan (top) and Olink (bottom), showing all samples (left; colored by diagnostic group [Control, AD, FTLD, Other] and control-sample type [SomaScan: Buffer, Calibrator, QC; Olink: Plate control, Sample control, Negative control]) and experimental samples before (middle) and after (right) removal of outlier samples with aberrant network connectivity (WGCNA sample network connectivity algorithm, >3 SD below mean connectivity; SomaScan, n = 7 removed; Olink, n = 1 removed). **b**, Assay filtering. Schematic of sequential analyte filtering: assays with bimodal abundance distributions (Hartigan’s dip test P < 0.05 and a two-component Gaussian mixture model fit; example, ABO) were removed, followed by assays with >75% of values below the limit of detection (LOD; median log2 buffer [SomaScan] or negative control [Olink] signal + 3 SD; example, AGR2, dashed line denotes LOD). Assay counts remaining after each QC step, by platform (top right). Proportion of assays reaching nominal significance (P < 0.05, all disease cases combined vs. CTL) among assays passing vs. failing QC, by platform (bottom right). **c**, Cross-platform correlation. Distribution of biweight midcorrelations (bicor) between SomaScan and Olink abundance for 2,663 matched UniProt-ID analyte pairs across 216 samples assayed on both platforms, colored by QC status (passed QC on both platforms, one platform only, or failed QC on both platforms). Box plot (right) of cross-platform bicor by QC status group. **d**, Covariate regression. Percent variance in protein abundance attributable to principal component 1 (PC1), blood eigenprotein contamination score (BloodEP), age, sex, and diagnosis (Dx), estimated by variance partition analysis, before (left) and after (right) covariate regression to remove nuisance covariance, for SomaScan (top) and Olink (bottom). Each point represents one protein/analyte; violin denotes the distribution of residual (unexplained) variance. Box plots (**b**, **c**) show median, 25th–75th percentiles (box) and range (whiskers), with individual data points overlaid.

**Supplementary Figure 2. Cross-cohort effect size concordance with an independent Olink FTLD proteomics dataset.** Comparison of log2 fold change (present study, x-axis) with log2 fold change (biweight midcorrelation [bicor]) reported by Hok-A-Hin et al.¹ (Olink; y-axis) for matched analyte pairs, across SomaScan (left column) and Olink (right column) platforms, for FTLD-TDP vs. CTL (top), FTLD-tau vs. CTL (middle), and FTLD-TDP vs. FTLD-tau (bottom). Points are colored by nominal significance (P < 0.05) in the present study’s x-axis contrast only, the Hok-A-Hin et al. y-axis contrast only, or both.

**Supplementary Figure 3. Olink HOMER1 abundance across FTLD molecular classes and subtypes and cross-platform concordance with SomaScan. a**, Olink HOMER1 abundance (NPX) by molecular class (FTLD-TDP, FTLD-tau, AD, CTL). *P < 0.05, ***P < 0.001 (nominal P pairwise contrasts). **b**, Olink HOMER1 abundance (NPX) by pathological subtype (TDPa, TDPb, TDPc, PiD, CBD, PSP, AD, LBD, CTL). P value, one-way ANOVA. **c**, Pearson correlation between SomaScan HOMER1 (aptamer seq.19229.92; RFU) and Olink HOMER1 (2^NPX) abundance, with linear fit and 95% confidence band (shaded). Box plots (**a**, **b**) show median, 25th–75th percentiles (box) and range (whiskers), with individual data points overlaid.

**Supplementary Figure 4. Multi-platform proteomic network harmonization removes platform effects. a**, Multidimensional scaling (MDS) plots of all post-QC analytes (SomaScan and Olink) across the n = 216 samples assayed on both platforms, before (Pre-TAMPOR, left) and after (Post-TAMPOR, right) cross-platform normalization, colored by platform. **b**, MDS plots restricted to post-QC analyte pairs with matching UniProt IDs across platforms (global internal standard [GIS] analytes; n = 1,143) used to anchor TAMPOR normalization, before (left) and after (right) harmonization, colored by platform. Four representative matched analyte pairs (NEFL, MMP10, SMOC1, NPTX2) are highlighted to illustrate their convergence from platform-driven separation (Pre-TAMPOR) to cross-platform concordance (Post-TAMPOR).

**Supplementary Figure 5. Protein co-expression network biological enrichment.** Gene ontology (GO) and pathway analysis was performed to ascertain the principal biology represented by the constituent proteins in each protein co-expression module. Enrichment for a given ontology is shown by z score, transformed from a Fisher’s exact test.

---

**Extended Data Figure 1.**
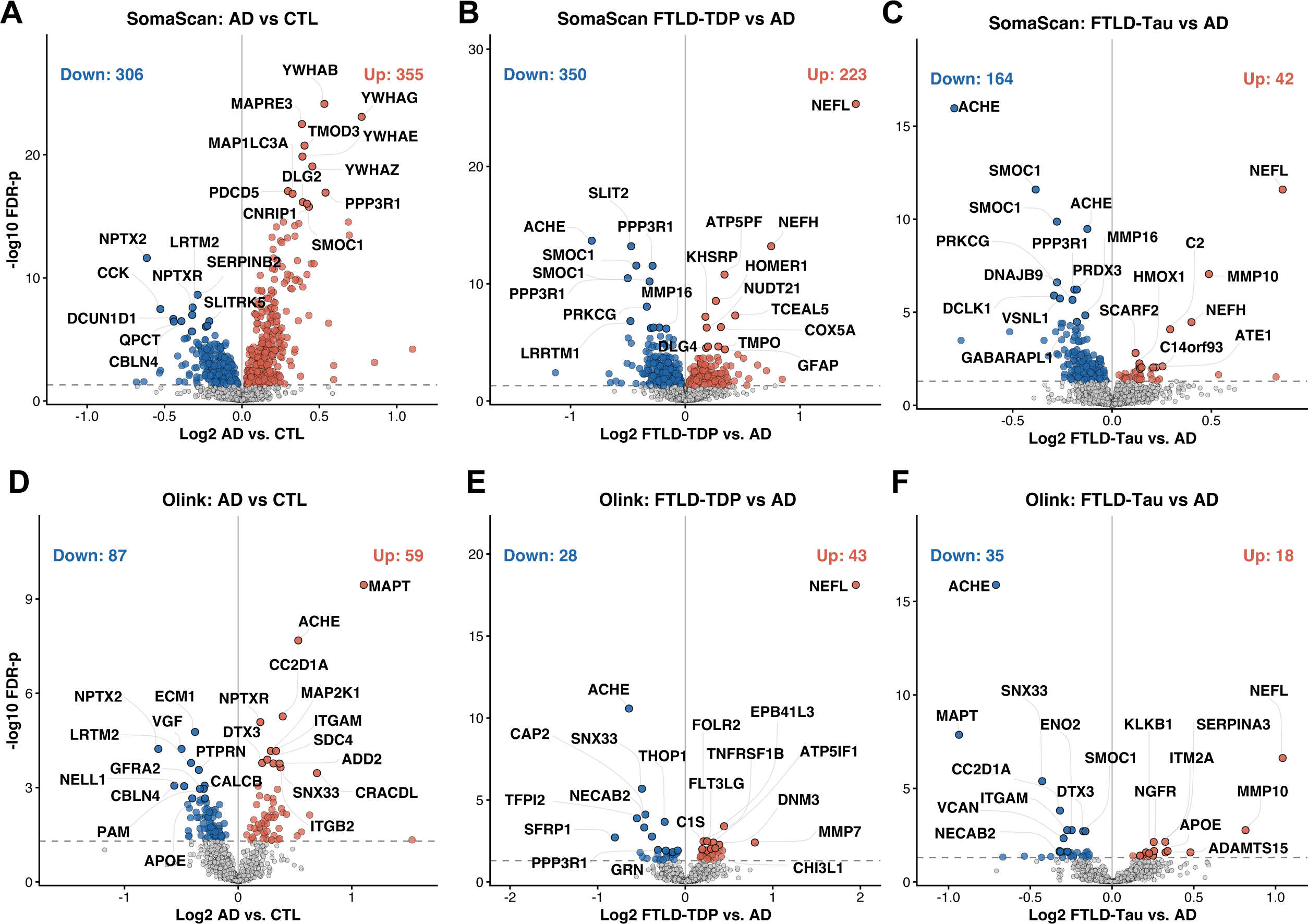
CSF proteomics differentiates FTLD from AD. **a–c**, Volcano plots of SomaScan differential protein abundance for **a,** AD vs. CTL, **b,** FTLD-TDP vs. AD, and **c,** FTLD-tau vs. AD. Dashed line, FDR-P = 0.05; counts denote aptamers reaching FDR significance in each direction (up=red, down=blue). **d–f,** Volcano plots of Olink differential protein abundance for the same contrasts as **a–c**: **d,** AD vs. CTL, **e,** FTLD-TDP vs. AD, **f,** FTLD-tau vs. AD.

**Extended Data Figure 2.**
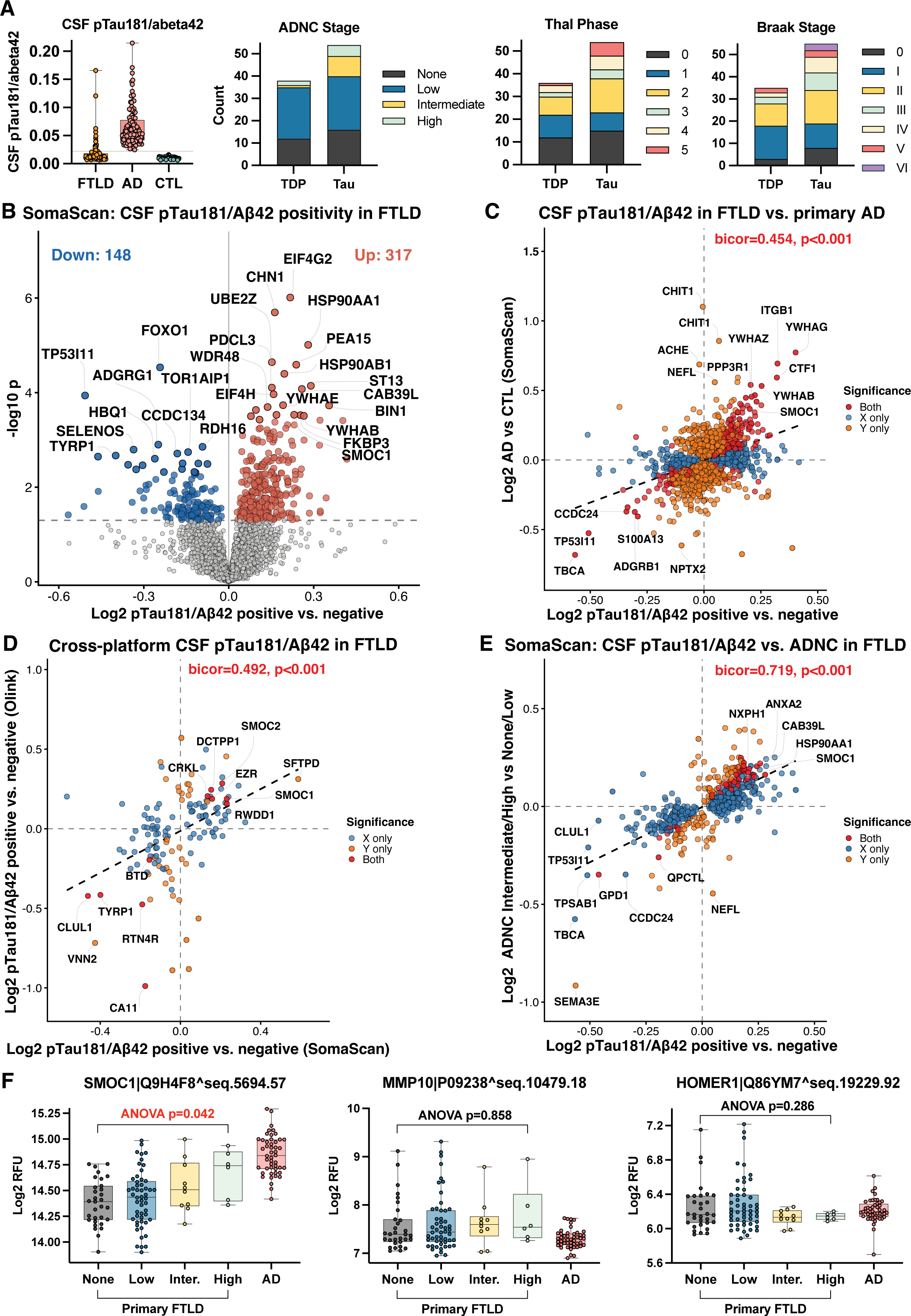
CSF proteomic signatures of AD co-pathology within FTLD. **a**, CSF pTau181/Aβ42 ratio (Elecsys immunoassay) by molecular class (FTLD, AD, CTL); horizontal line denotes the pTau181/Aβ42 positivity cut-off (0.022). Distribution of ADNC (AD neuropathologic change) stage, Thal amyloid phase, and Braak neurofibrillary tangle stage among autopsy-confirmed FTLD-TDP and FTLD-tau cases. **b**, Volcano plot of SomaScan differential protein abundance for CSF pTau181/Aβ42-positive (n = 18) vs. -negative (n = 88) FTLD-spectrum cases. Dashed line, nominal P= 0.05; counts denote aptamers reaching nominal significance in each direction (up = red, down = blue). **c**, Comparison of SomaScan log2 fold change for CSF pTau181/Aβ42 positivity in FTLD (x-axis) vs. primary AD vs. CTL (y-axis; biweight midcorrelation [bicor]), restricted to aptamers with nominal P < 0.05 in either contrast and colored by significance. **d**, Cross-platform comparison of log2 fold change for CSF pTau181/Aβ42 positivity in FTLD between SomaScan (x-axis) and Olink (y-axis; bicor), restricted to matched analyte pairs with nominal P < 0.05 on either platform and colored by platform significance. **e**, Comparison of SomaScan log2 fold change for CSF pTau181/Aβ42 positivity (x-axis) vs. ADNC intermediate/high vs. none/low (y-axis; bicor) in autopsy-confirmed FTLD (n = 17 intermediate/high vs. n = 40 none/low), restricted to aptamers with nominal P < 0.05 in either contrast and colored by significance. **f**, SomaScan abundance (log2 RFU) of SMOC1, MMP10, and HOMER1 by ADNC stage (none, low, intermediate, high) in autopsy-confirmed primary FTLD, with primary AD shown for reference. P values, one-way ANOVA across FTLD ADNC stages. Box plots (a, f) show median, 25th–75th percentiles (box) and range (whiskers), with individual data points overlaid.

**Extended Data Figure 3.**
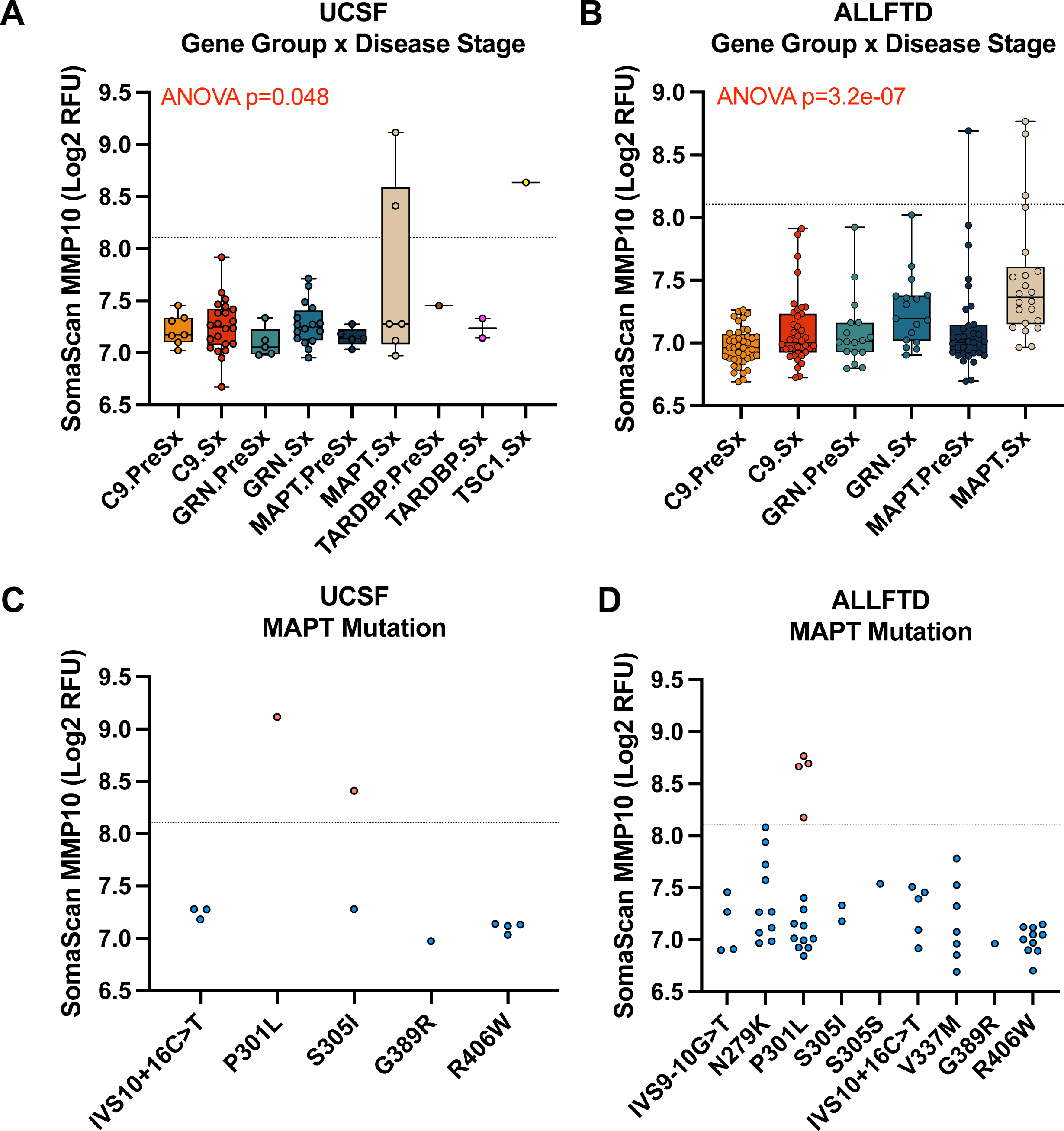
CSF MMP10 is elevated in a subset of *MAPT* mutation carriers. **a**, SomaScan MMP10 abundance by familial FTLD gene group and disease stage (presymptomatic [PreSx] or symptomatic [Sx] *C9orf72*, *GRN*, *MAPT*, *TARDBP,* and *TSC1* mutation carriers) in the UCSF cohort. P value, one-way ANOVA. **b,** SomaScan MMP10 abundance by specific *MAPT* mutation in the UCSF cohort. **c,** SomaScan MMP10 abundance by familial FTLD gene group and disease stage (*C9orf72*, *GRN*, *MAPT*) in the ALLFTD cohort. P value, one-way ANOVA. **d,** SomaScan MMP10 abundance by specific *MAPT* mutation in the ALLFTD cohort. Dashed line (**a–d**), PiD classification cut-off (from Fig. 3d). Box plots show median, 25th–75th percentiles (box) and range (whiskers), with individual data points overlaid.

