## Supplemental Figures for "Large-scale cerebrospinal fluid proteomics identifies diagnostic biomarker candidates across the clinico-pathological continuum of frontotemporal lobar degeneration"

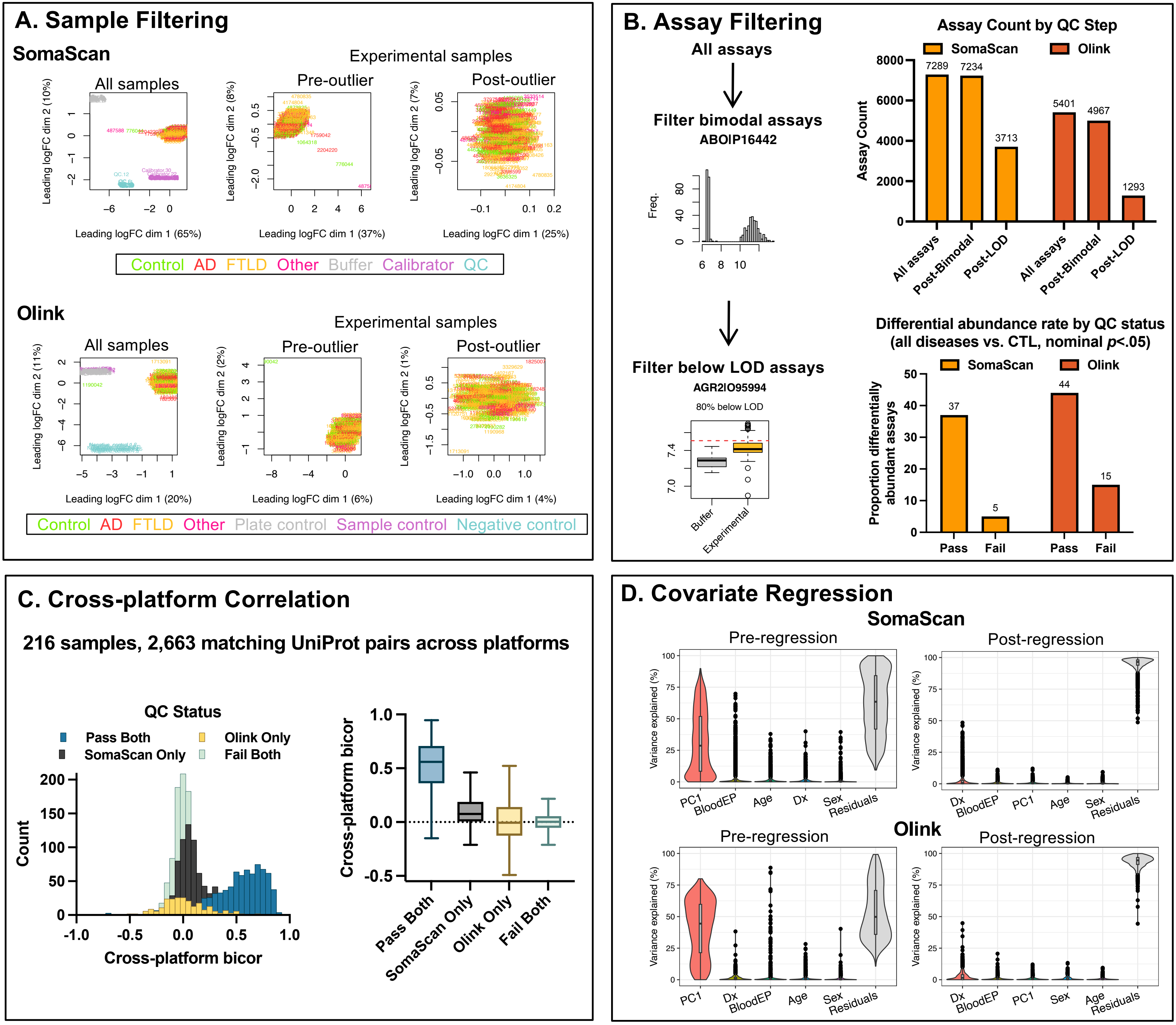
A. Sample Filtering
SomaScan
Experimental samples
All samples
Pre-outlier
Post-outlier
Leading logFC dim 2 (10%)
Leading logFC dim 2 (8%)
Leading logFC dim 2 (7%)
Leading logFC dim 1 (65%)
Leading logFC dim 1 (37%)
Leading logFC dim 1 (25%)
Control AD FTLD Other Buffer Calibrator QC
Olink
Experimental samples
All samples
Pre-outlier
Post-outlier
Leading logFC dim 2 (11%)
Leading logFC dim 2 (2%)
Leading logFC dim 2 (1%)
Leading logFC dim 1 (20%)
Leading logFC dim 1 (6%)
Leading logFC dim 1 (4%)
Control AD FTLD Other Plate control Sample control Negative control
B. Assay Filtering
All assays
Filter bimodal assays
ABOIP16442
Freq.
Filter below LOD assays
AGR2IO95994
80% below LOD
Buffer
Experimental
Assay Count by QC Step
SomaScan
Olink
Assay Count
7289
7234
3713
5401
4967
1293
All assays
Post-Bimodal
Post-LOD
All assays
Post-Bimodal
Post-LOD
Differential abundance rate by QC status
(all diseases vs. CTL, nominal p<.05)
SomaScan
Olink
Proportion differentially abundant assays
37
5
44
15
Pass
Fail
Pass
Fail
C. Cross-platform Correlation
216 samples, 2,663 matching UniProt pairs across platforms
QC Status
Pass Both
SomaScan Only
Olink Only
Fail Both
Count
Cross-platform bicor
Cross-platform bicor
Pass Both
SomaScan Only
Olink Only
Fail Both
D. Covariate Regression
SomaScan
Pre-regression
Post-regression
Variance explained (%)
PC1
BloodEP
Age
Dx
Sex
Residuals
Dx
BloodEP
PC1
Age
Sex
Residuals
Olink
Pre-regression
Post-regression
PC1
Dx
BloodEP
Age
Sex
Residuals
Dx
BloodEP
PC1
Sex
Age
Residuals

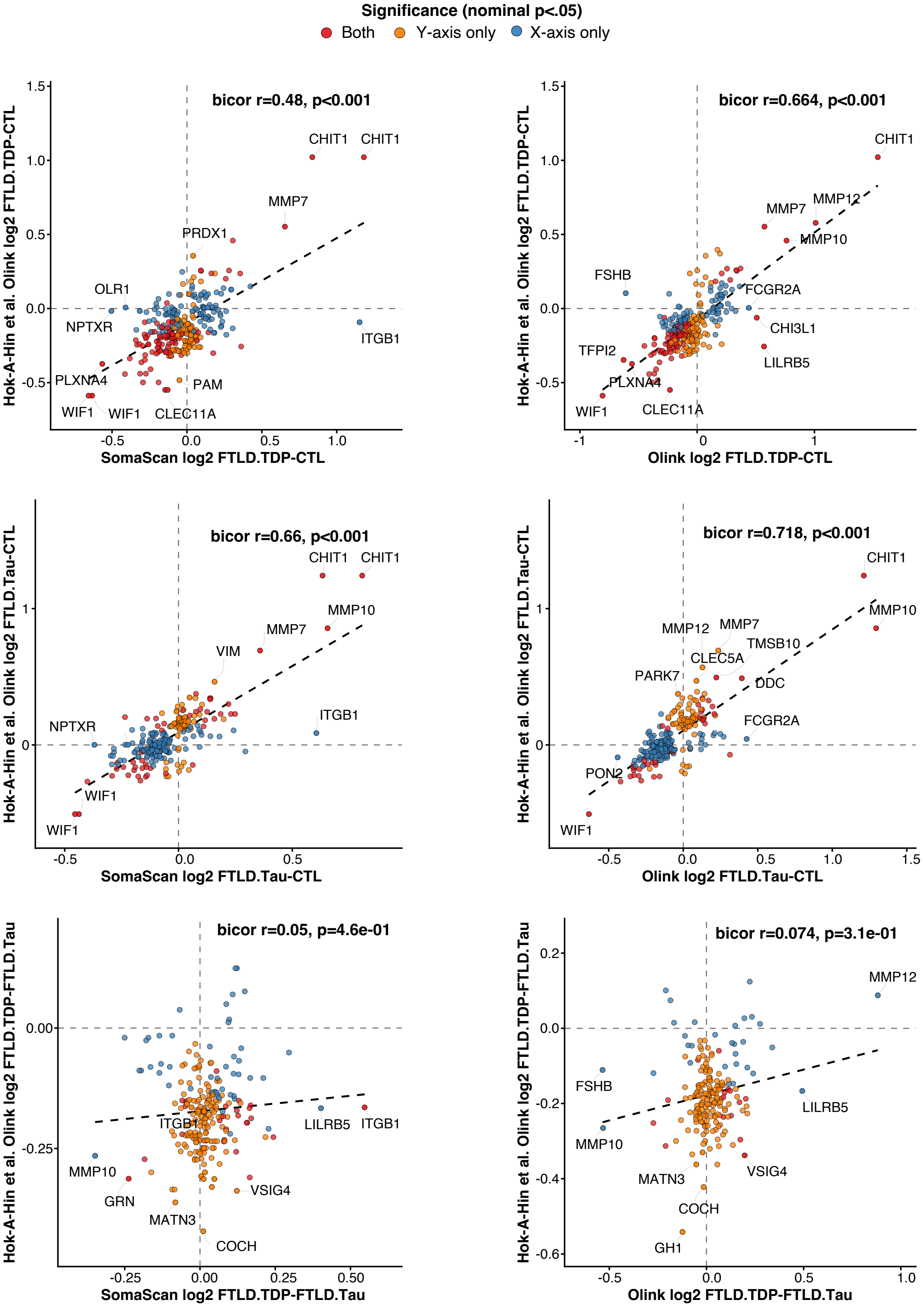

Significance (nominal p<.05)
Both
Y-axis only
X-axis only
bicor r=0.48, p<0.001
Hok-A-Hin et al. Olink log2 FTLD.TDP-CTL
SomaScan log2 FTLD.TDP-CTL
CHIT1
CHIT1
MMP7
PRDX1
OLR1
NPTXR
ITGB1
PLXNA4
PAM
WIF1
WIF1
CLEC11A
bicor r=0.664, p<0.001
Olink log2 FTLD.TDP-CTL
CHIT1
MMP12
MMP7
MMP10
FSHB
FCGR2A
CHI3L1
TFPI2
LILRB5
PLXNA4
WIF1
CLEC11A
bicor r=0.66, p<0.001
Hok-A-Hin et al. Olink log2 FTLD.Tau-CTL
SomaScan log2 FTLD.Tau-CTL
CHIT1
CHIT1
MMP10
MMP7
VIM
ITGB1
NPTXR
WIF1
WIF1
bicor r=0.718, p<0.001
Olink log2 FTLD.Tau-CTL
CHIT1
MMP10
MMP12
MMP7
TMSB10
CLEC5A
PARK7
DDC
FCGR2A
PON2
WIF1
bicor r=0.05, p=4.6e-01
Hok-A-Hin et al. Olink log2 FTLD.TDP-FTLD.Tau
SomaScan log2 FTLD.TDP-FTLD.Tau
ITGB1
LILRB5
ITGB1
MMP10
GRN
VSIG4
MATN3
COCH
bicor r=0.074, p=3.1e-01
Olink log2 FTLD.TDP-FTLD.Tau
MMP12
FSHB
LILRB5
MMP10
VSIG4
MATN3
COCH
GH1

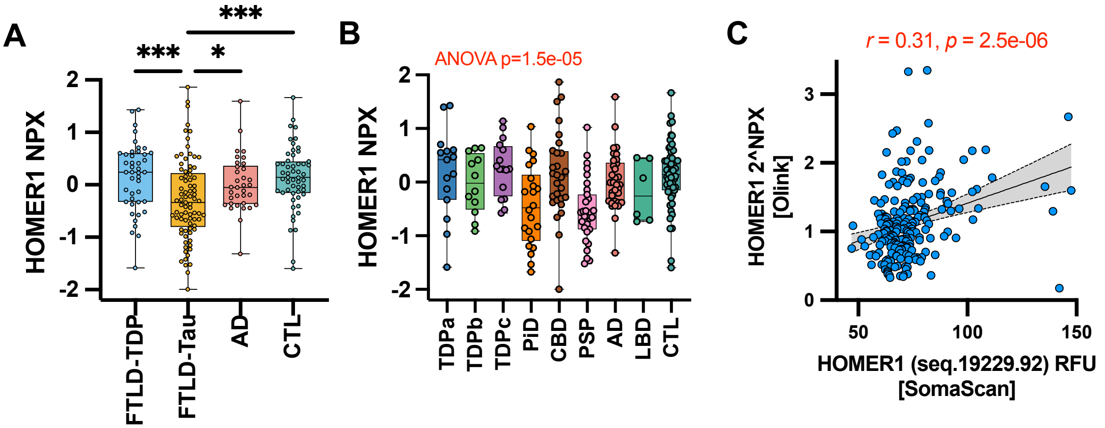

A
***
***
*
HOMER1 NPX
2
1
0
-1
-2
FTLD-TDP
FTLD-Tau
AD
CTL
B
ANOVA p=1.5e-05
HOMER1 NPX
TDPa
TDPb
TDPc
PiD
CBD
PSP
AD
LBD
CTL
C
r = 0.31, p = 2.5e-06
HOMER1 2^NPX
[Olink]
3
2
1
0
50
100
150
HOMER1 (seq.19229.92) RFU
[SomaScan]

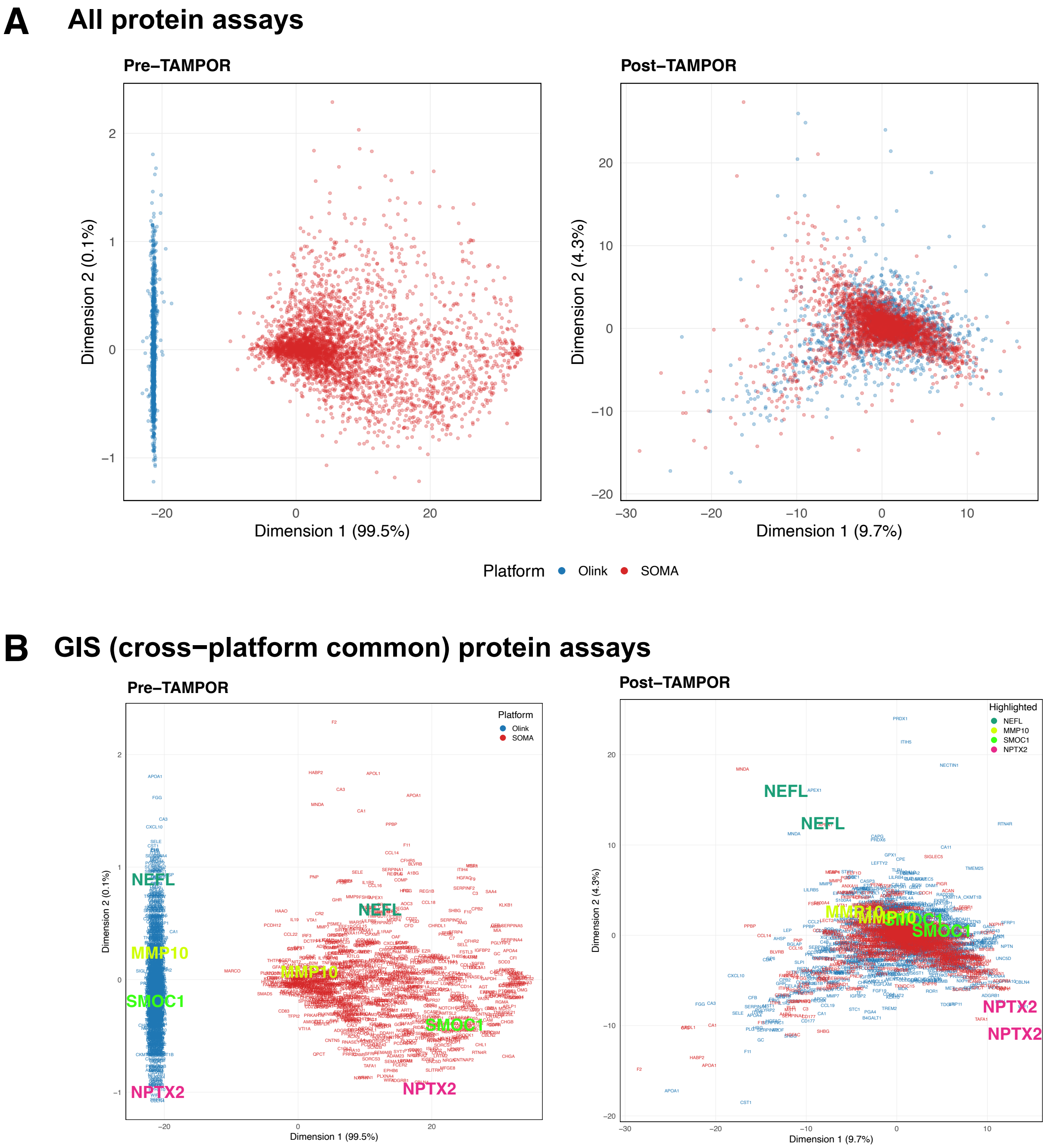
A
All protein assays
Pre−TAMPOR
Dimension 2 (0.1%)
Dimension 1 (99.5%)
Post−TAMPOR
Dimension 2 (4.3%)
Dimension 1 (9.7%)
Platform
Olink
SOMA
B
GIS (cross−platform common) protein assays
Pre−TAMPOR
Platform
Olink
SOMA
NEFL
MMP10
SMOC1
NPTX2
Dimension 2 (0.1%)
Dimension 1 (99.5%)
Post−TAMPOR
Highlighted
NEFL
MMP10
SMOC1
NPTX2
Dimension 2 (4.3%)
Dimension 1 (9.7%)

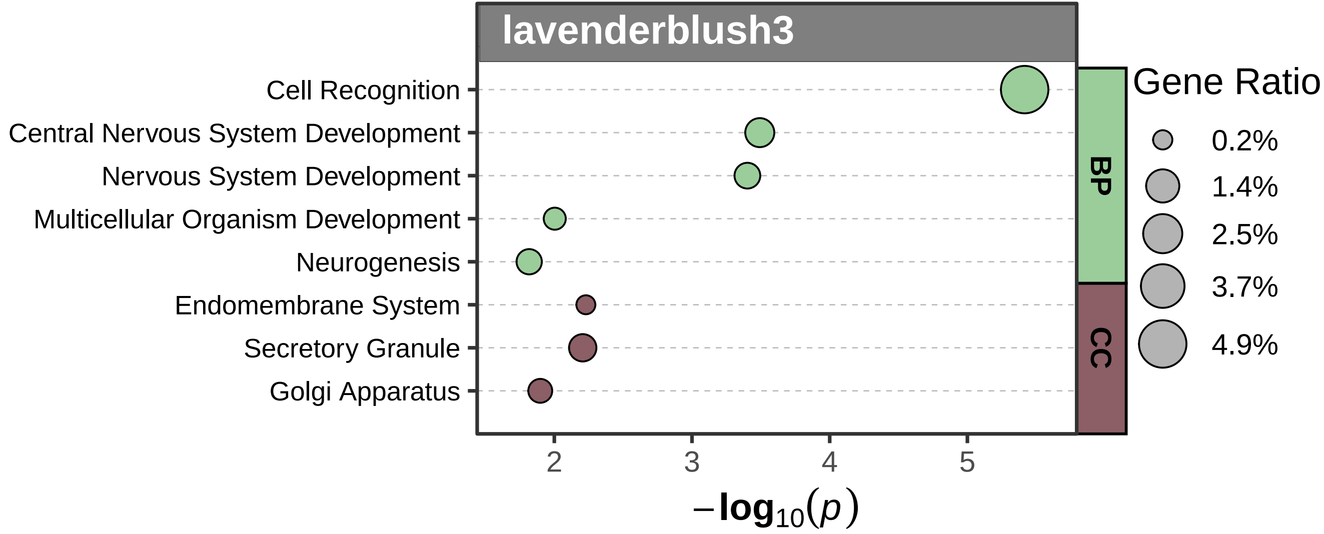

lavenderblush3
Cell Recognition
Central Nervous System Development
Nervous System Development
Multicellular Organism Development
Neurogenesis
Endomembrane System
Secretory Granule
Golgi Apparatus
BP
CC
Gene Ratio
0.2%
1.4%
2.5%
3.7%
4.9%
2
3
4
5
$-\log_{10}(p)$

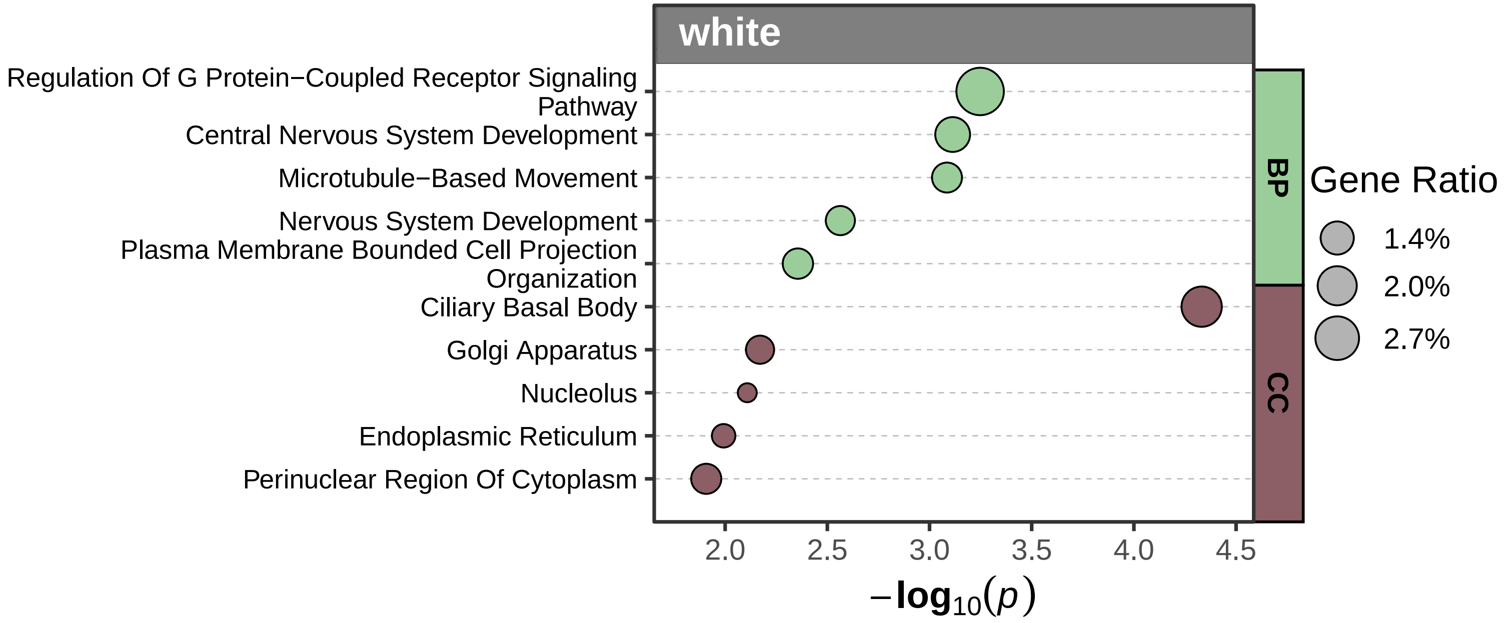

white
Regulation Of G Protein−Coupled Receptor Signaling Pathway
Central Nervous System Development
Microtubule−Based Movement
Nervous System Development
Plasma Membrane Bounded Cell Projection Organization
Ciliary Basal Body
Golgi Apparatus
Nucleolus
Endoplasmic Reticulum
Perinuclear Region Of Cytoplasm
BP
CC
Gene Ratio
1.4%
2.0%
2.7%
2.0
2.5
3.0
3.5
4.0
4.5
−log10(p)

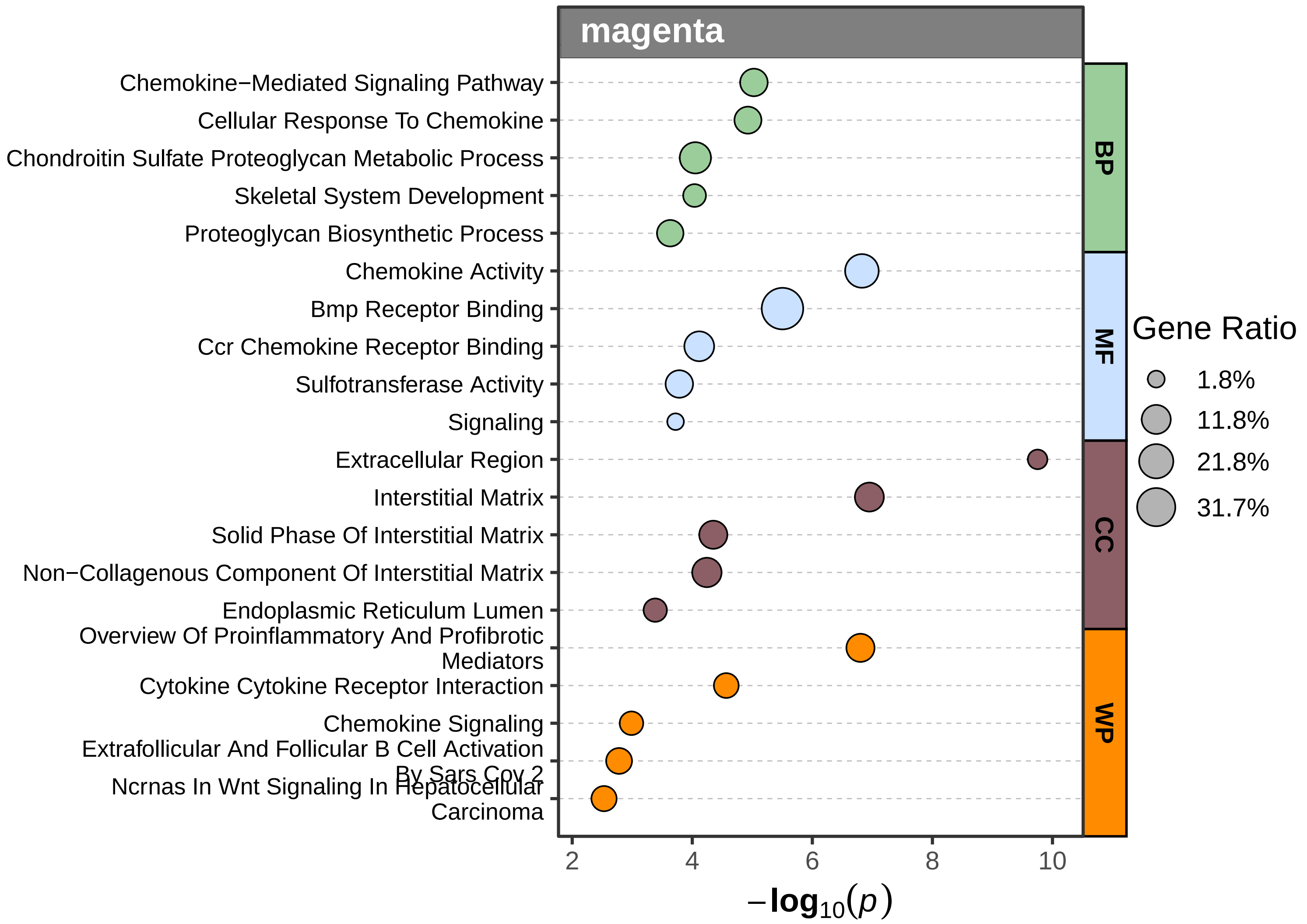
magenta
Chemokine−Mediated Signaling Pathway
Cellular Response To Chemokine
Chondroitin Sulfate Proteoglycan Metabolic Process
Skeletal System Development
Proteoglycan Biosynthetic Process
Chemokine Activity
Bmp Receptor Binding
Ccr Chemokine Receptor Binding
Sulfotransferase Activity
Signaling
Extracellular Region
Interstitial Matrix
Solid Phase Of Interstitial Matrix
Non−Collagenous Component Of Interstitial Matrix
Endoplasmic Reticulum Lumen
Overview Of Proinflammatory And Profibrotic Mediators
Cytokine Cytokine Receptor Interaction
Chemokine Signaling
Extrafollicular And Follicular B Cell Activation By Sars Cov 2
Ncrnas In Wnt Signaling In Hepatocellular Carcinoma
BP
MF
CC
WP
Gene Ratio
1.8%
11.8%
21.8%
31.7%
2
4
6
8
10
$-\log_{10}(p)$

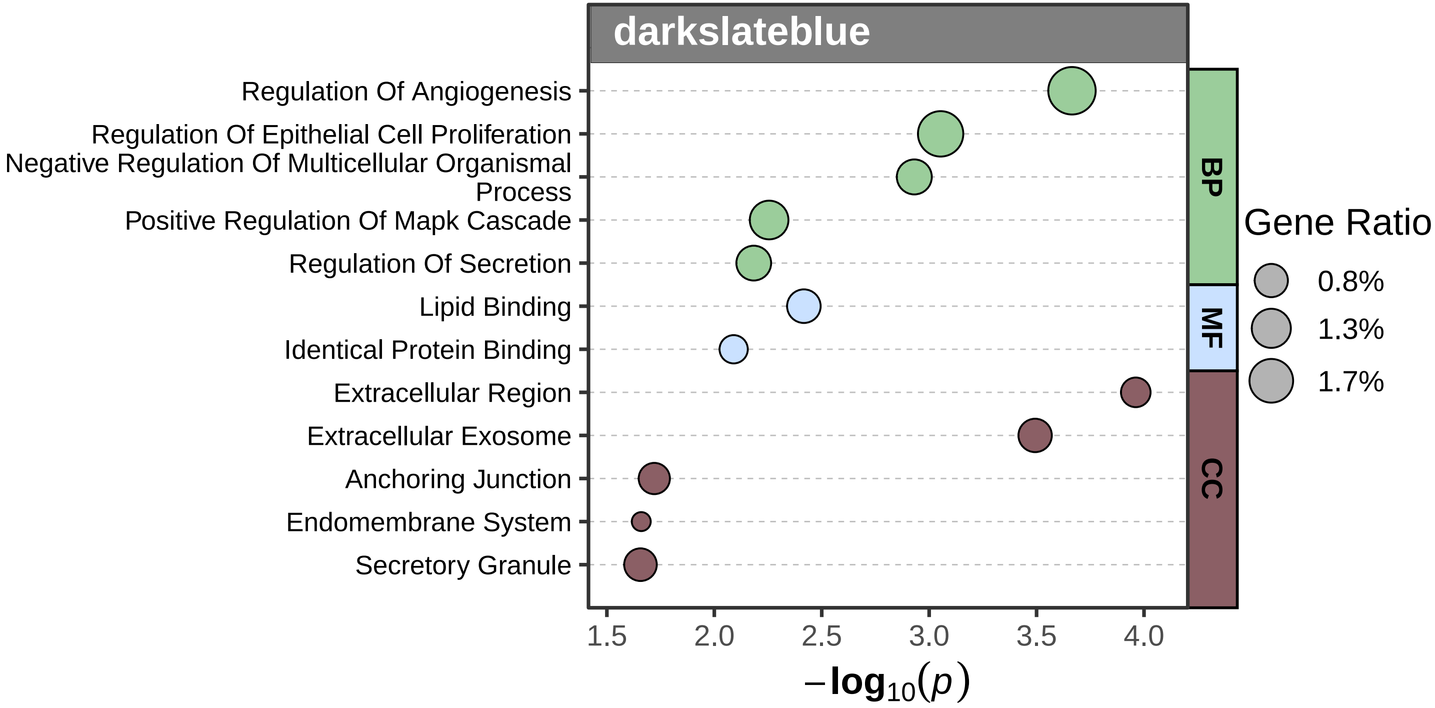
darkslateblue
Regulation Of Angiogenesis
Regulation Of Epithelial Cell Proliferation
Negative Regulation Of Multicellular Organismal Process
Positive Regulation Of Mapk Cascade
Regulation Of Secretion
Lipid Binding
Identical Protein Binding
Extracellular Region
Extracellular Exosome
Anchoring Junction
Endomembrane System
Secretory Granule
BP
MF
CC
Gene Ratio
0.8%
1.3%
1.7%
1.5
2.0
2.5
3.0
3.5
4.0
$-\log_{10}(p)$

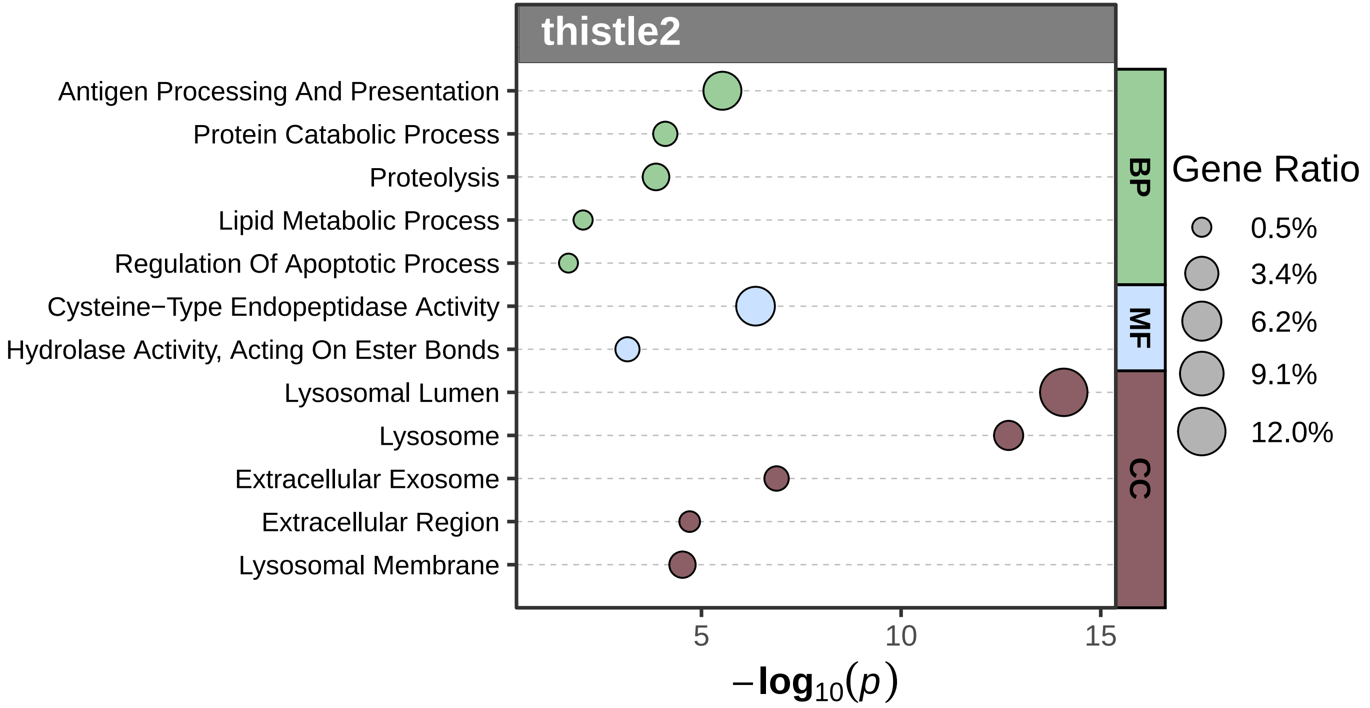

thistle2
Antigen Processing And Presentation
Protein Catabolic Process
Proteolysis
Lipid Metabolic Process
Regulation Of Apoptotic Process
Cysteine−Type Endopeptidase Activity
Hydrolase Activity, Acting On Ester Bonds
Lysosomal Lumen
Lysosome
Extracellular Exosome
Extracellular Region
Lysosomal Membrane
BP
MF
CC
Gene Ratio
0.5%
3.4%
6.2%
9.1%
12.0%
5
10
15
$-\log_{10}(p)$

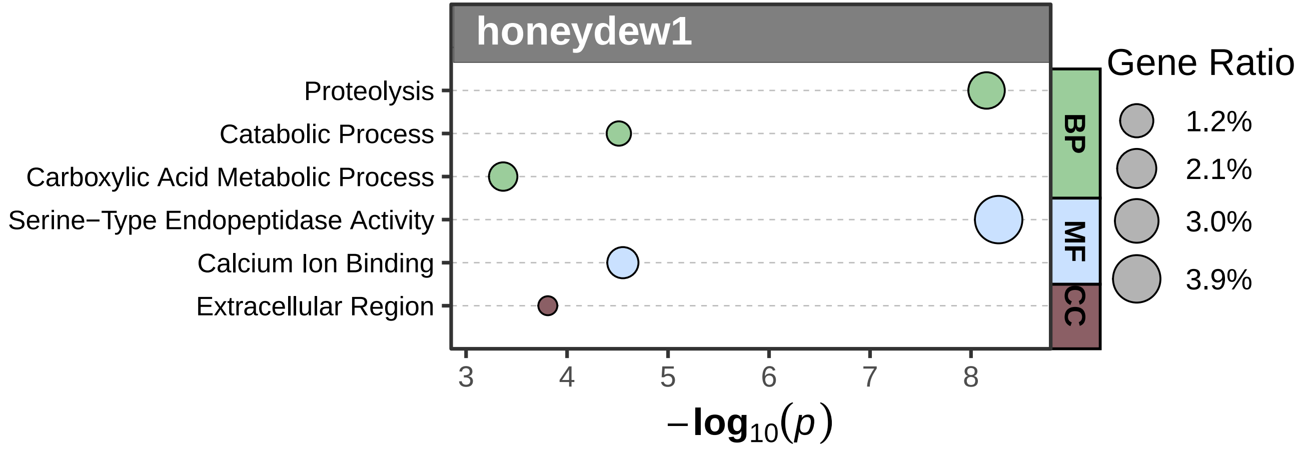

honeydew1
Proteolysis
Catabolic Process
Carboxylic Acid Metabolic Process
Serine−Type Endopeptidase Activity
Calcium Ion Binding
Extracellular Region
BP
MF
CC
Gene Ratio
1.2%
2.1%
3.0%
3.9%
3
4
5
6
7
8
$-\log_{10}(p)$

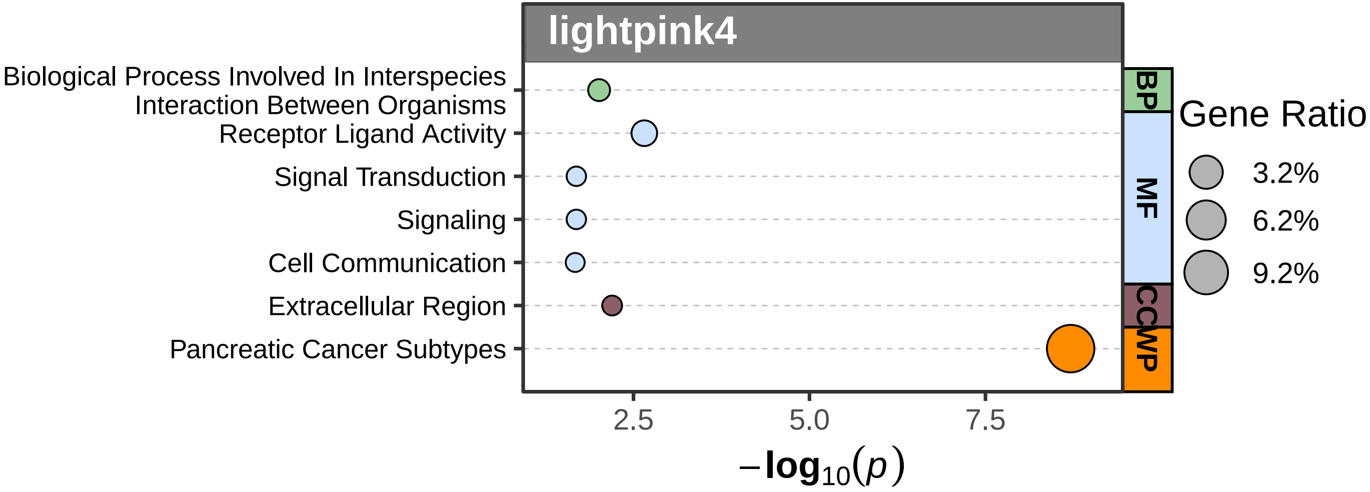

lightpink4
Biological Process Involved In Interspecies Interaction Between Organisms
Receptor Ligand Activity
Signal Transduction
Signaling
Cell Communication
Extracellular Region
Pancreatic Cancer Subtypes
BP
MF
CC
WP
2.5
5.0
7.5
$-\log_{10}(p)$
Gene Ratio
3.2%
6.2%
9.2%

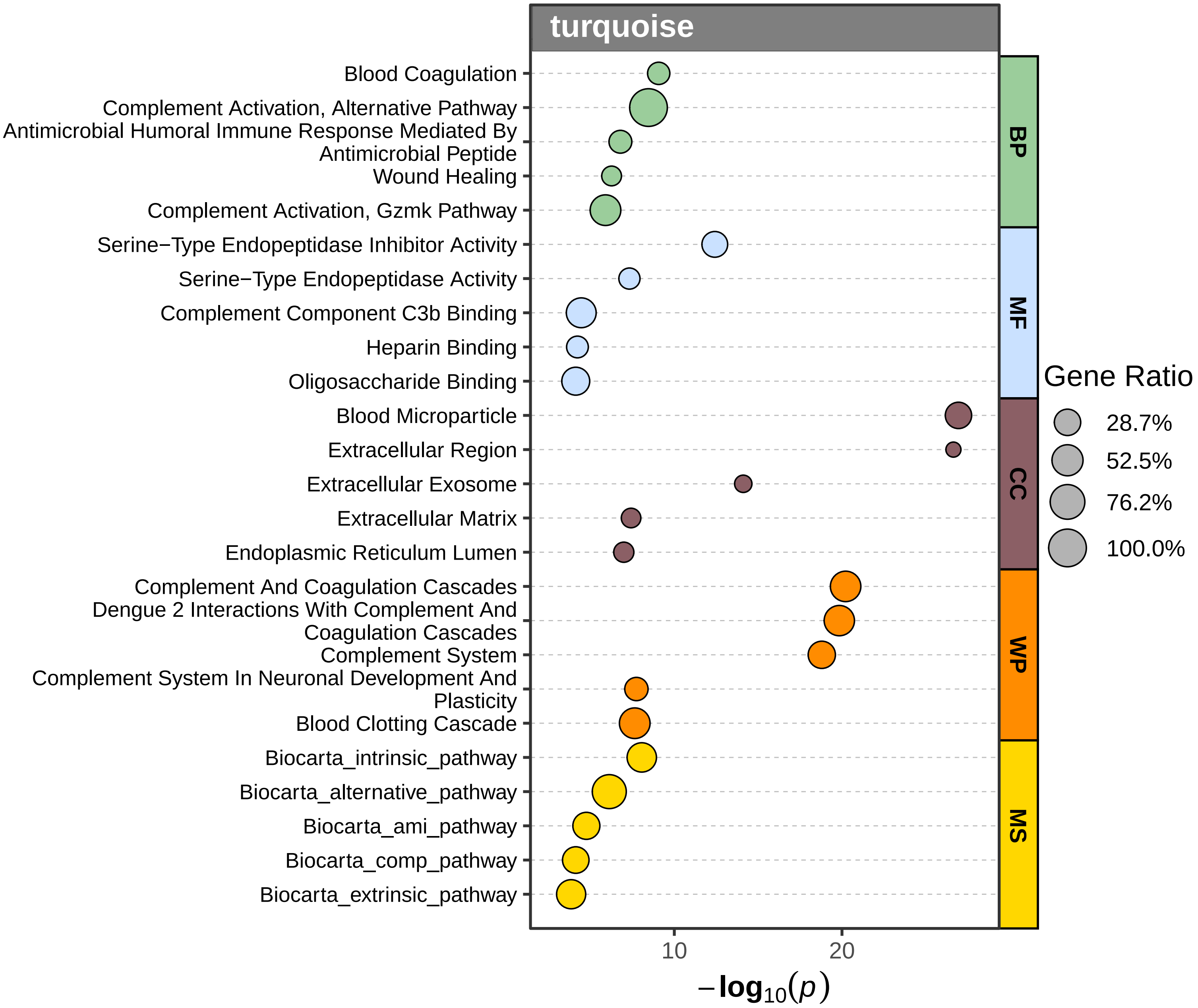

turquoise
Blood Coagulation
Complement Activation, Alternative Pathway
Antimicrobial Humoral Immune Response Mediated By Antimicrobial Peptide
Wound Healing
Complement Activation, Gzmk Pathway
Serine−Type Endopeptidase Inhibitor Activity
Serine−Type Endopeptidase Activity
Complement Component C3b Binding
Heparin Binding
Oligosaccharide Binding
Blood Microparticle
Extracellular Region
Extracellular Exosome
Extracellular Matrix
Endoplasmic Reticulum Lumen
Complement And Coagulation Cascades
Dengue 2 Interactions With Complement And Coagulation Cascades
Complement System
Complement System In Neuronal Development And Plasticity
Blood Clotting Cascade
Biocarta_intrinsic_pathway
Biocarta_alternative_pathway
Biocarta_ami_pathway
Biocarta_comp_pathway
Biocarta_extrinsic_pathway
BP
MF
CC
WP
MS
Gene Ratio
28.7%
52.5%
76.2%
100.0%
10
20
$-\log_{10}(p)$

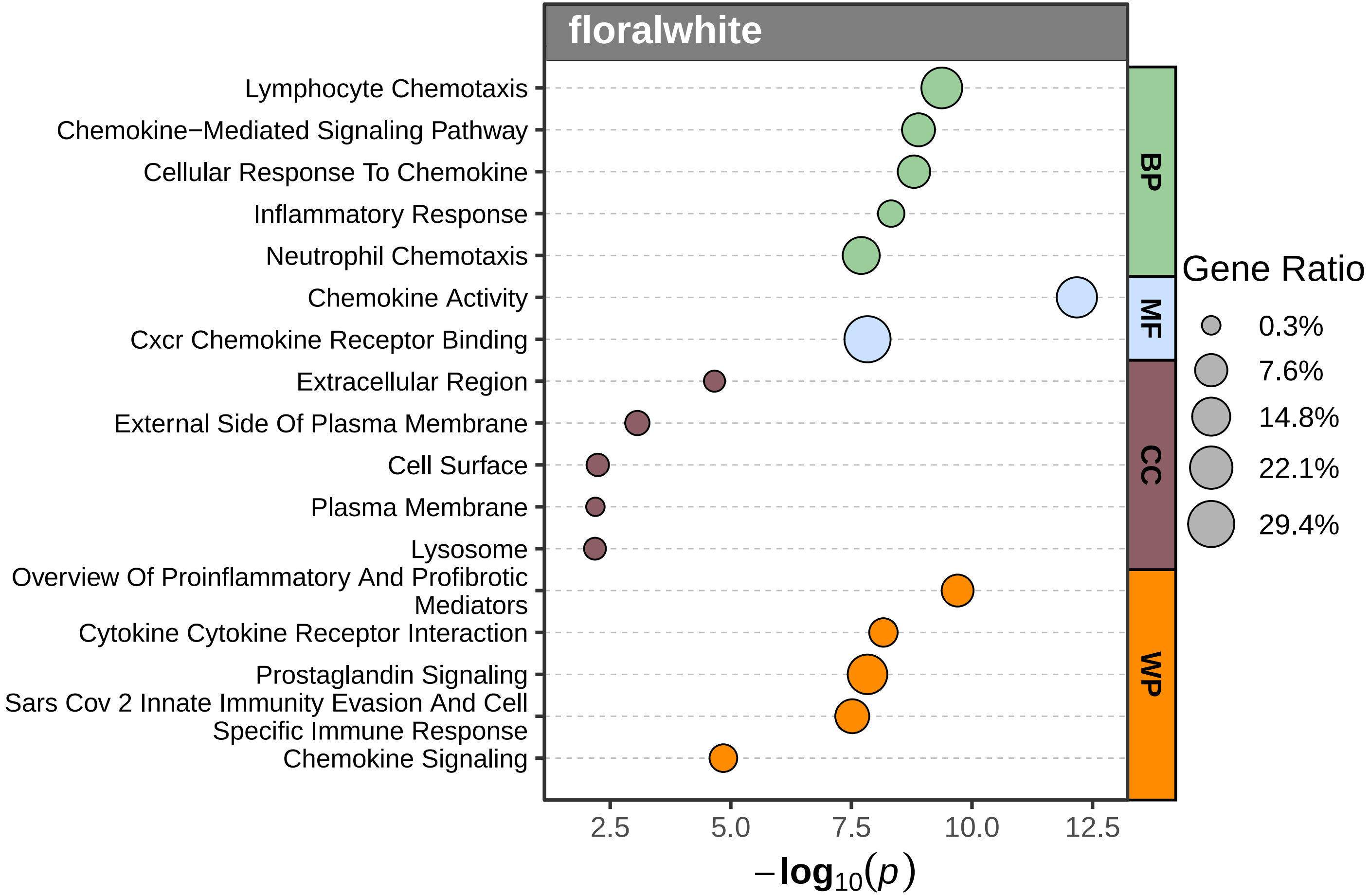

floralwhite
Lymphocyte Chemotaxis
Chemokine−Mediated Signaling Pathway
Cellular Response To Chemokine
Inflammatory Response
Neutrophil Chemotaxis
Chemokine Activity
Cxcr Chemokine Receptor Binding
Extracellular Region
External Side Of Plasma Membrane
Cell Surface
Plasma Membrane
Lysosome
Overview Of Proinflammatory And Profibrotic Mediators
Cytokine Cytokine Receptor Interaction
Prostaglandin Signaling
Sars Cov 2 Innate Immunity Evasion And Cell Specific Immune Response
Chemokine Signaling
BP
MF
CC
WP
Gene Ratio
0.3%
7.6%
14.8%
22.1%
29.4%
2.5
5.0
7.5
10.0
12.5
$-\log_{10}(p)$

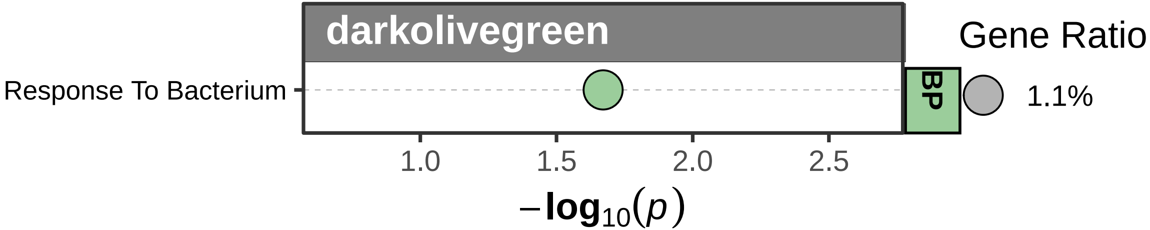

darkolivegreen
Gene Ratio
Response To Bacterium
BP
1.1%
1.0
1.5
2.0
2.5
$-\log_{10}(p)$

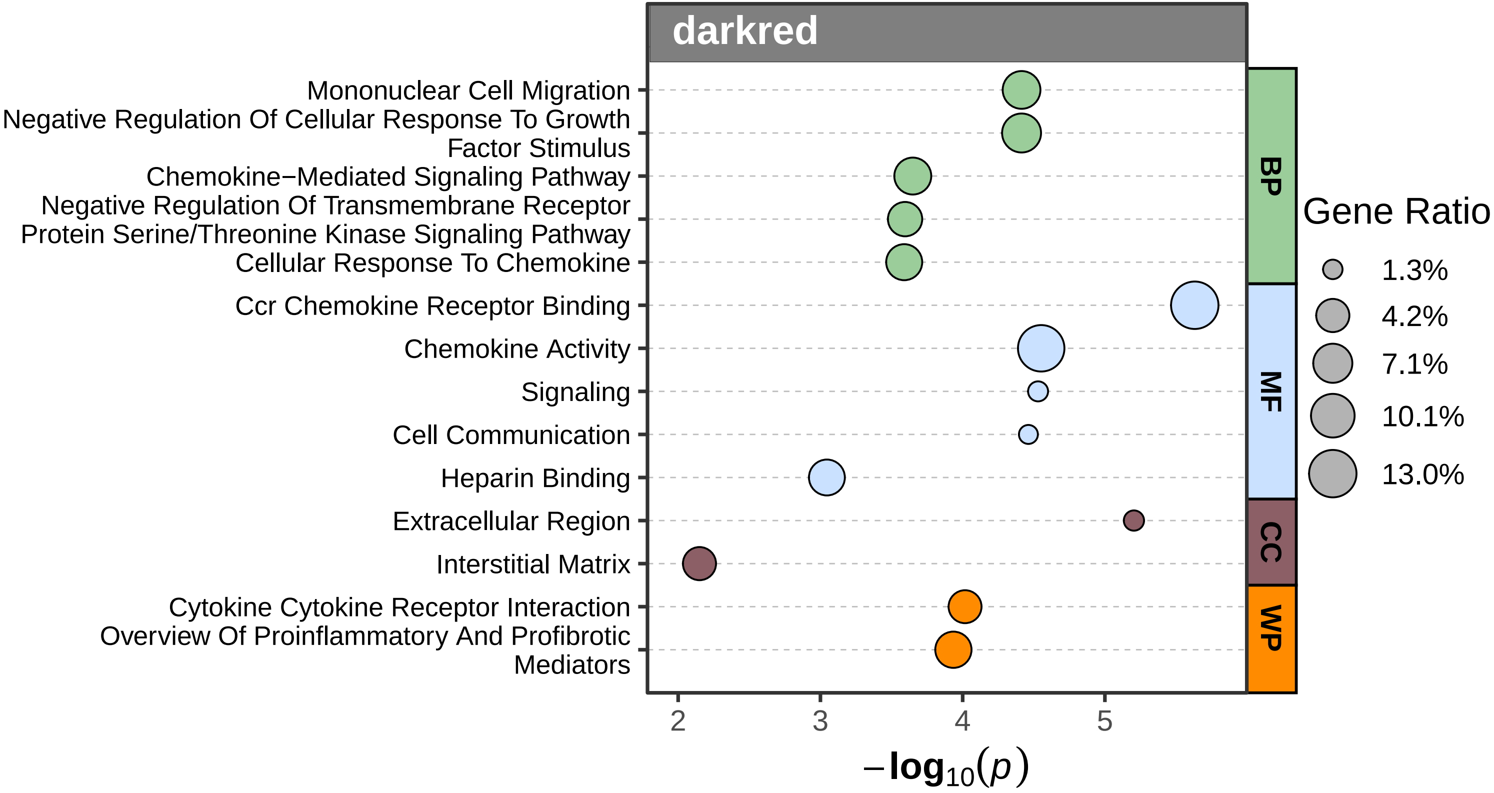

darkred
Mononuclear Cell Migration
Negative Regulation Of Cellular Response To Growth Factor Stimulus
Chemokine−Mediated Signaling Pathway
Negative Regulation Of Transmembrane Receptor Protein Serine/Threonine Kinase Signaling Pathway
Cellular Response To Chemokine
Ccr Chemokine Receptor Binding
Chemokine Activity
Signaling
Cell Communication
Heparin Binding
Extracellular Region
Interstitial Matrix
Cytokine Cytokine Receptor Interaction
Overview Of Proinflammatory And Profibrotic Mediators
BP
MF
CC
WP
2
3
4
5
−log10(p)
Gene Ratio
1.3%
4.2%
7.1%
10.1%
13.0%

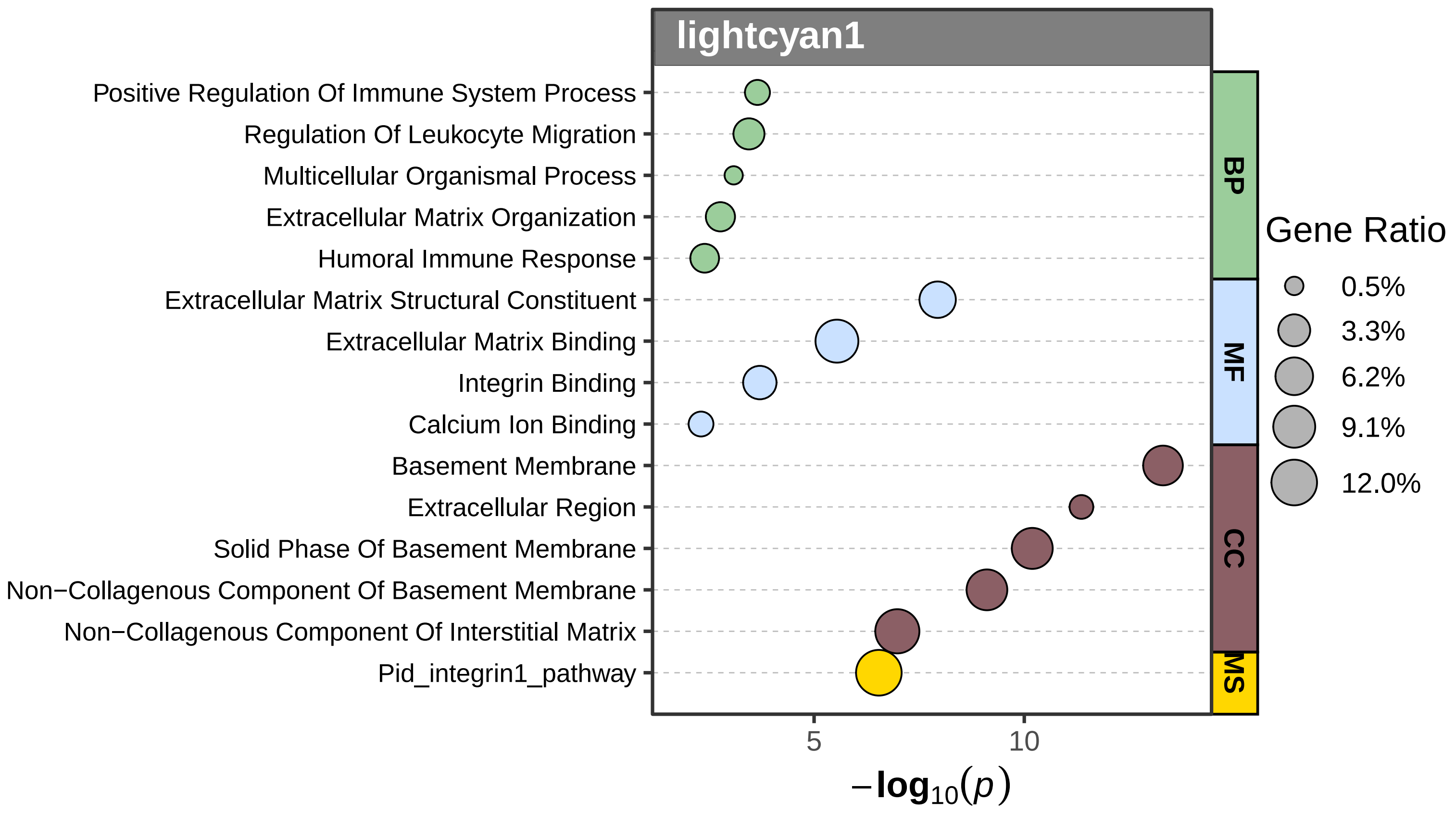
lightcyan1
Positive Regulation Of Immune System Process
Regulation Of Leukocyte Migration
Multicellular Organismal Process
Extracellular Matrix Organization
Humoral Immune Response
Extracellular Matrix Structural Constituent
Extracellular Matrix Binding
Integrin Binding
Calcium Ion Binding
Basement Membrane
Extracellular Region
Solid Phase Of Basement Membrane
Non−Collagenous Component Of Basement Membrane
Non−Collagenous Component Of Interstitial Matrix
Pid_integrin1_pathway
BP
MF
CC
MS
5
10
$-\log_{10}(p)$
Gene Ratio
0.5%
3.3%
6.2%
9.1%
12.0%

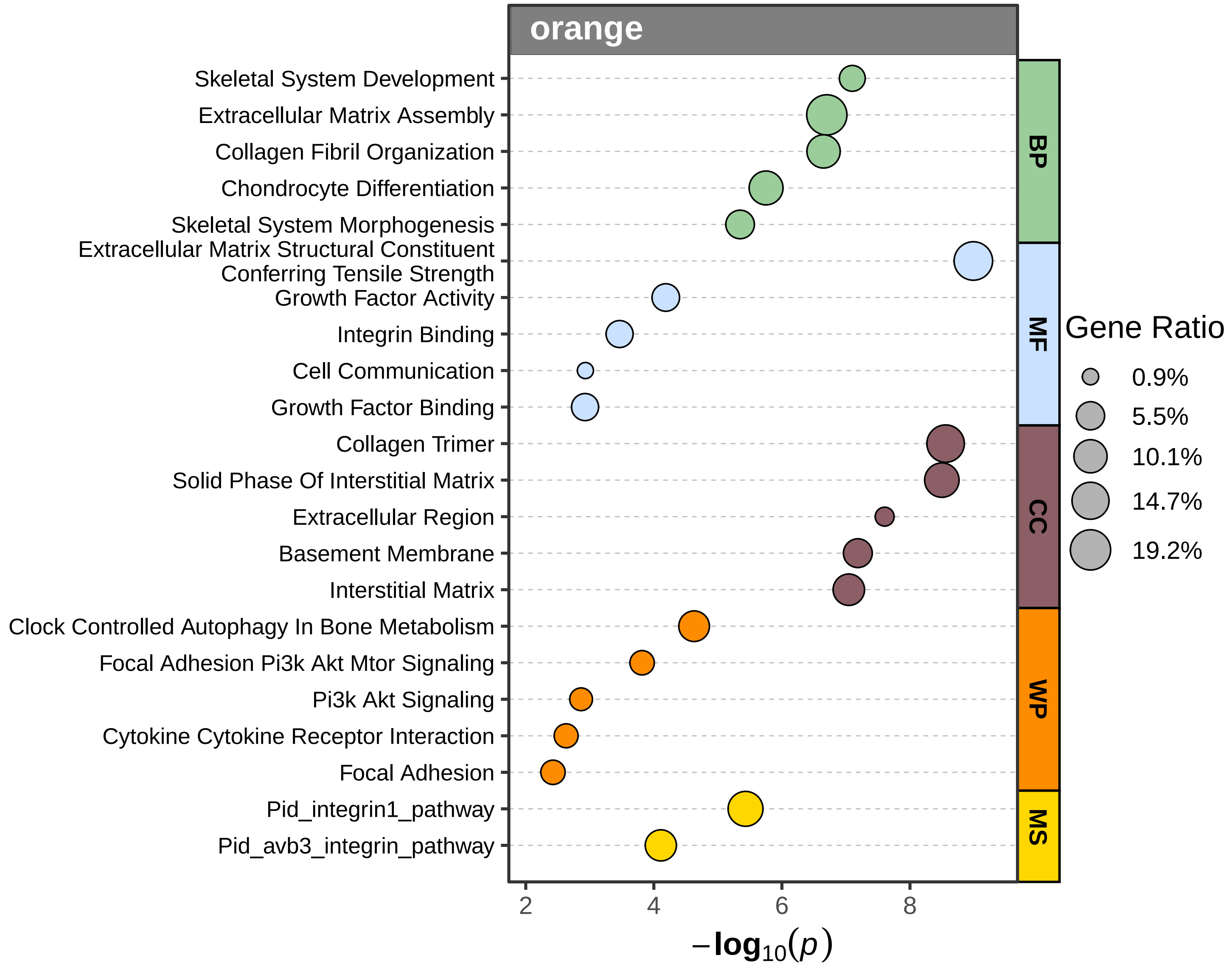
orange
Skeletal System Development
Extracellular Matrix Assembly
Collagen Fibril Organization
Chondrocyte Differentiation
Skeletal System Morphogenesis
Extracellular Matrix Structural Constituent Conferring Tensile Strength
Growth Factor Activity
Integrin Binding
Cell Communication
Growth Factor Binding
Collagen Trimer
Solid Phase Of Interstitial Matrix
Extracellular Region
Basement Membrane
Interstitial Matrix
Clock Controlled Autophagy In Bone Metabolism
Focal Adhesion Pi3k Akt Mtor Signaling
Pi3k Akt Signaling
Cytokine Cytokine Receptor Interaction
Focal Adhesion
Pid_integrin1_pathway
Pid_avb3_integrin_pathway
BP
MF
CC
WP
MS
Gene Ratio
0.9%
5.5%
10.1%
14.7%
19.2%
2
4
6
8
$-\log_{10}(p)$

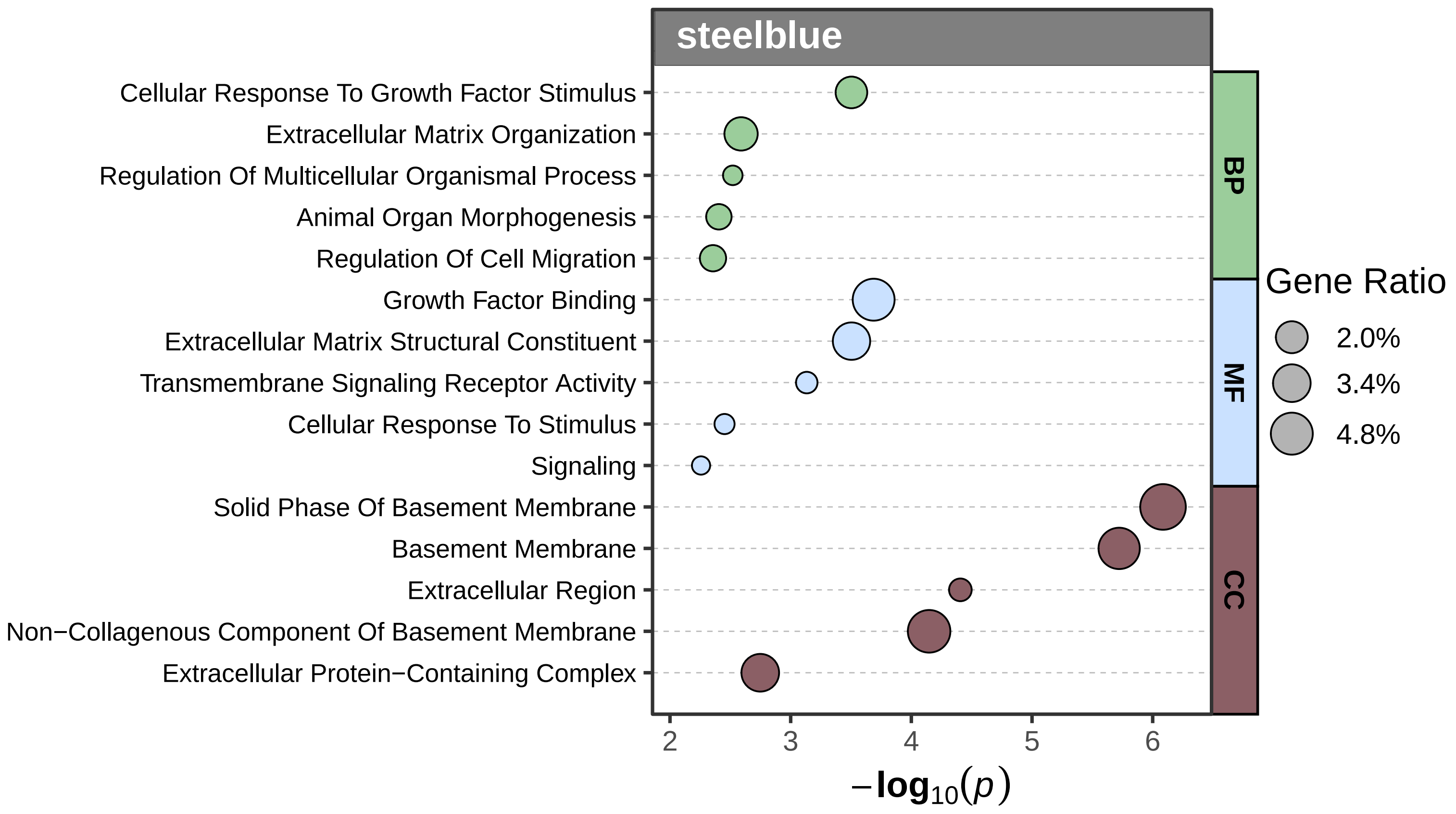

steelblue
Cellular Response To Growth Factor Stimulus
Extracellular Matrix Organization
Regulation Of Multicellular Organismal Process
Animal Organ Morphogenesis
Regulation Of Cell Migration
Growth Factor Binding
Extracellular Matrix Structural Constituent
Transmembrane Signaling Receptor Activity
Cellular Response To Stimulus
Signaling
Solid Phase Of Basement Membrane
Basement Membrane
Extracellular Region
Non−Collagenous Component Of Basement Membrane
Extracellular Protein−Containing Complex
BP
MF
CC
Gene Ratio
2.0%
3.4%
4.8%
2
3
4
5
6
$-\log_{10}(p)$

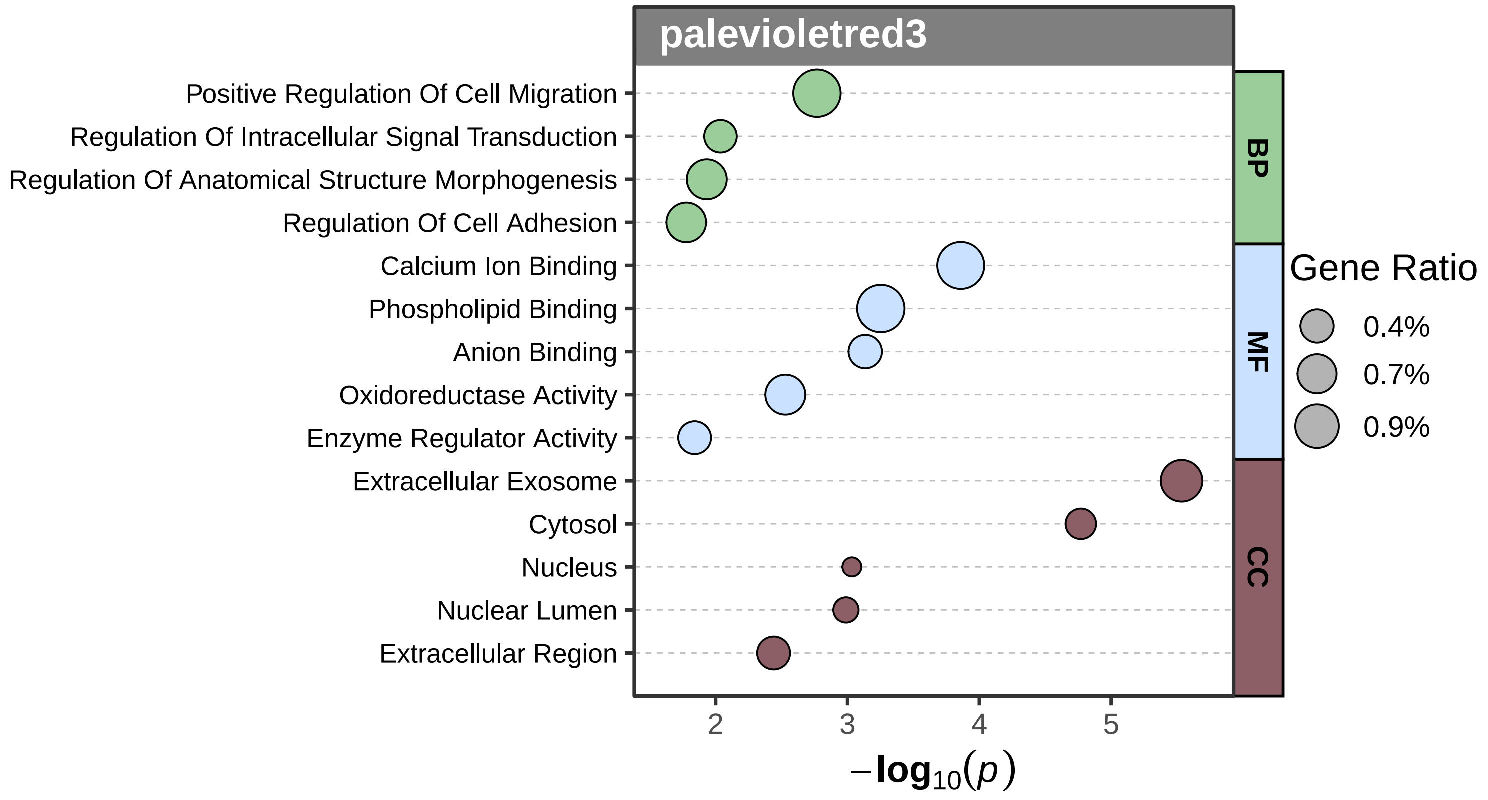
palevioletred3
Positive Regulation Of Cell Migration
Regulation Of Intracellular Signal Transduction
Regulation Of Anatomical Structure Morphogenesis
Regulation Of Cell Adhesion
Calcium Ion Binding
Phospholipid Binding
Anion Binding
Oxidoreductase Activity
Enzyme Regulator Activity
Extracellular Exosome
Cytosol
Nucleus
Nuclear Lumen
Extracellular Region
BP
MF
CC
Gene Ratio
0.4%
0.7%
0.9%
2
3
4
5
$-\log_{10}(p)$

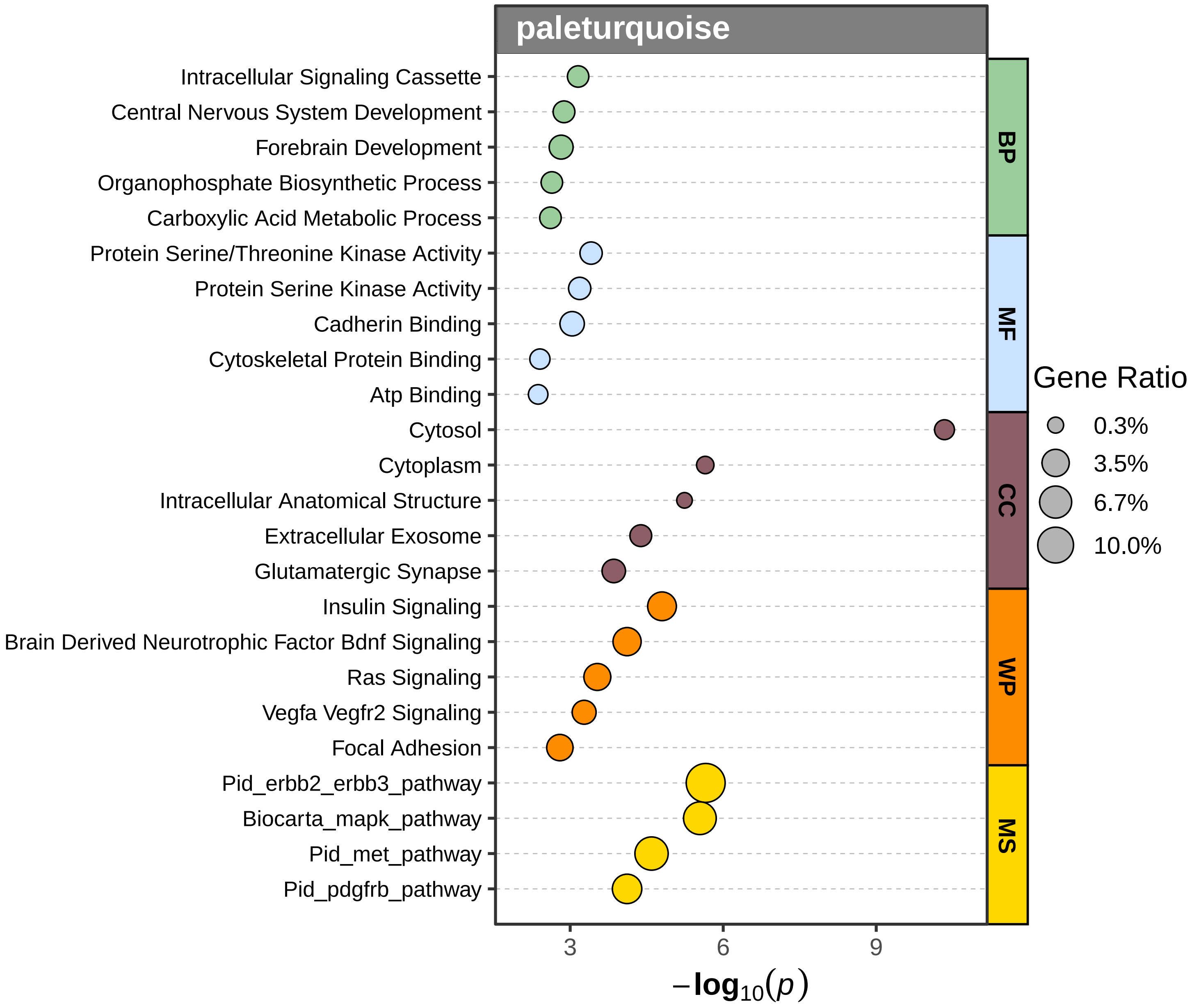

paleturquoise
Intracellular Signaling Cassette
Central Nervous System Development
Forebrain Development
Organophosphate Biosynthetic Process
Carboxylic Acid Metabolic Process
Protein Serine/Threonine Kinase Activity
Protein Serine Kinase Activity
Cadherin Binding
Cytoskeletal Protein Binding
Atp Binding
Cytosol
Cytoplasm
Intracellular Anatomical Structure
Extracellular Exosome
Glutamatergic Synapse
Insulin Signaling
Brain Derived Neurotrophic Factor Bdnf Signaling
Ras Signaling
Vegfa Vegfr2 Signaling
Focal Adhesion
Pid_erbb2_erbb3_pathway
Biocarta_mapk_pathway
Pid_met_pathway
Pid_pdgfrb_pathway
BP
MF
CC
WP
MS
Gene Ratio
0.3%
3.5%
6.7%
10.0%
3
6
9
$-\log_{10}(p)$

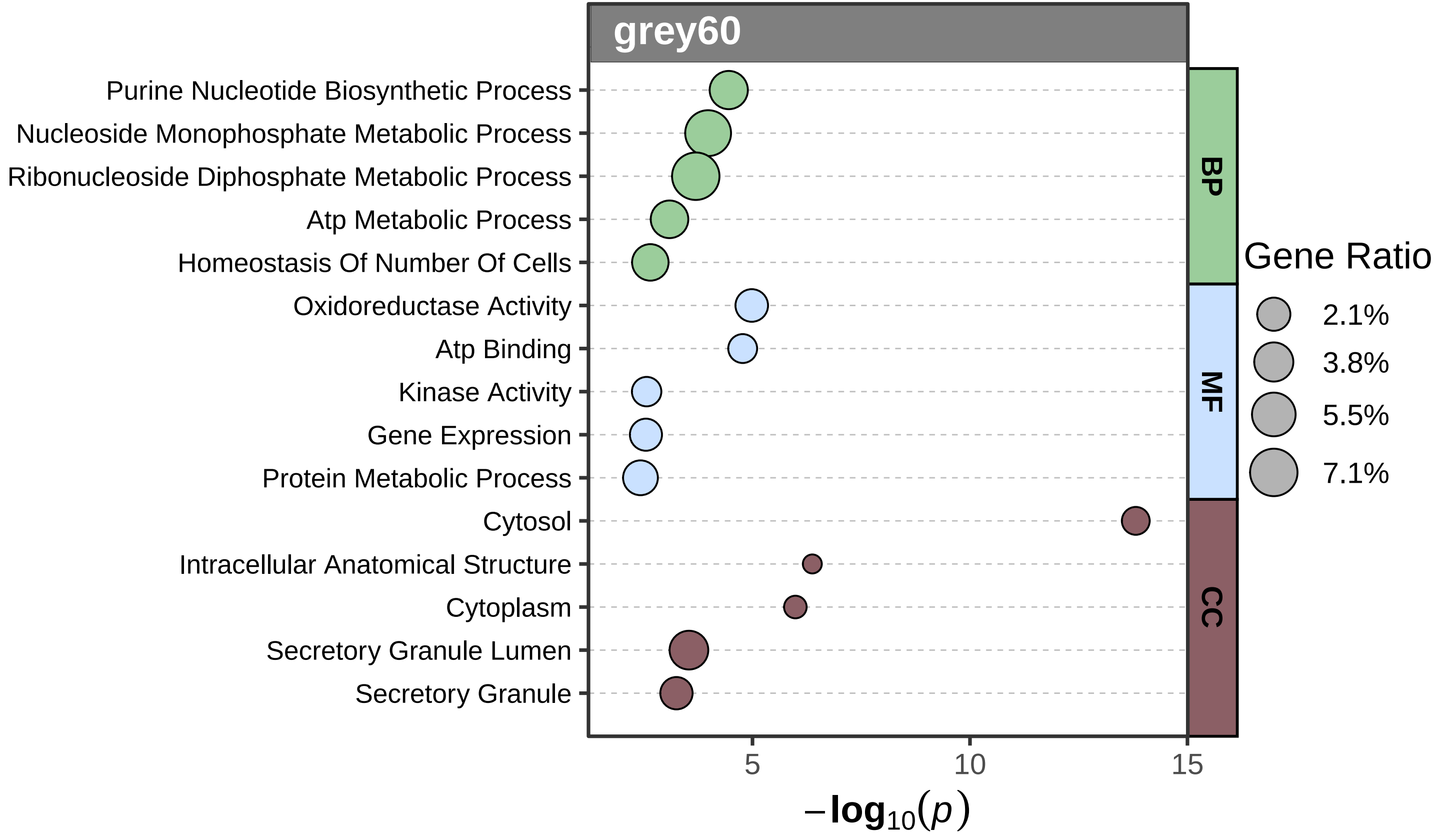

grey60
Purine Nucleotide Biosynthetic Process
Nucleoside Monophosphate Metabolic Process
Ribonucleoside Diphosphate Metabolic Process
Atp Metabolic Process
Homeostasis Of Number Of Cells
Oxidoreductase Activity
Atp Binding
Kinase Activity
Gene Expression
Protein Metabolic Process
Cytosol
Intracellular Anatomical Structure
Cytoplasm
Secretory Granule Lumen
Secretory Granule
BP
MF
CC
Gene Ratio
2.1%
3.8%
5.5%
7.1%
5
10
15
$-\log_{10}(p)$

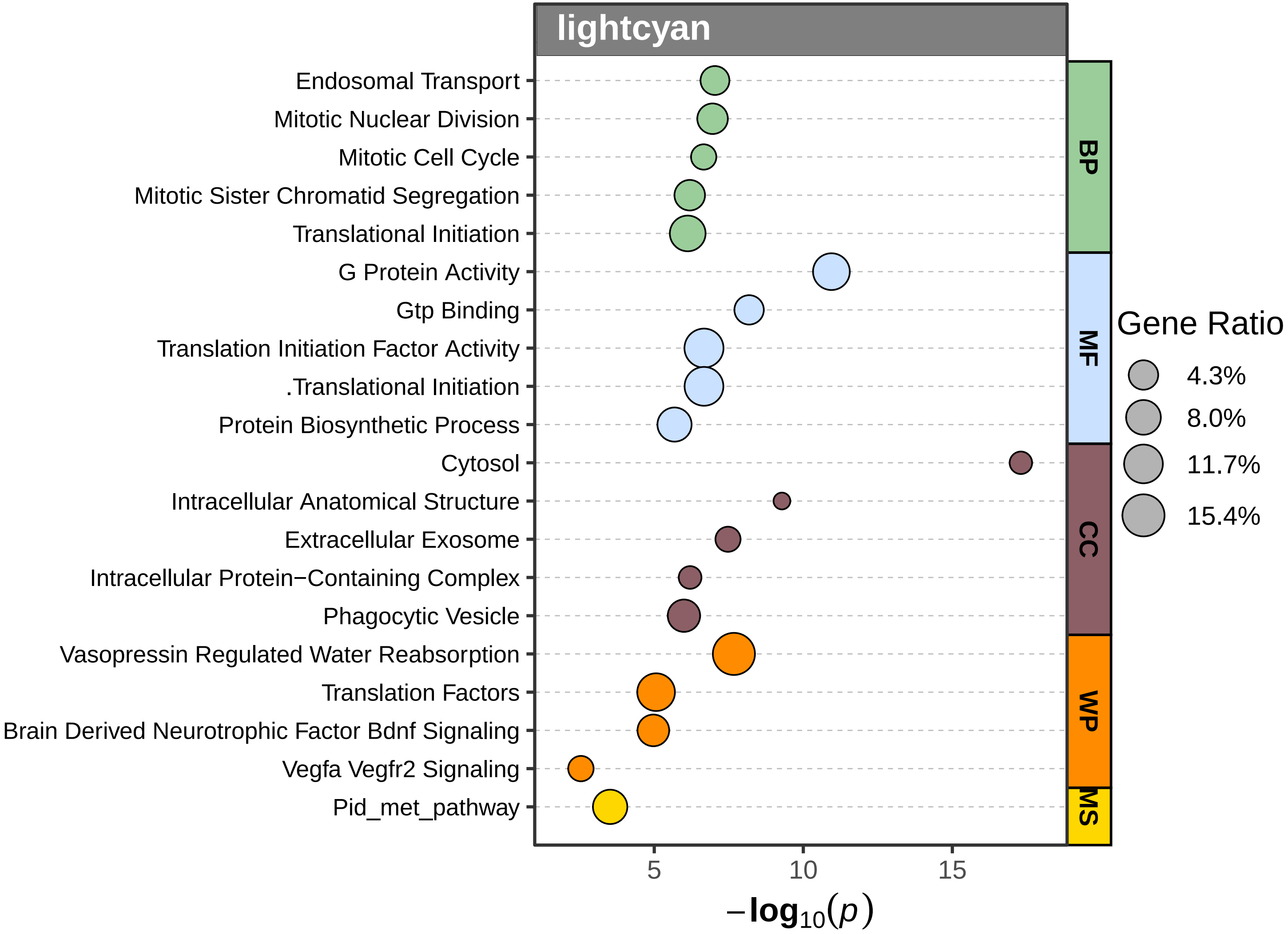

lightcyan
Endosomal Transport
Mitotic Nuclear Division
Mitotic Cell Cycle
Mitotic Sister Chromatid Segregation
Translational Initiation
G Protein Activity
Gtp Binding
Translation Initiation Factor Activity
.Translational Initiation
Protein Biosynthetic Process
Cytosol
Intracellular Anatomical Structure
Extracellular Exosome
Intracellular Protein−Containing Complex
Phagocytic Vesicle
Vasopressin Regulated Water Reabsorption
Translation Factors
Brain Derived Neurotrophic Factor Bdnf Signaling
Vegfa Vegfr2 Signaling
Pid_met_pathway
BP
MF
CC
WP
MS
5
10
15
$-\log_{10}(p)$
Gene Ratio
4.3%
8.0%
11.7%
15.4%

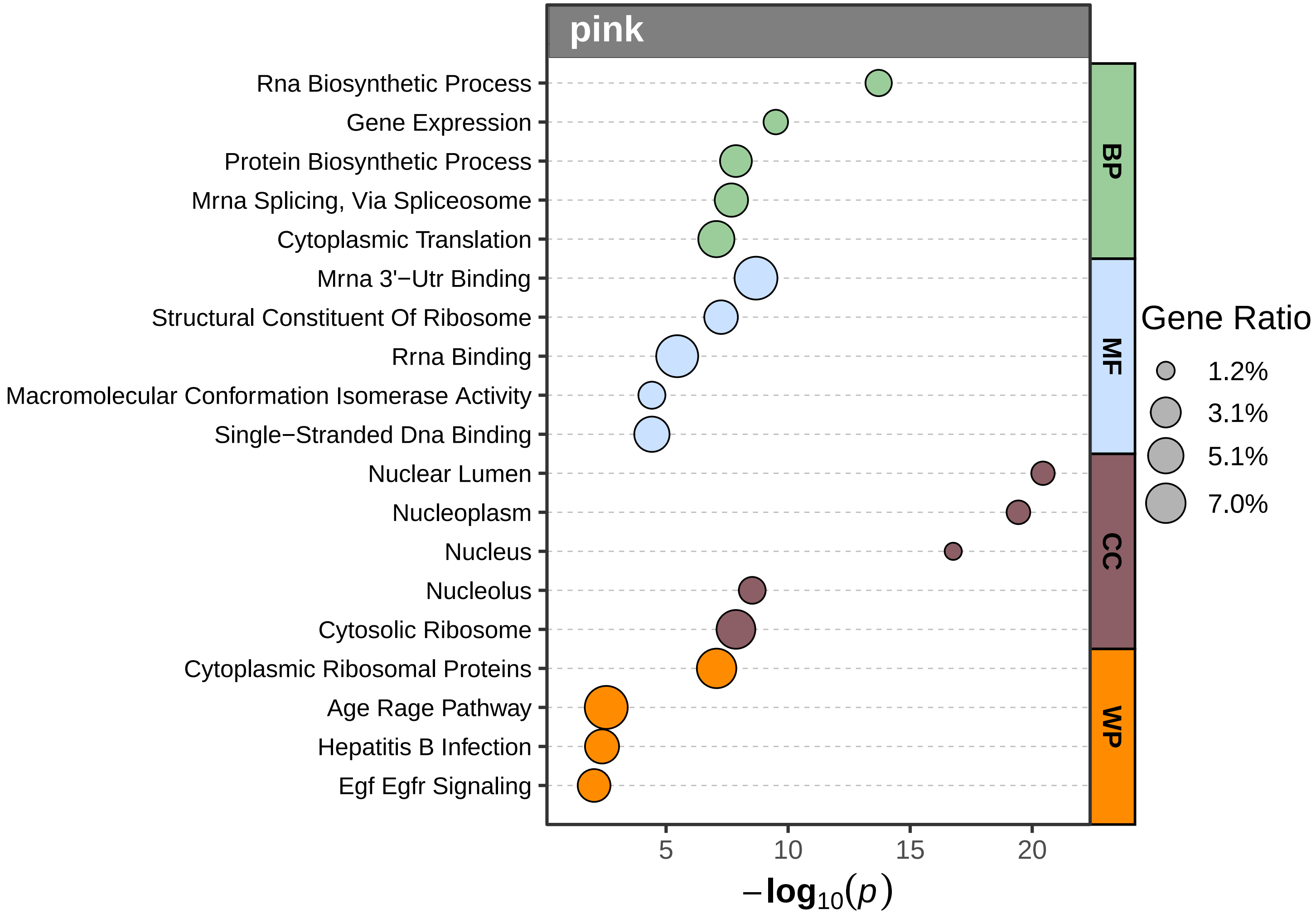

pink
Rna Biosynthetic Process
Gene Expression
Protein Biosynthetic Process
Mrna Splicing, Via Spliceosome
Cytoplasmic Translation
Mrna 3'−Utr Binding
Structural Constituent Of Ribosome
Rrna Binding
Macromolecular Conformation Isomerase Activity
Single−Stranded Dna Binding
Nuclear Lumen
Nucleoplasm
Nucleus
Nucleolus
Cytosolic Ribosome
Cytoplasmic Ribosomal Proteins
Age Rage Pathway
Hepatitis B Infection
Egf Egfr Signaling
5
10
15
20
$-\log_{10}(p)$
BP
MF
CC
WP
Gene Ratio
1.2%
3.1%
5.1%
7.0%

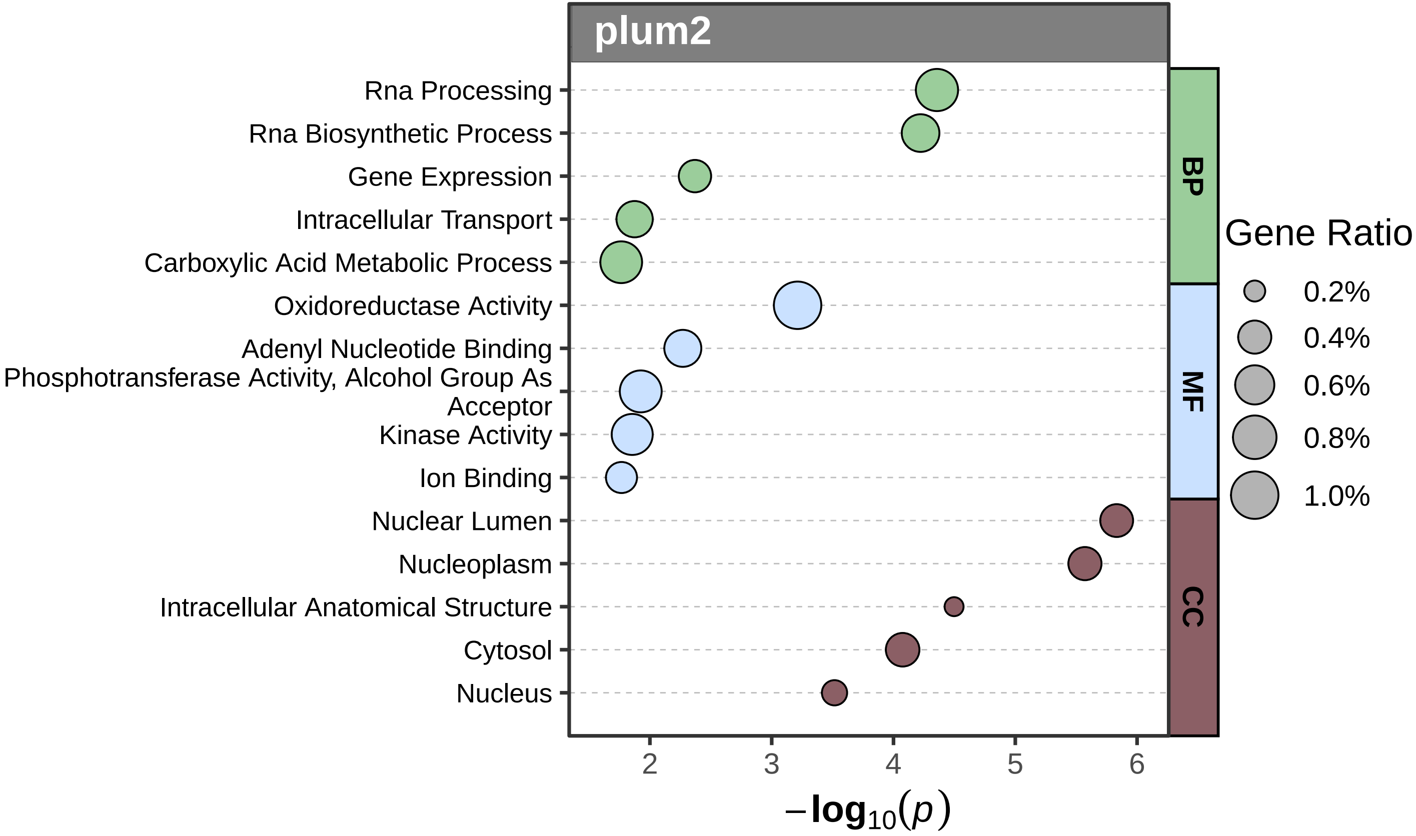
plum2
Rna Processing
Rna Biosynthetic Process
Gene Expression
Intracellular Transport
Carboxylic Acid Metabolic Process
Oxidoreductase Activity
Adenyl Nucleotide Binding
Phosphotransferase Activity, Alcohol Group As Acceptor
Kinase Activity
Ion Binding
Nuclear Lumen
Nucleoplasm
Intracellular Anatomical Structure
Cytosol
Nucleus
BP
MF
CC
Gene Ratio
0.2%
0.4%
0.6%
0.8%
1.0%
2
3
4
5
6
$-\log_{10}(p)$

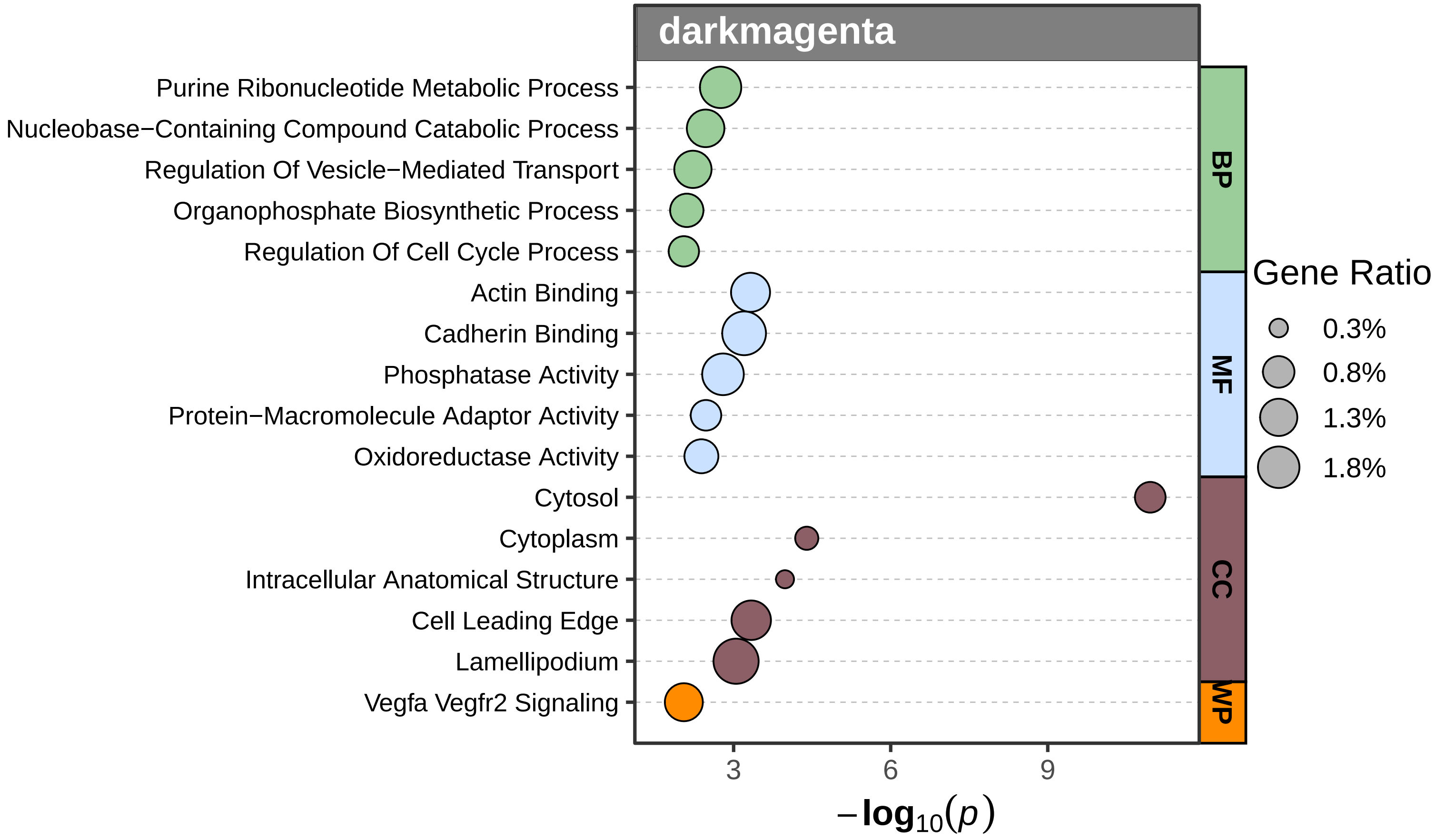
darkmagenta
Purine Ribonucleotide Metabolic Process
Nucleobase−Containing Compound Catabolic Process
Regulation Of Vesicle−Mediated Transport
Organophosphate Biosynthetic Process
Regulation Of Cell Cycle Process
Actin Binding
Cadherin Binding
Phosphatase Activity
Protein−Macromolecule Adaptor Activity
Oxidoreductase Activity
Cytosol
Cytoplasm
Intracellular Anatomical Structure
Cell Leading Edge
Lamellipodium
Vegfa Vegfr2 Signaling
BP
MF
CC
WP
Gene Ratio
0.3%
0.8%
1.3%
1.8%
3
6
9
−log10(p)

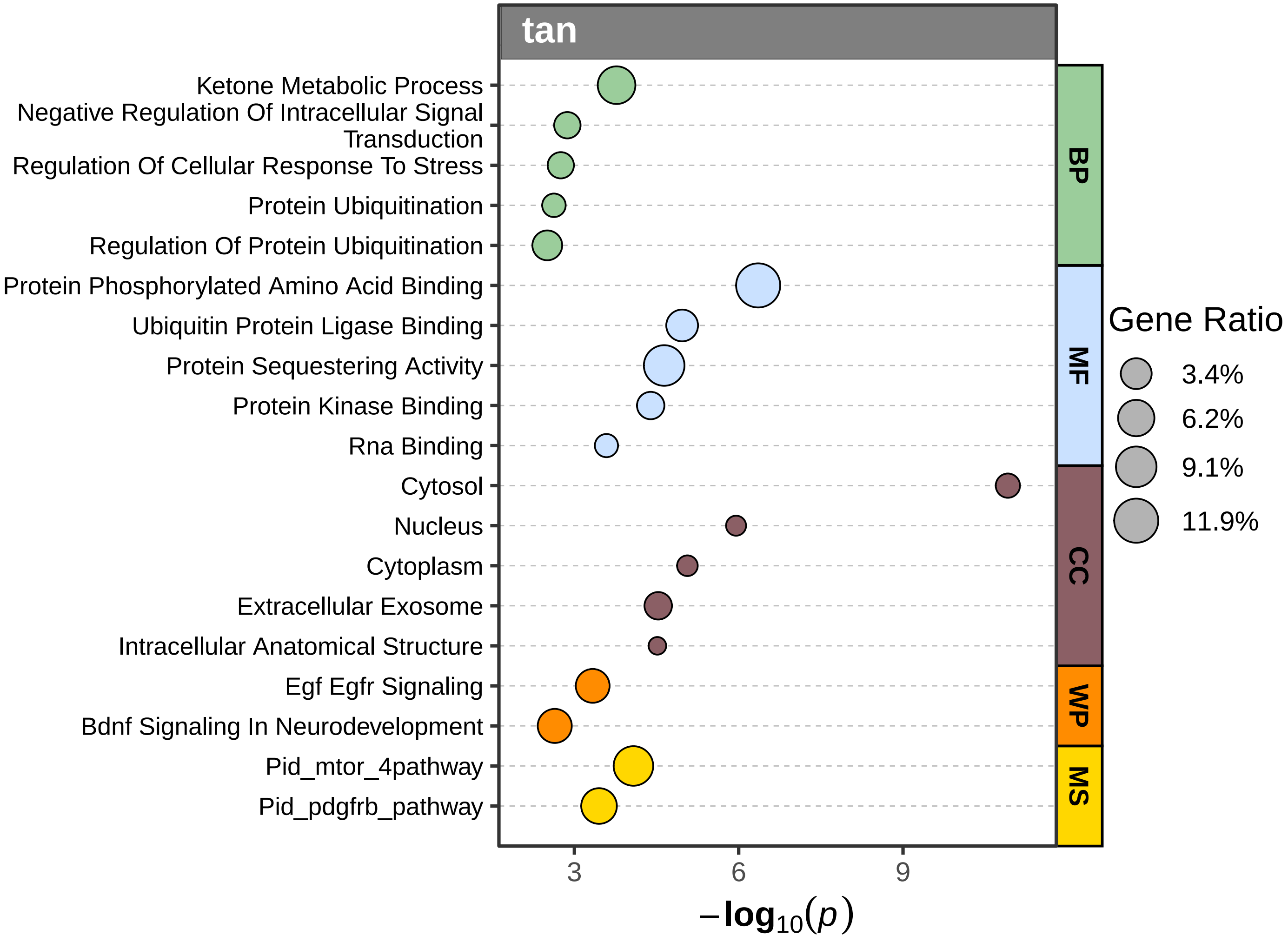

tan
Ketone Metabolic Process
Negative Regulation Of Intracellular Signal Transduction
Regulation Of Cellular Response To Stress
Protein Ubiquitination
Regulation Of Protein Ubiquitination
Protein Phosphorylated Amino Acid Binding
Ubiquitin Protein Ligase Binding
Protein Sequestering Activity
Protein Kinase Binding
Rna Binding
Cytosol
Nucleus
Cytoplasm
Extracellular Exosome
Intracellular Anatomical Structure
Egf Egfr Signaling
Bdnf Signaling In Neurodevelopment
Pid_mtor_4pathway
Pid_pdgfrb_pathway
BP
MF
CC
WP
MS
3
6
9
$-\log_{10}(p)$
Gene Ratio
3.4%
6.2%
9.1%
11.9%

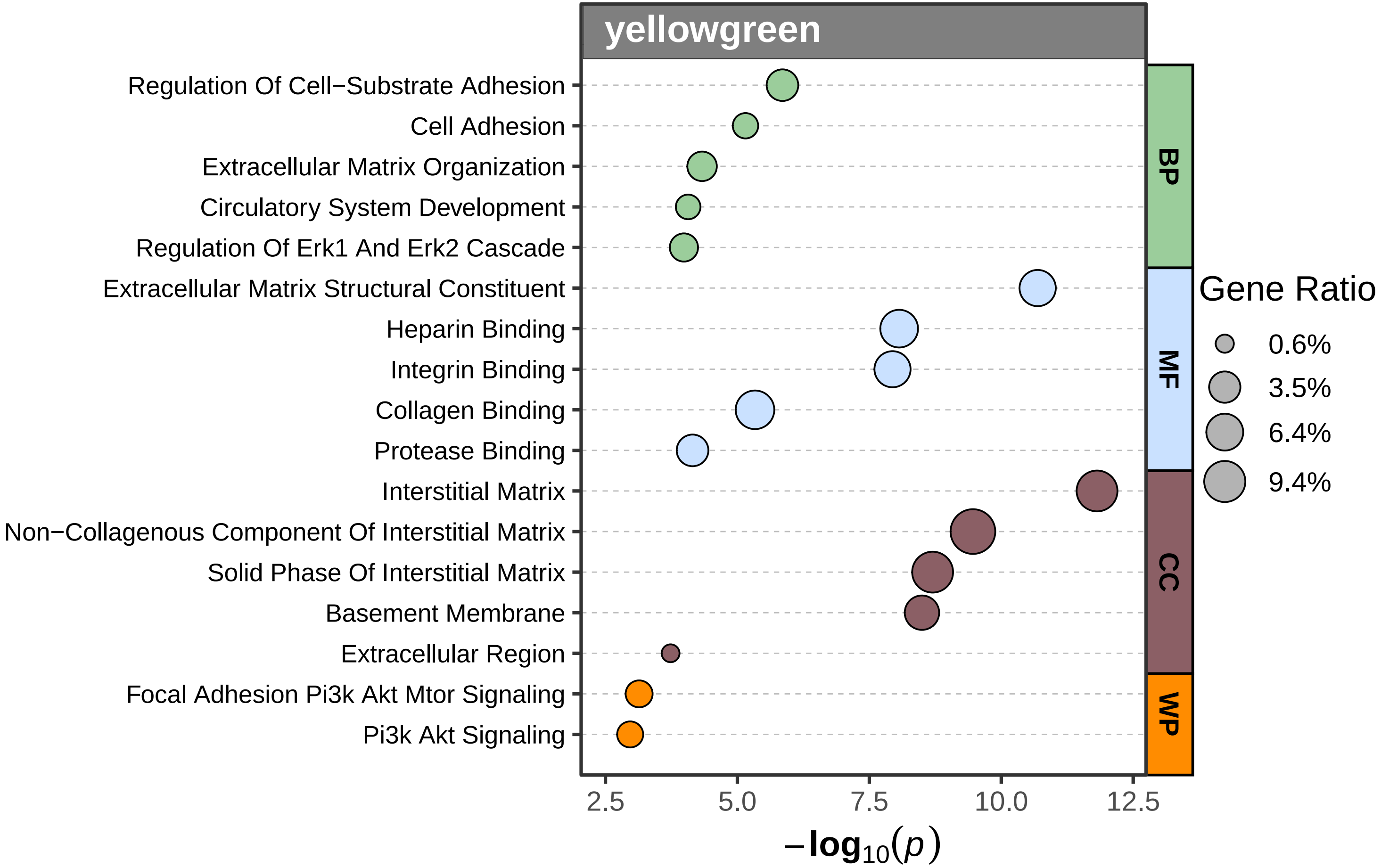
yellowgreen
Regulation Of Cell−Substrate Adhesion
Cell Adhesion
Extracellular Matrix Organization
Circulatory System Development
Regulation Of Erk1 And Erk2 Cascade
Extracellular Matrix Structural Constituent
Heparin Binding
Integrin Binding
Collagen Binding
Protease Binding
Interstitial Matrix
Non−Collagenous Component Of Interstitial Matrix
Solid Phase Of Interstitial Matrix
Basement Membrane
Extracellular Region
Focal Adhesion Pi3k Akt Mtor Signaling
Pi3k Akt Signaling
BP
MF
CC
WP
Gene Ratio
0.6%
3.5%
6.4%
9.4%
2.5
5.0
7.5
10.0
12.5
$-\log_{10}(p)$

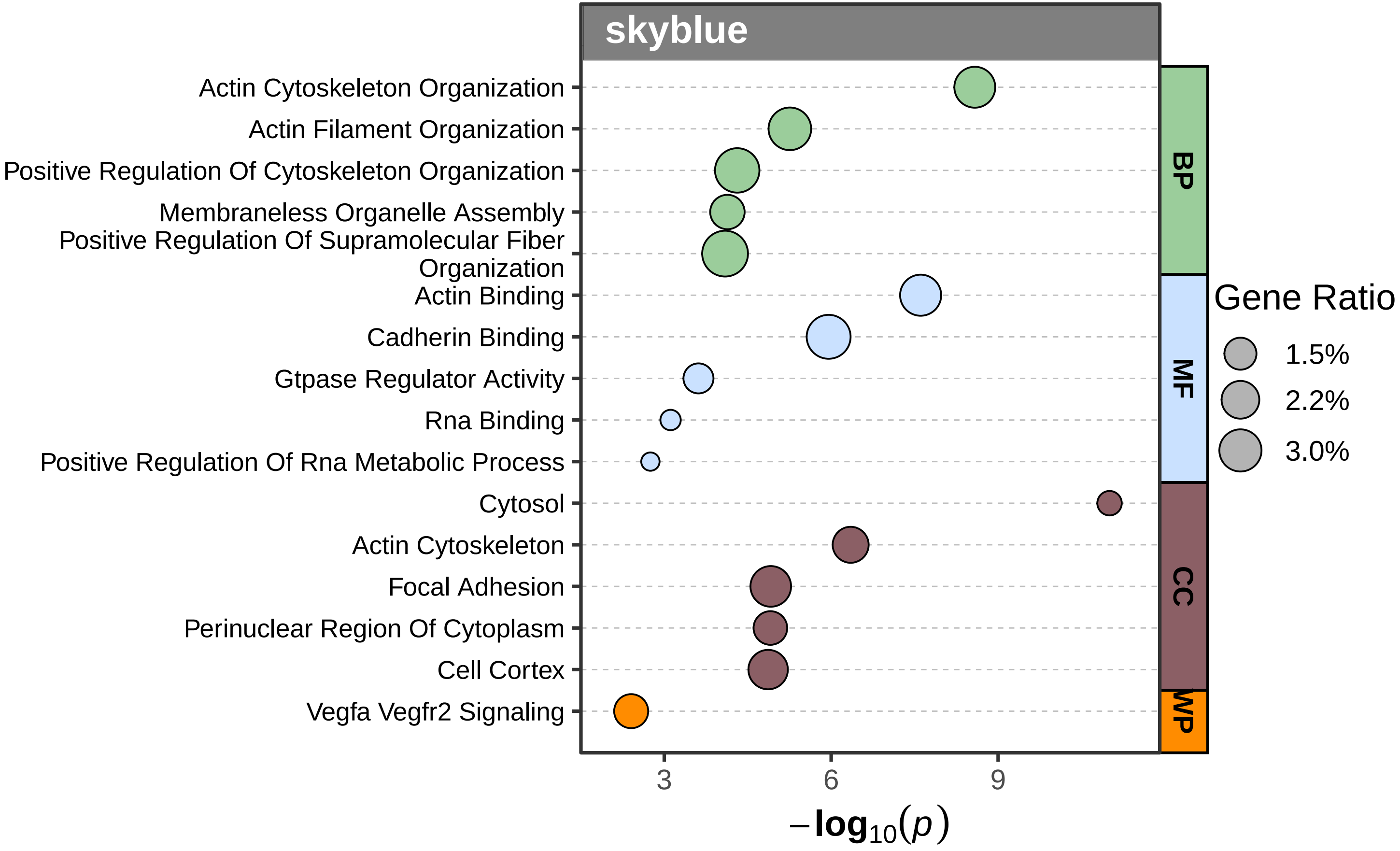
skyblue
Actin Cytoskeleton Organization
Actin Filament Organization
Positive Regulation Of Cytoskeleton Organization
Membraneless Organelle Assembly
Positive Regulation Of Supramolecular Fiber Organization
Actin Binding
Cadherin Binding
Gtpase Regulator Activity
Rna Binding
Positive Regulation Of Rna Metabolic Process
Cytosol
Actin Cytoskeleton
Focal Adhesion
Perinuclear Region Of Cytoplasm
Cell Cortex
Vegfa Vegfr2 Signaling
BP
MF
CC
WP
Gene Ratio
1.5%
2.2%
3.0%
3
6
9
$-\log_{10}(p)$

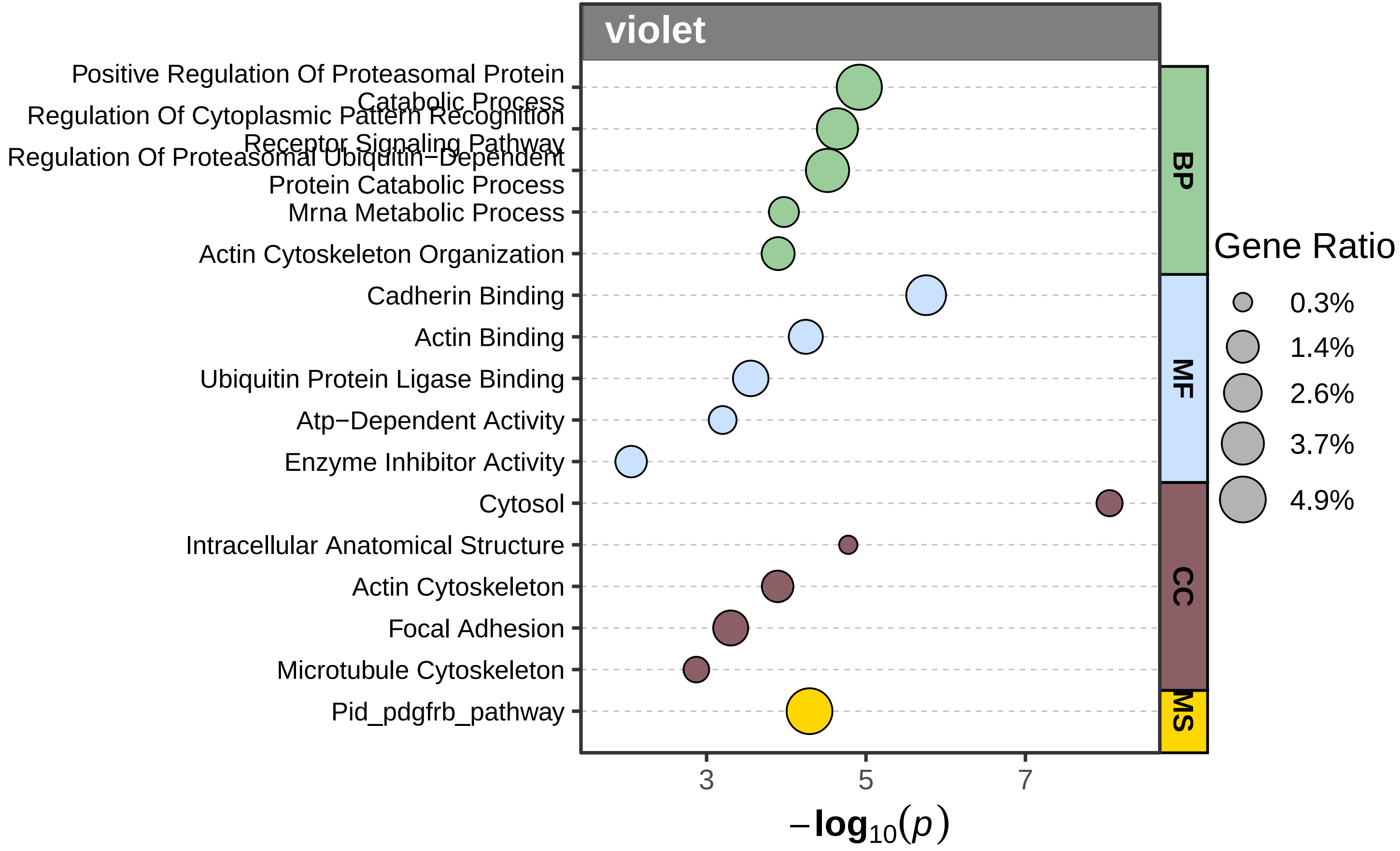
violet
Positive Regulation Of Proteasomal Protein Catabolic Process
Regulation Of Cytoplasmic Pattern Recognition Receptor Signaling Pathway
Regulation Of Proteasomal Ubiquitin−Dependent Protein Catabolic Process
Mrna Metabolic Process
Actin Cytoskeleton Organization
Cadherin Binding
Actin Binding
Ubiquitin Protein Ligase Binding
Atp−Dependent Activity
Enzyme Inhibitor Activity
Cytosol
Intracellular Anatomical Structure
Actin Cytoskeleton
Focal Adhesion
Microtubule Cytoskeleton
Pid_pdgfrb_pathway
BP
MF
CC
MS
Gene Ratio
0.3%
1.4%
2.6%
3.7%
4.9%
3
5
7
$-\log_{10}(p)$

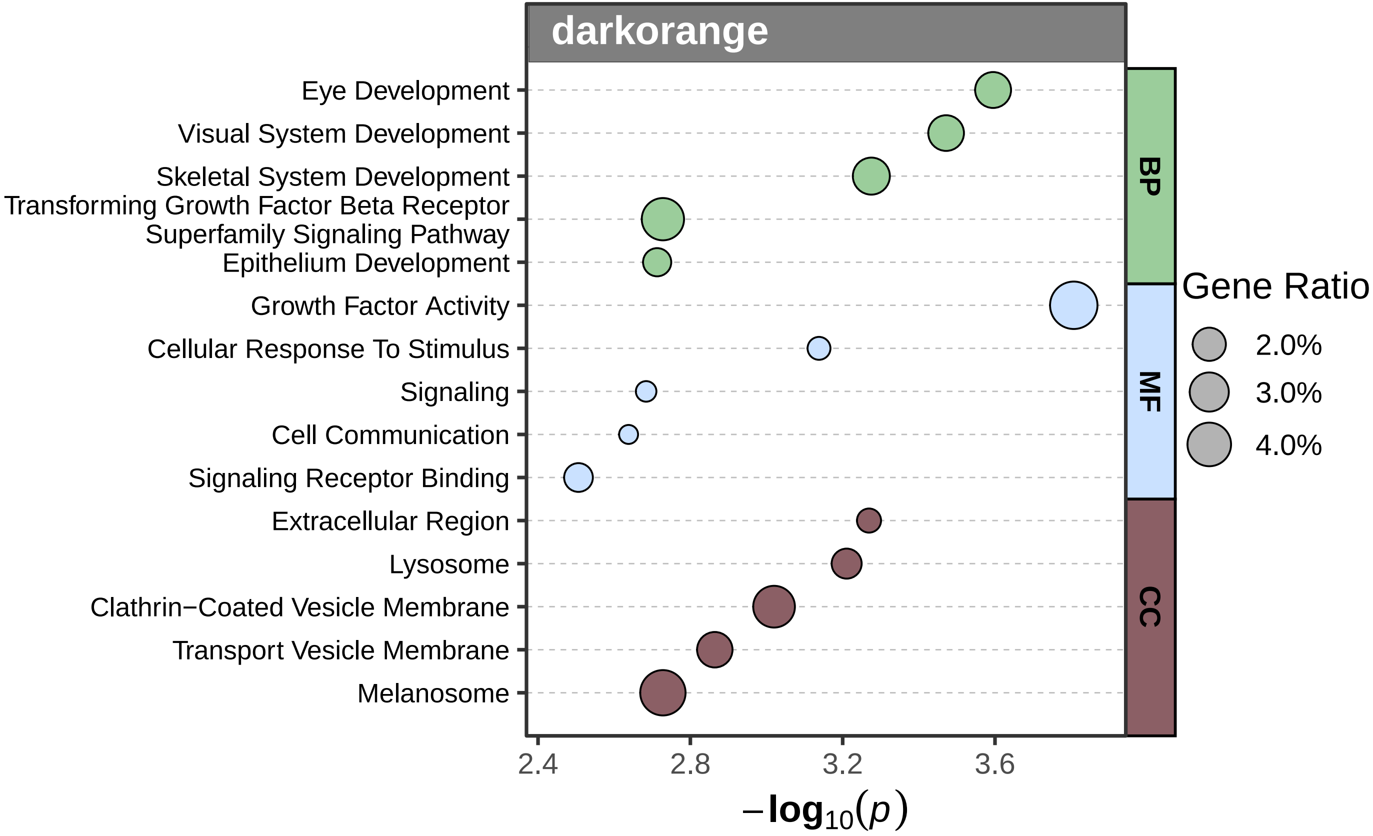
darkorange
Eye Development
Visual System Development
Skeletal System Development
Transforming Growth Factor Beta Receptor Superfamily Signaling Pathway
Epithelium Development
Growth Factor Activity
Cellular Response To Stimulus
Signaling
Cell Communication
Signaling Receptor Binding
Extracellular Region
Lysosome
Clathrin−Coated Vesicle Membrane
Transport Vesicle Membrane
Melanosome
BP
MF
CC
Gene Ratio
2.0%
3.0%
4.0%
2.4
2.8
3.2
3.6
$-\log_{10}(p)$

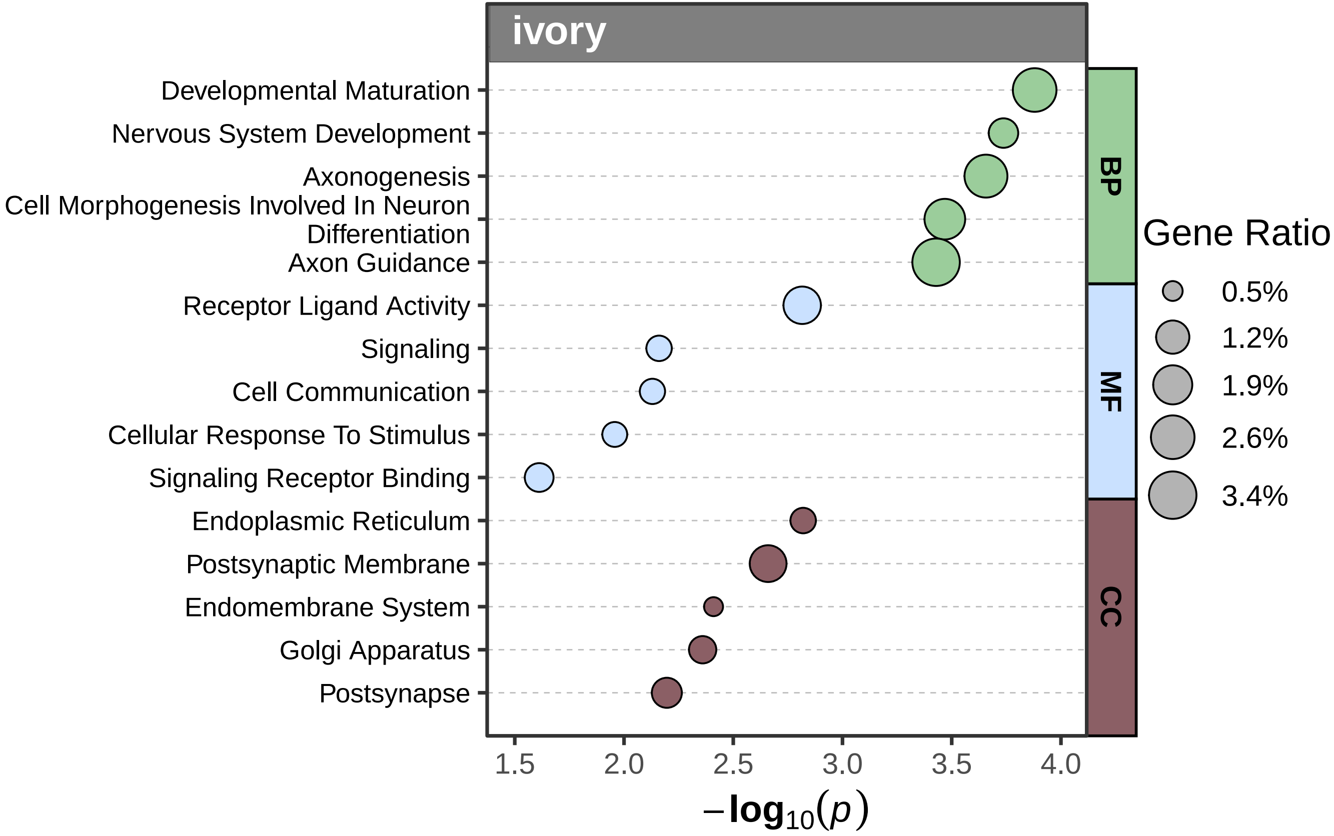
ivory
Developmental Maturation
Nervous System Development
Axonogenesis
Cell Morphogenesis Involved In Neuron Differentiation
Axon Guidance
Receptor Ligand Activity
Signaling
Cell Communication
Cellular Response To Stimulus
Signaling Receptor Binding
Endoplasmic Reticulum
Postsynaptic Membrane
Endomembrane System
Golgi Apparatus
Postsynapse
BP
MF
CC
Gene Ratio
0.5%
1.2%
1.9%
2.6%
3.4%
1.5
2.0
2.5
3.0
3.5
4.0
$-\log_{10}(p)$

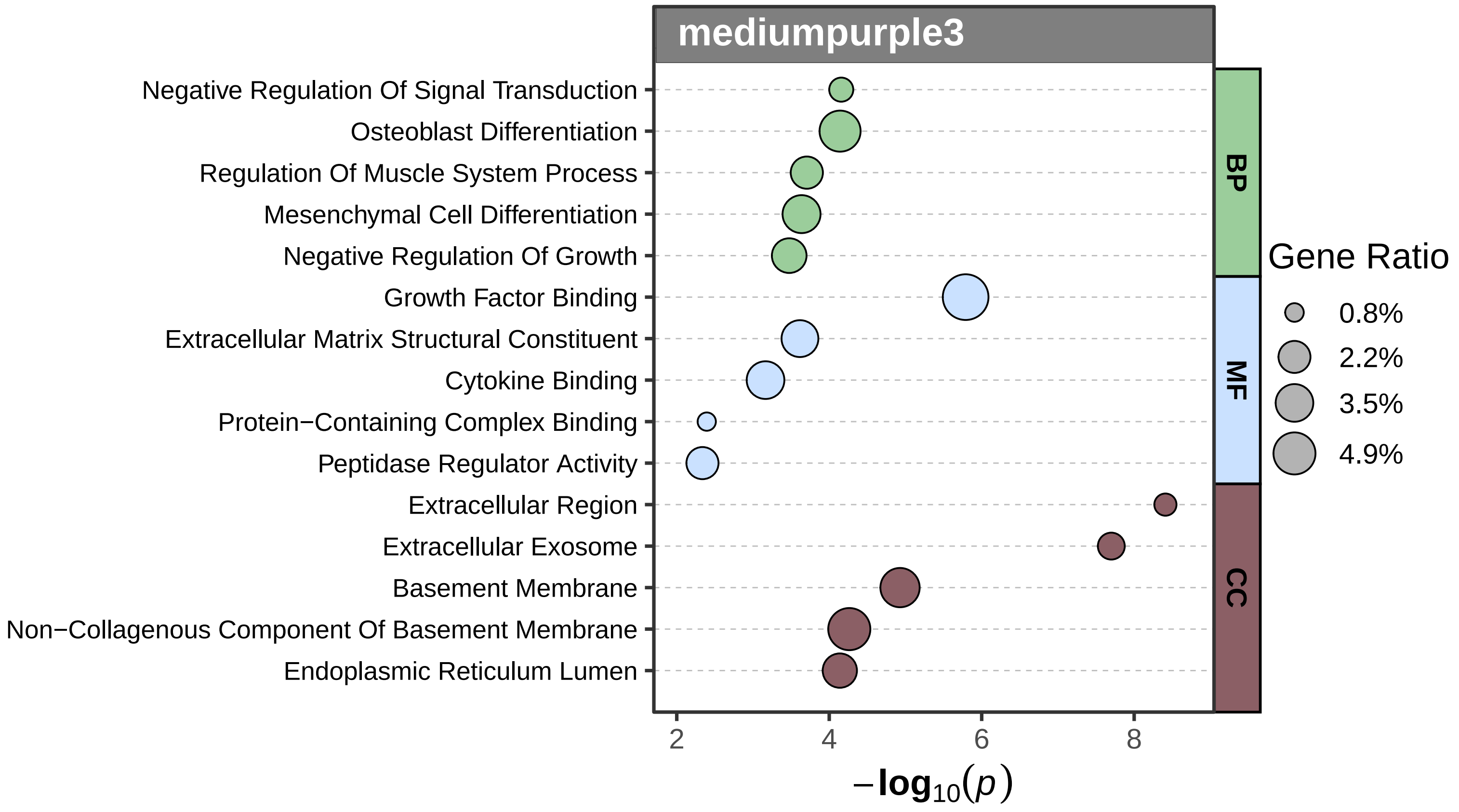

mediumpurple3
Negative Regulation Of Signal Transduction
Osteoblast Differentiation
Regulation Of Muscle System Process
Mesenchymal Cell Differentiation
Negative Regulation Of Growth
Growth Factor Binding
Extracellular Matrix Structural Constituent
Cytokine Binding
Protein−Containing Complex Binding
Peptidase Regulator Activity
Extracellular Region
Extracellular Exosome
Basement Membrane
Non−Collagenous Component Of Basement Membrane
Endoplasmic Reticulum Lumen
BP
MF
CC
Gene Ratio
0.8%
2.2%
3.5%
4.9%
2
4
6
8
$-\log_{10}(p)$

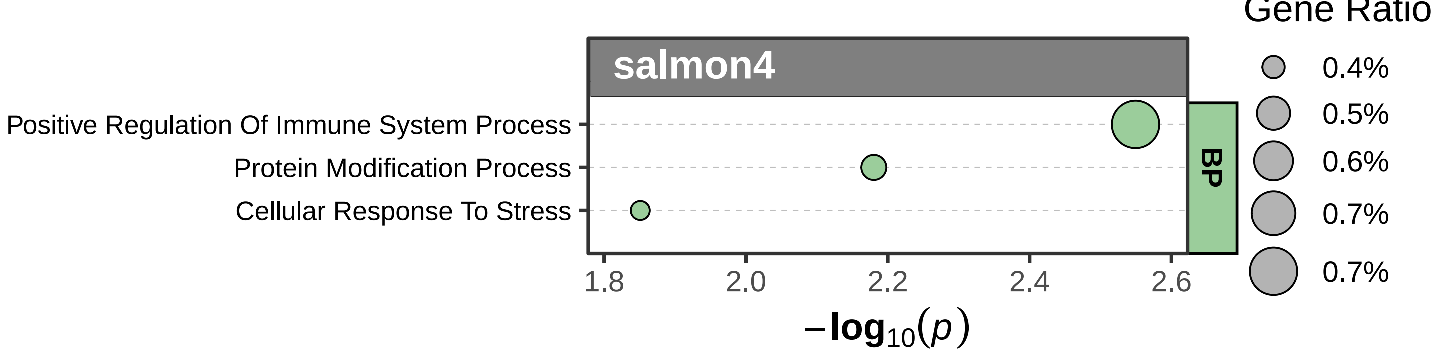
Gene Ratio
salmon4
Positive Regulation Of Immune System Process
Protein Modification Process
Cellular Response To Stress
BP
1.8
2.0
2.2
2.4
2.6
$-\log_{10}(p)$
0.4%
0.5%
0.6%
0.7%
0.7%

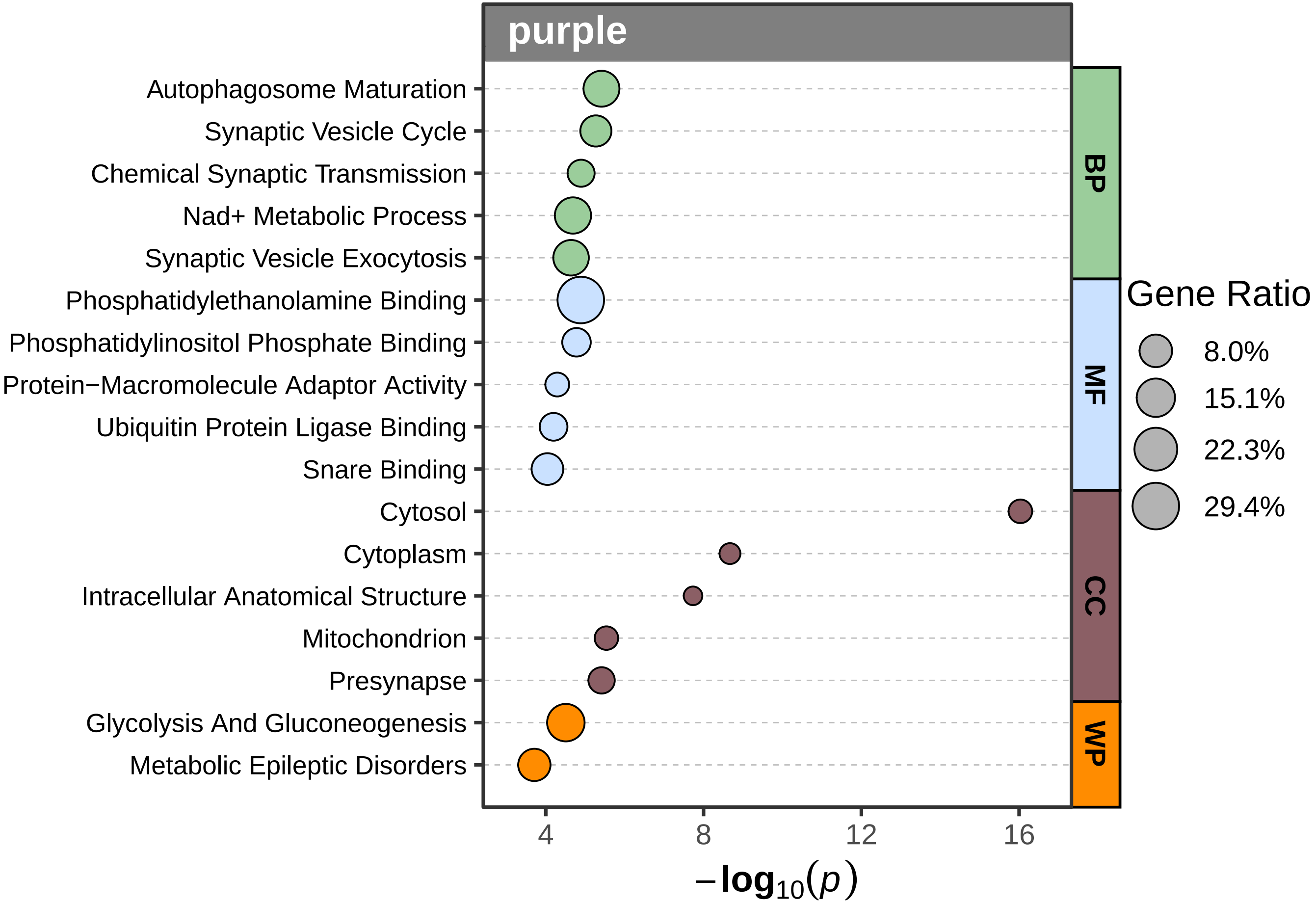
purple
Autophagosome Maturation
Synaptic Vesicle Cycle
Chemical Synaptic Transmission
Nad+ Metabolic Process
Synaptic Vesicle Exocytosis
Phosphatidylethanolamine Binding
Phosphatidylinositol Phosphate Binding
Protein−Macromolecule Adaptor Activity
Ubiquitin Protein Ligase Binding
Snare Binding
Cytosol
Cytoplasm
Intracellular Anatomical Structure
Mitochondrion
Presynapse
Glycolysis And Gluconeogenesis
Metabolic Epileptic Disorders
BP
MF
CC
WP
Gene Ratio
8.0%
15.1%
22.3%
29.4%
4
8
12
16
$-\log_{10}(p)$

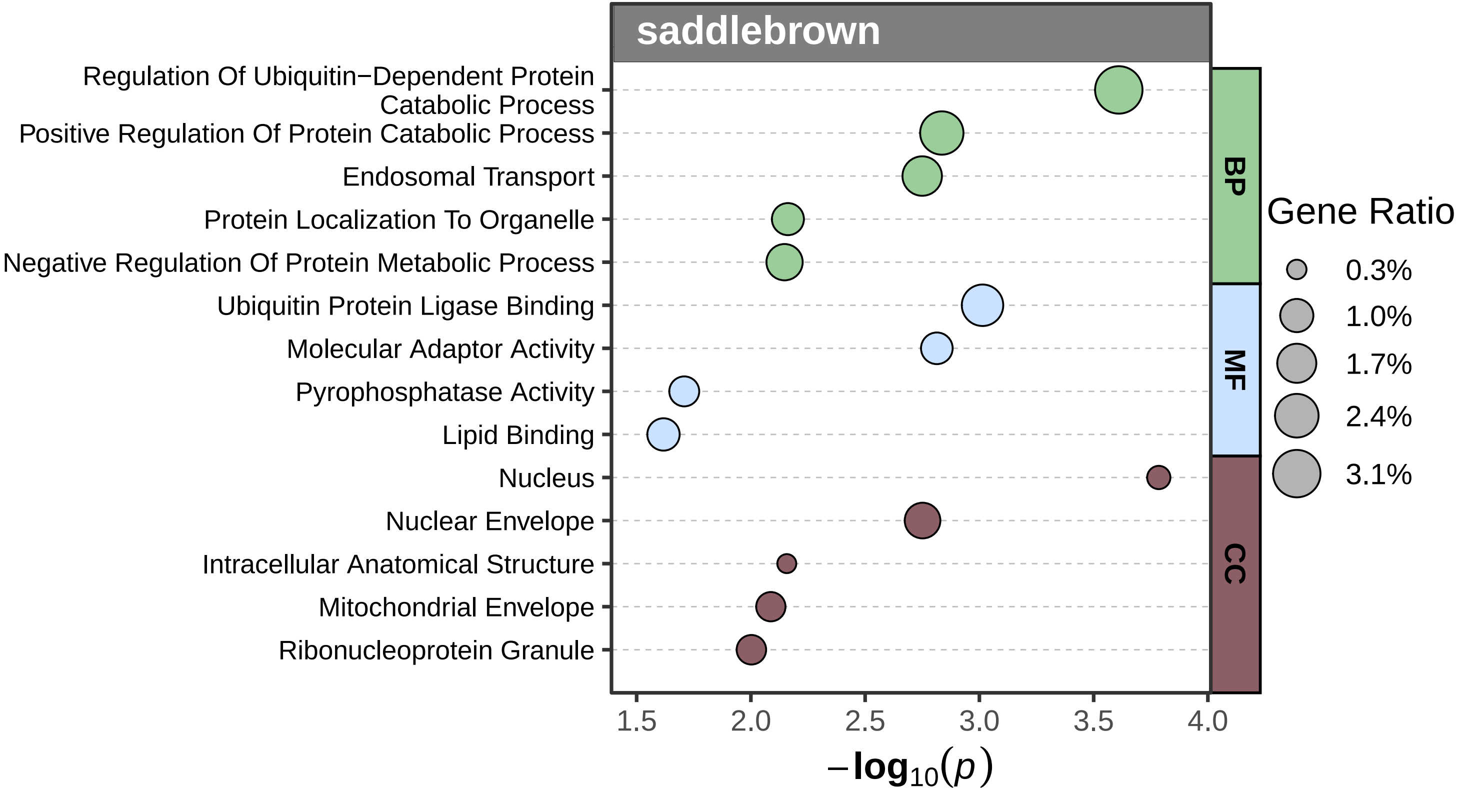
saddlebrown
Regulation Of Ubiquitin−Dependent Protein Catabolic Process
Positive Regulation Of Protein Catabolic Process
Endosomal Transport
Protein Localization To Organelle
Negative Regulation Of Protein Metabolic Process
Ubiquitin Protein Ligase Binding
Molecular Adaptor Activity
Pyrophosphatase Activity
Lipid Binding
Nucleus
Nuclear Envelope
Intracellular Anatomical Structure
Mitochondrial Envelope
Ribonucleoprotein Granule
BP
MF
CC
Gene Ratio
0.3%
1.0%
1.7%
2.4%
3.1%
1.5
2.0
2.5
3.0
3.5
4.0
$-\log_{10}(p)$

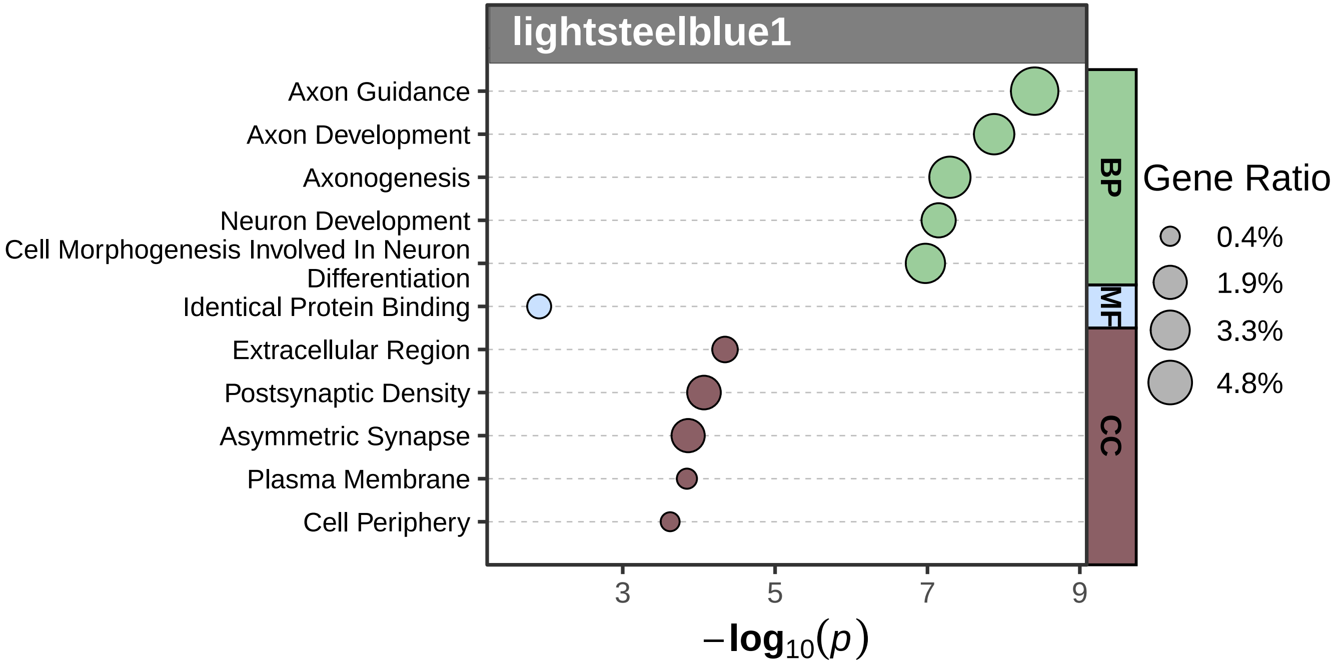
lightsteelblue1
Axon Guidance
Axon Development
Axonogenesis
Neuron Development
Cell Morphogenesis Involved In Neuron Differentiation
Identical Protein Binding
Extracellular Region
Postsynaptic Density
Asymmetric Synapse
Plasma Membrane
Cell Periphery
BP
MF
CC
Gene Ratio
0.4%
1.9%
3.3%
4.8%
3
5
7
9
$-\log_{10}(p)$

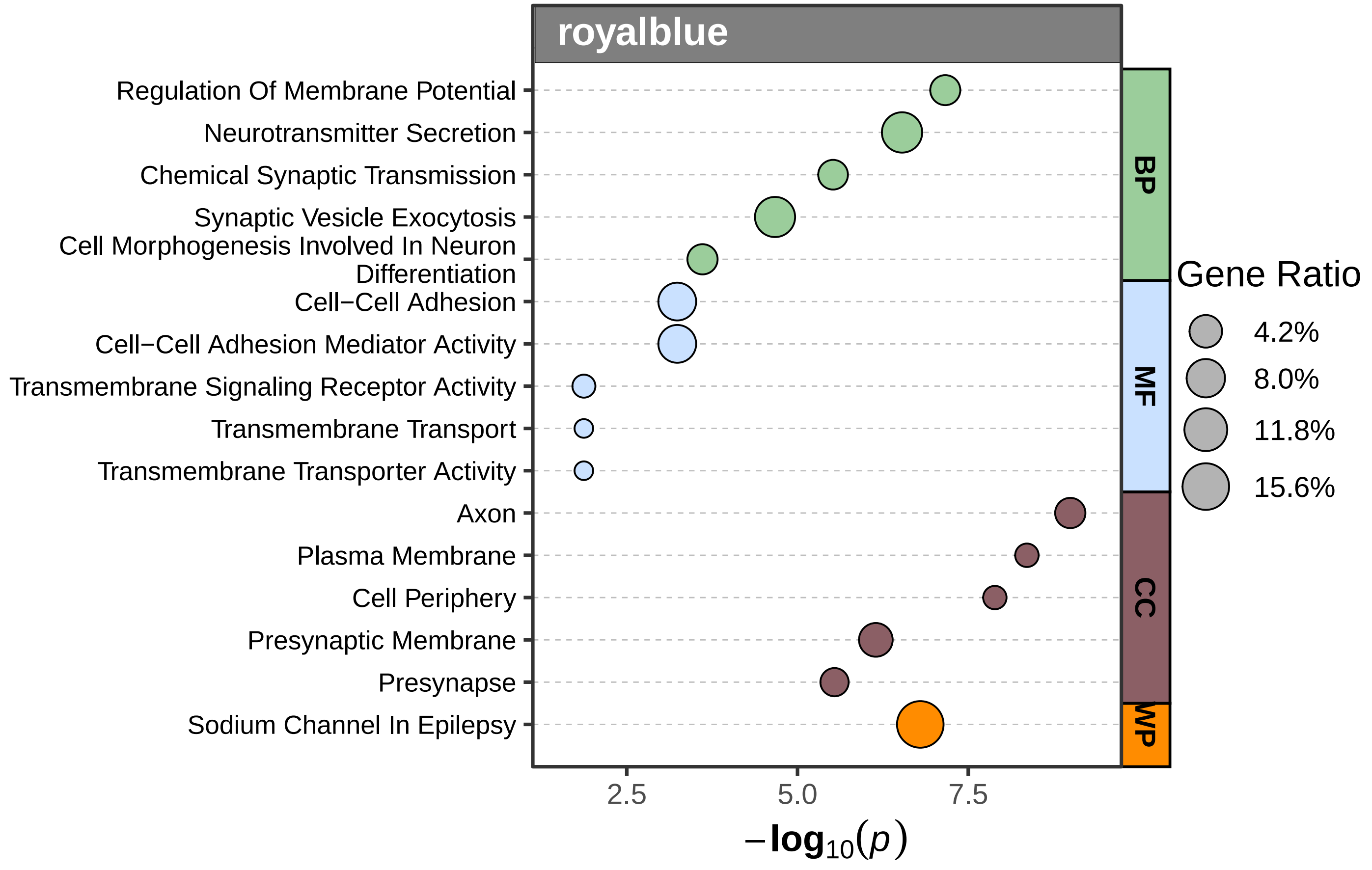

royalblue
Regulation Of Membrane Potential
Neurotransmitter Secretion
Chemical Synaptic Transmission
Synaptic Vesicle Exocytosis
Cell Morphogenesis Involved In Neuron Differentiation
Cell−Cell Adhesion
Cell−Cell Adhesion Mediator Activity
Transmembrane Signaling Receptor Activity
Transmembrane Transport
Transmembrane Transporter Activity
Axon
Plasma Membrane
Cell Periphery
Presynaptic Membrane
Presynapse
Sodium Channel In Epilepsy
BP
MF
CC
WP
Gene Ratio
4.2%
8.0%
11.8%
15.6%
2.5
5.0
7.5
$-\log_{10}(p)$

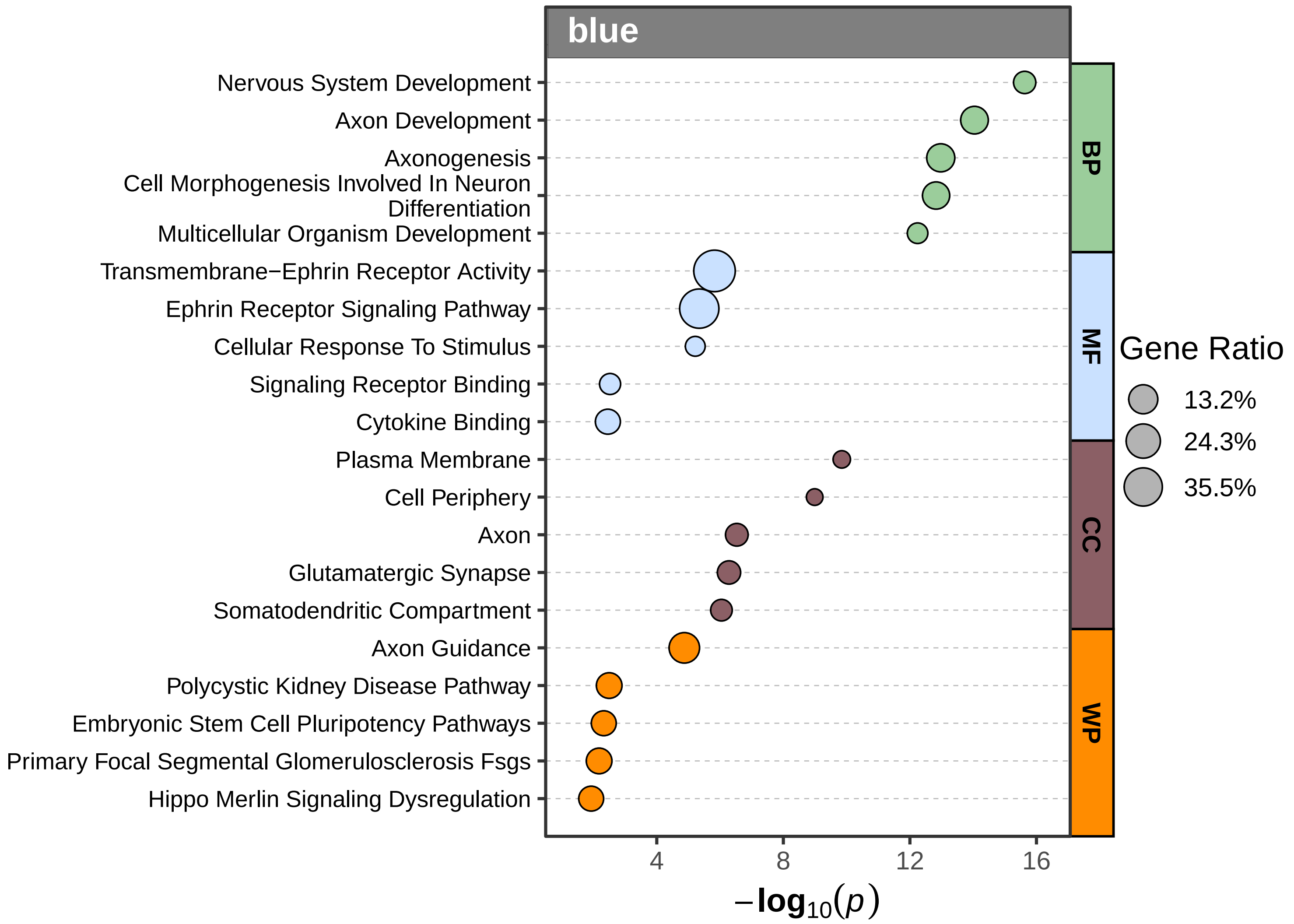
blue
Nervous System Development
Axon Development
Axonogenesis
Cell Morphogenesis Involved In Neuron Differentiation
Multicellular Organism Development
Transmembrane−Ephrin Receptor Activity
Ephrin Receptor Signaling Pathway
Cellular Response To Stimulus
Signaling Receptor Binding
Cytokine Binding
Plasma Membrane
Cell Periphery
Axon
Glutamatergic Synapse
Somatodendritic Compartment
Axon Guidance
Polycystic Kidney Disease Pathway
Embryonic Stem Cell Pluripotency Pathways
Primary Focal Segmental Glomerulosclerosis Fsgs
Hippo Merlin Signaling Dysregulation
BP
MF
CC
WP
Gene Ratio
13.2%
24.3%
35.5%
4
8
12
16
$-\log_{10}(p)$

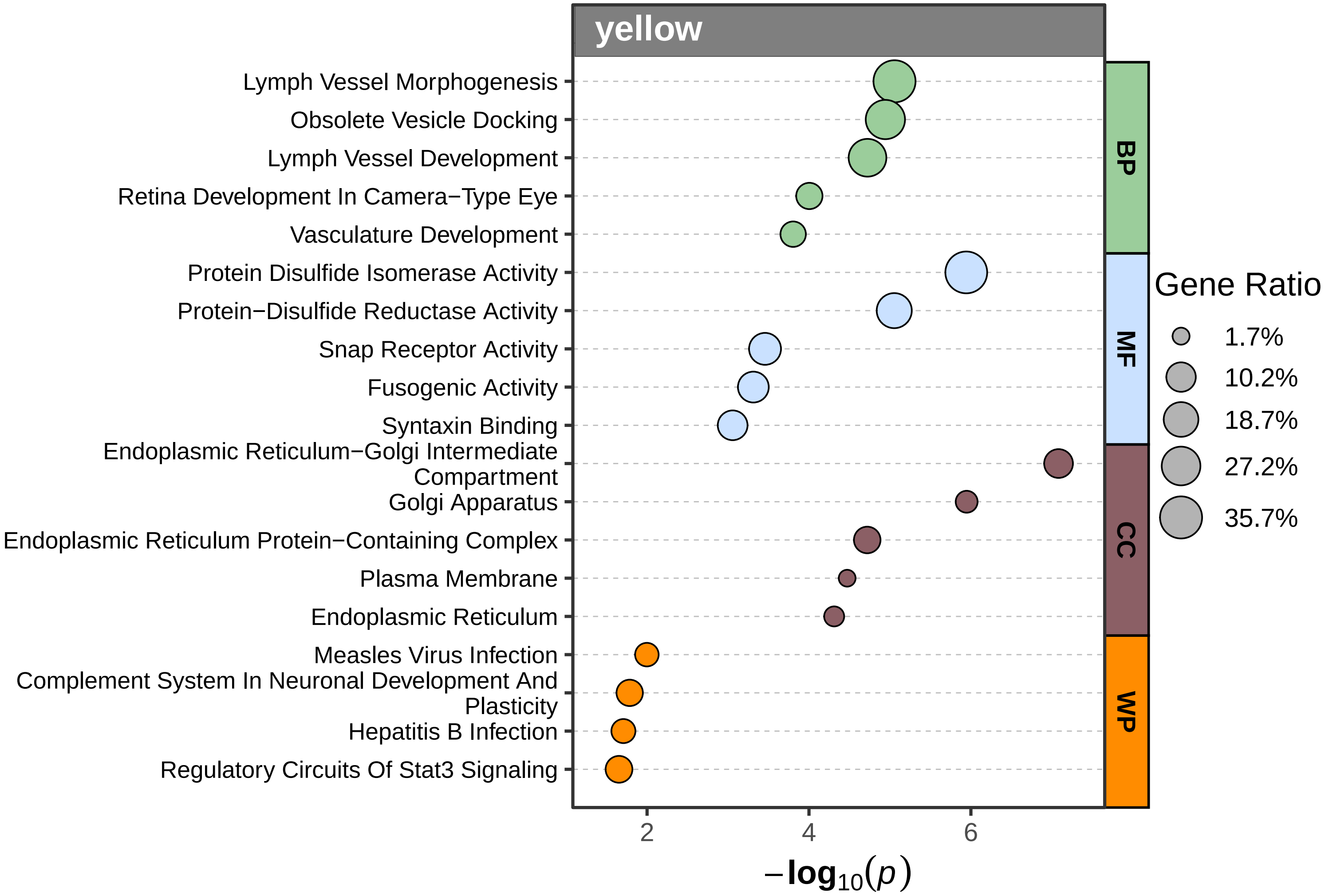
yellow
Lymph Vessel Morphogenesis
Obsolete Vesicle Docking
Lymph Vessel Development
Retina Development In Camera−Type Eye
Vasculature Development
Protein Disulfide Isomerase Activity
Protein−Disulfide Reductase Activity
Snap Receptor Activity
Fusogenic Activity
Syntaxin Binding
Endoplasmic Reticulum−Golgi Intermediate Compartment
Golgi Apparatus
Endoplasmic Reticulum Protein−Containing Complex
Plasma Membrane
Endoplasmic Reticulum
Measles Virus Infection
Complement System In Neuronal Development And Plasticity
Hepatitis B Infection
Regulatory Circuits Of Stat3 Signaling
BP
MF
CC
WP
2
4
6
$-\log_{10}(p)$
Gene Ratio
1.7%
10.2%
18.7%
27.2%
35.7%

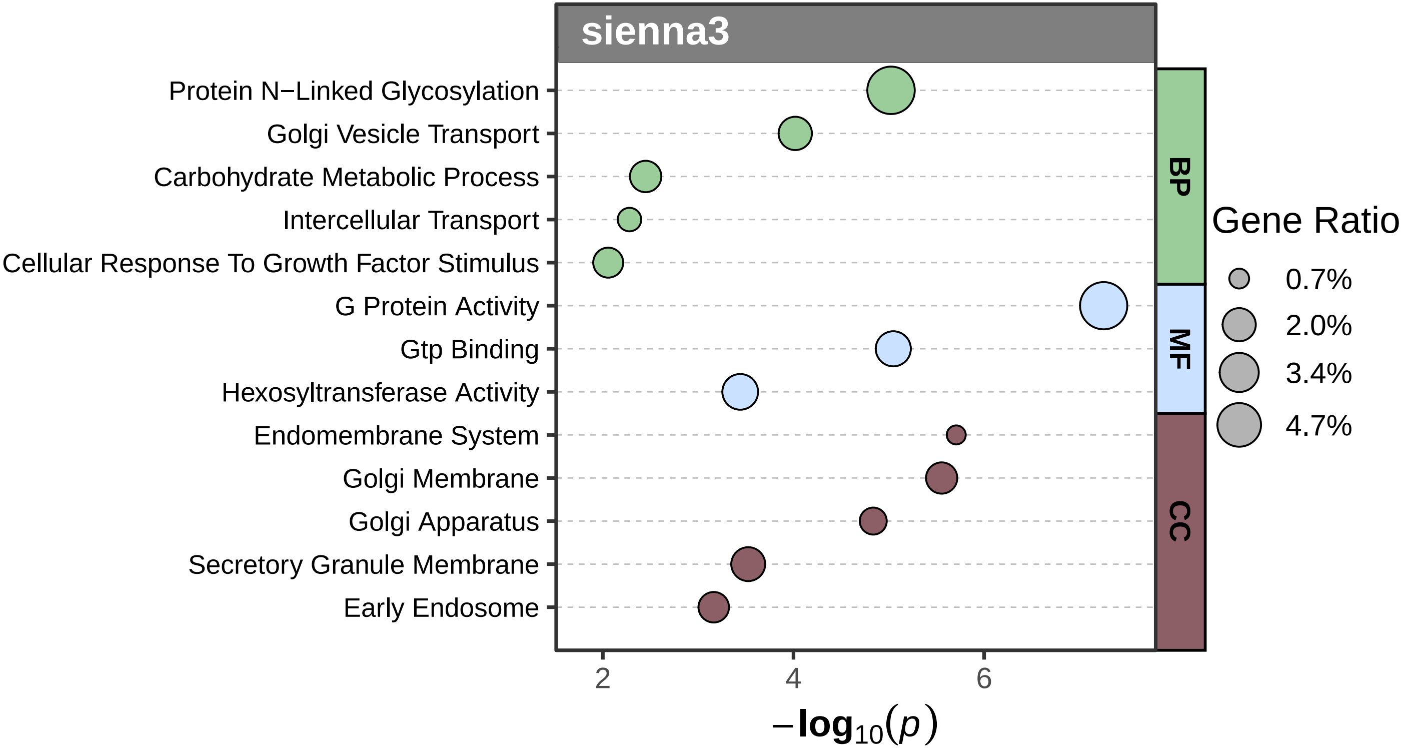
sienna3
Protein N−Linked Glycosylation
Golgi Vesicle Transport
Carbohydrate Metabolic Process
Intercellular Transport
Cellular Response To Growth Factor Stimulus
G Protein Activity
Gtp Binding
Hexosyltransferase Activity
Endomembrane System
Golgi Membrane
Golgi Apparatus
Secretory Granule Membrane
Early Endosome
BP
MF
CC
Gene Ratio
0.7%
2.0%
3.4%
4.7%
2
4
6
−log10(p)

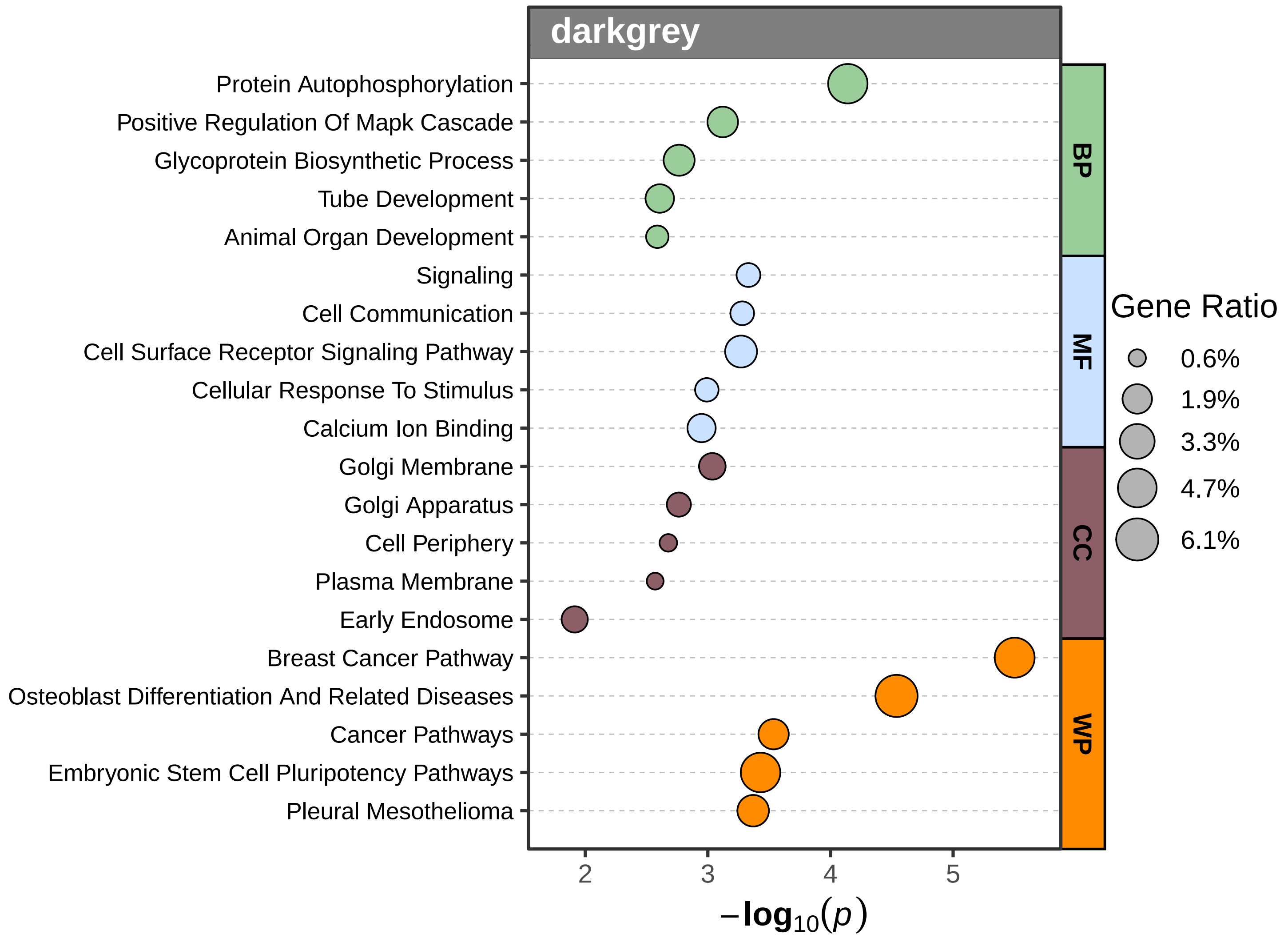
darkgrey
Protein Autophosphorylation
Positive Regulation Of Mapk Cascade
Glycoprotein Biosynthetic Process
Tube Development
Animal Organ Development
Signaling
Cell Communication
Cell Surface Receptor Signaling Pathway
Cellular Response To Stimulus
Calcium Ion Binding
Golgi Membrane
Golgi Apparatus
Cell Periphery
Plasma Membrane
Early Endosome
Breast Cancer Pathway
Osteoblast Differentiation And Related Diseases
Cancer Pathways
Embryonic Stem Cell Pluripotency Pathways
Pleural Mesothelioma
BP
MF
CC
WP
Gene Ratio
0.6%
1.9%
3.3%
4.7%
6.1%
2
3
4
5
$-\log_{10}(p)$

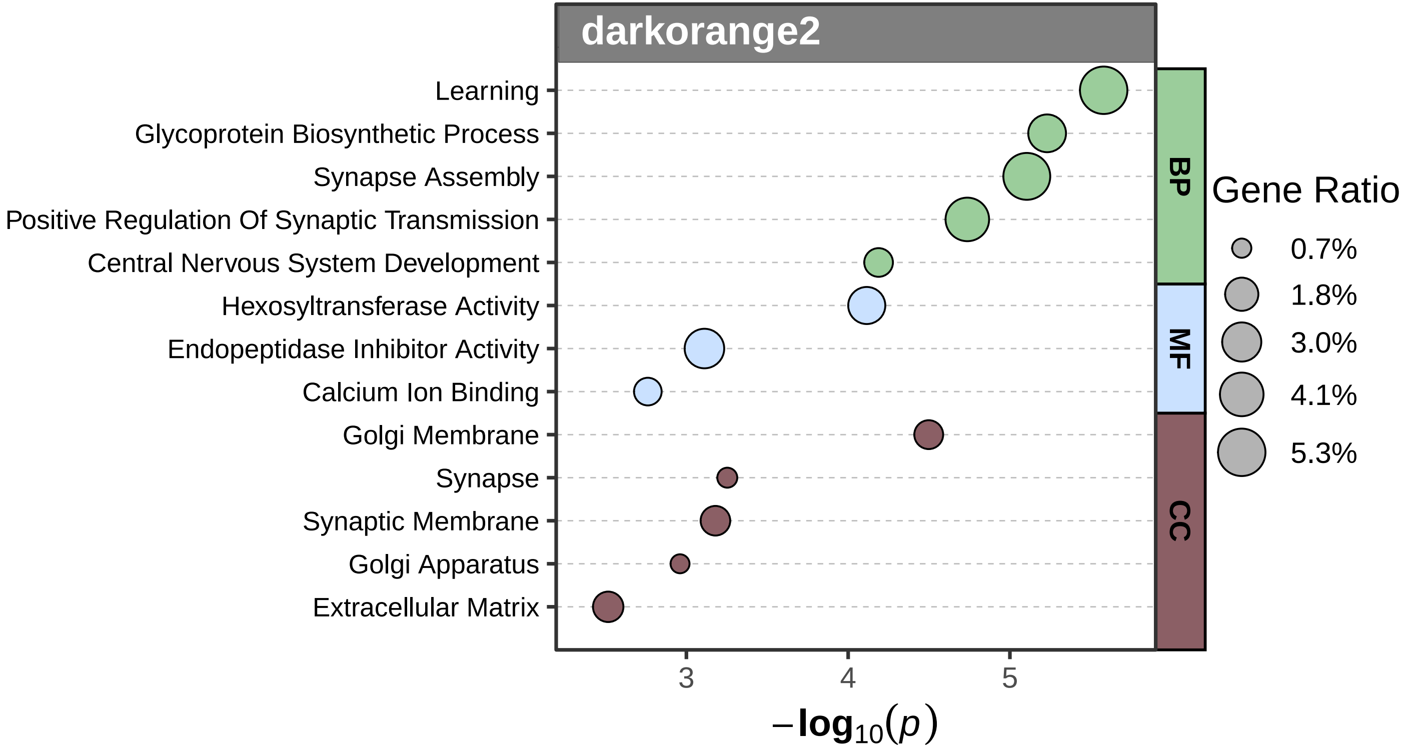
darkorange2
Learning
Glycoprotein Biosynthetic Process
Synapse Assembly
Positive Regulation Of Synaptic Transmission
Central Nervous System Development
Hexosyltransferase Activity
Endopeptidase Inhibitor Activity
Calcium Ion Binding
Golgi Membrane
Synapse
Synaptic Membrane
Golgi Apparatus
Extracellular Matrix
BP
MF
CC
Gene Ratio
0.7%
1.8%
3.0%
4.1%
5.3%
3
4
5
$-\log_{10}(p)$

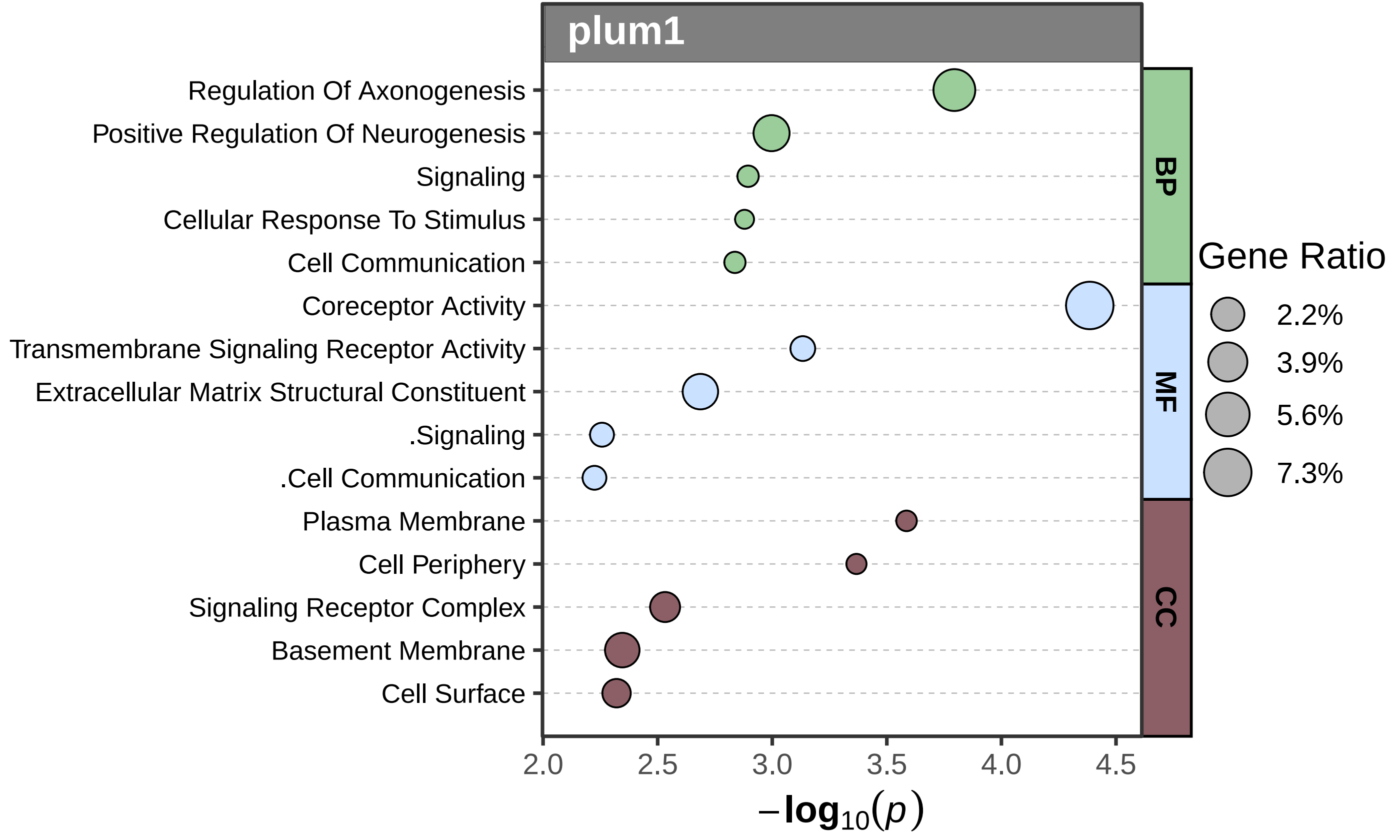
plum1
Regulation Of Axonogenesis
Positive Regulation Of Neurogenesis
Signaling
Cellular Response To Stimulus
Cell Communication
Coreceptor Activity
Transmembrane Signaling Receptor Activity
Extracellular Matrix Structural Constituent
.Signaling
.Cell Communication
Plasma Membrane
Cell Periphery
Signaling Receptor Complex
Basement Membrane
Cell Surface
BP
MF
CC
Gene Ratio
2.2%
3.9%
5.6%
7.3%
2.0
2.5
3.0
3.5
4.0
4.5
$-\log_{10}(p)$

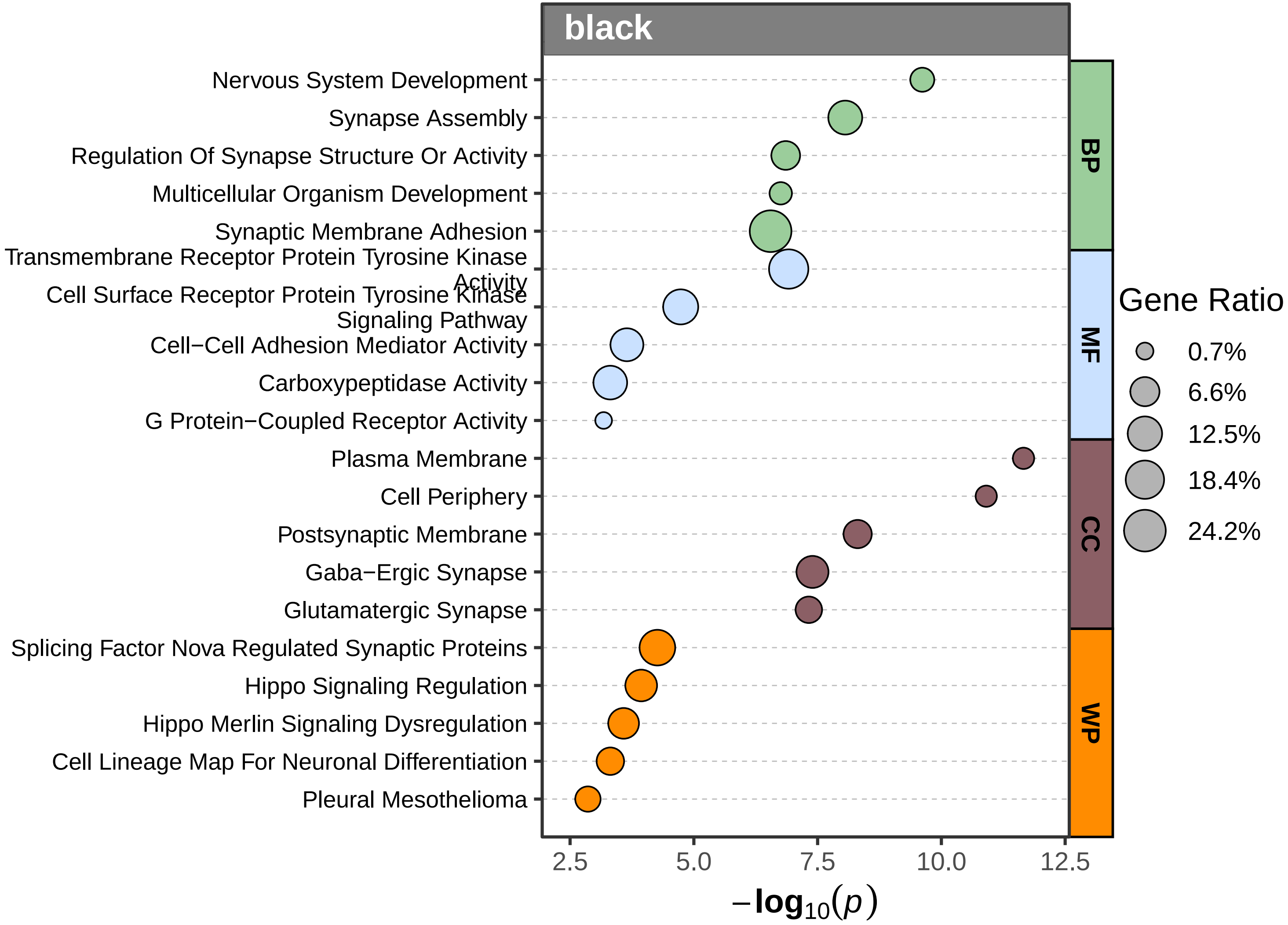

black
Nervous System Development
Synapse Assembly
Regulation Of Synapse Structure Or Activity
Multicellular Organism Development
Synaptic Membrane Adhesion
Transmembrane Receptor Protein Tyrosine Kinase Activity
Cell Surface Receptor Protein Tyrosine Kinase Signaling Pathway
Cell−Cell Adhesion Mediator Activity
Carboxypeptidase Activity
G Protein−Coupled Receptor Activity
Plasma Membrane
Cell Periphery
Postsynaptic Membrane
Gaba−Ergic Synapse
Glutamatergic Synapse
Splicing Factor Nova Regulated Synaptic Proteins
Hippo Signaling Regulation
Hippo Merlin Signaling Dysregulation
Cell Lineage Map For Neuronal Differentiation
Pleural Mesothelioma
BP
MF
CC
WP
Gene Ratio
0.7%
6.6%
12.5%
18.4%
24.2%
2.5
5.0
7.5
10.0
12.5
$-\log_{10}(p)$

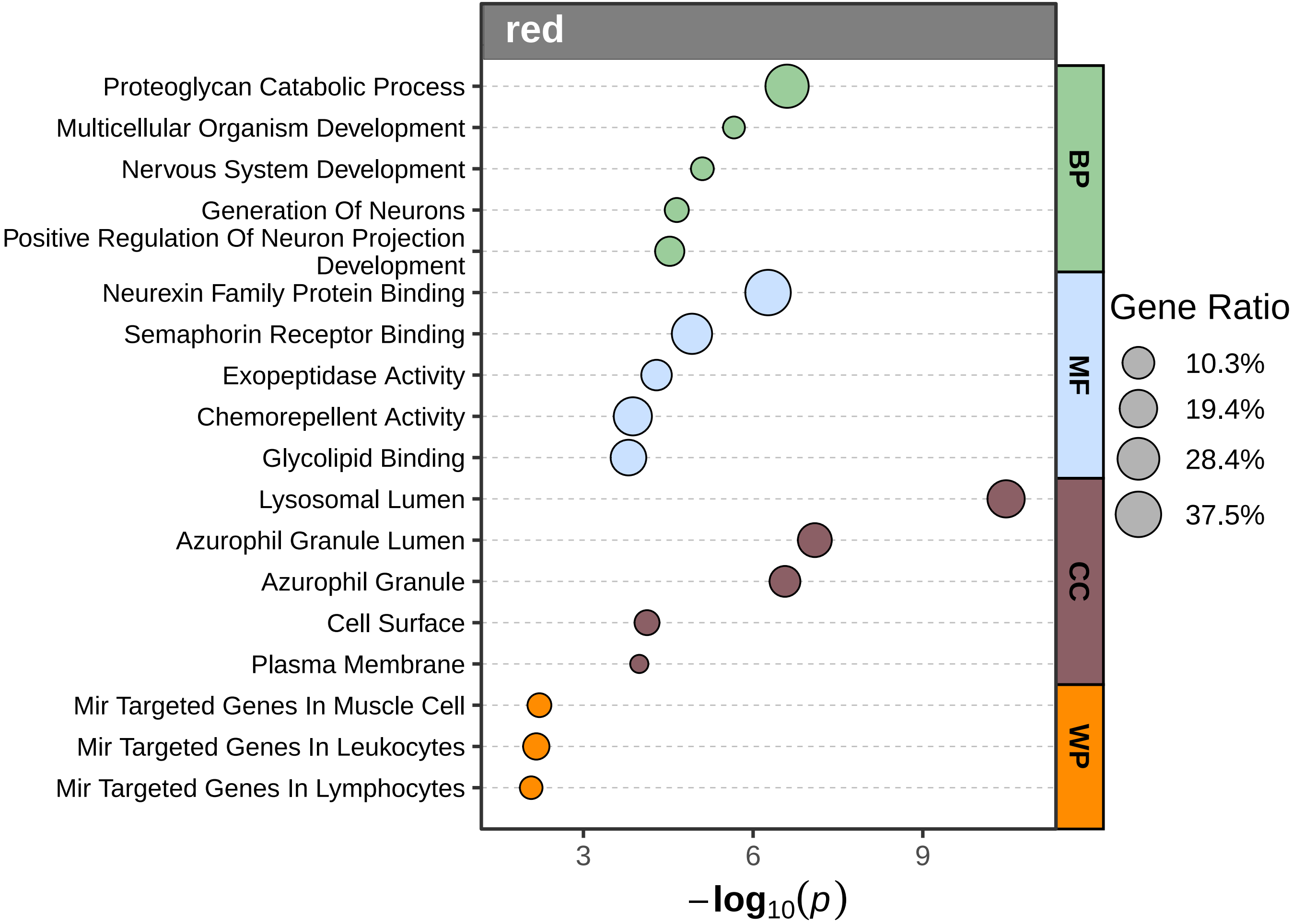

red
Proteoglycan Catabolic Process
Multicellular Organism Development
Nervous System Development
Generation Of Neurons
Positive Regulation Of Neuron Projection Development
Neurexin Family Protein Binding
Semaphorin Receptor Binding
Exopeptidase Activity
Chemorepellent Activity
Glycolipid Binding
Lysosomal Lumen
Azurophil Granule Lumen
Azurophil Granule
Cell Surface
Plasma Membrane
Mir Targeted Genes In Muscle Cell
Mir Targeted Genes In Leukocytes
Mir Targeted Genes In Lymphocytes
BP
MF
CC
WP
3
6
9
$-\log_{10}(p)$
Gene Ratio
10.3%
19.4%
28.4%
37.5%

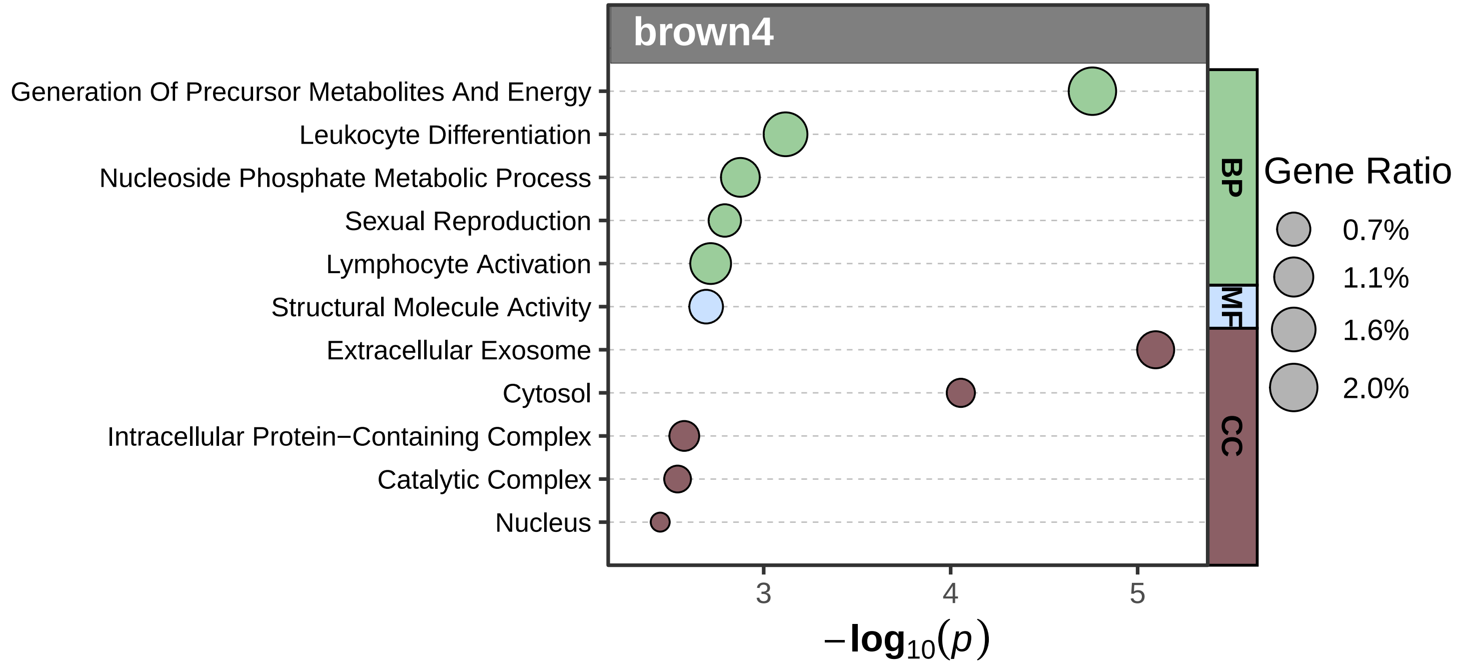
brown4
Generation Of Precursor Metabolites And Energy
Leukocyte Differentiation
Nucleoside Phosphate Metabolic Process
Sexual Reproduction
Lymphocyte Activation
Structural Molecule Activity
Extracellular Exosome
Cytosol
Intracellular Protein−Containing Complex
Catalytic Complex
Nucleus
BP
MF
CC
Gene Ratio
0.7%
1.1%
1.6%
2.0%
3
4
5
$-\log_{10}(p)$

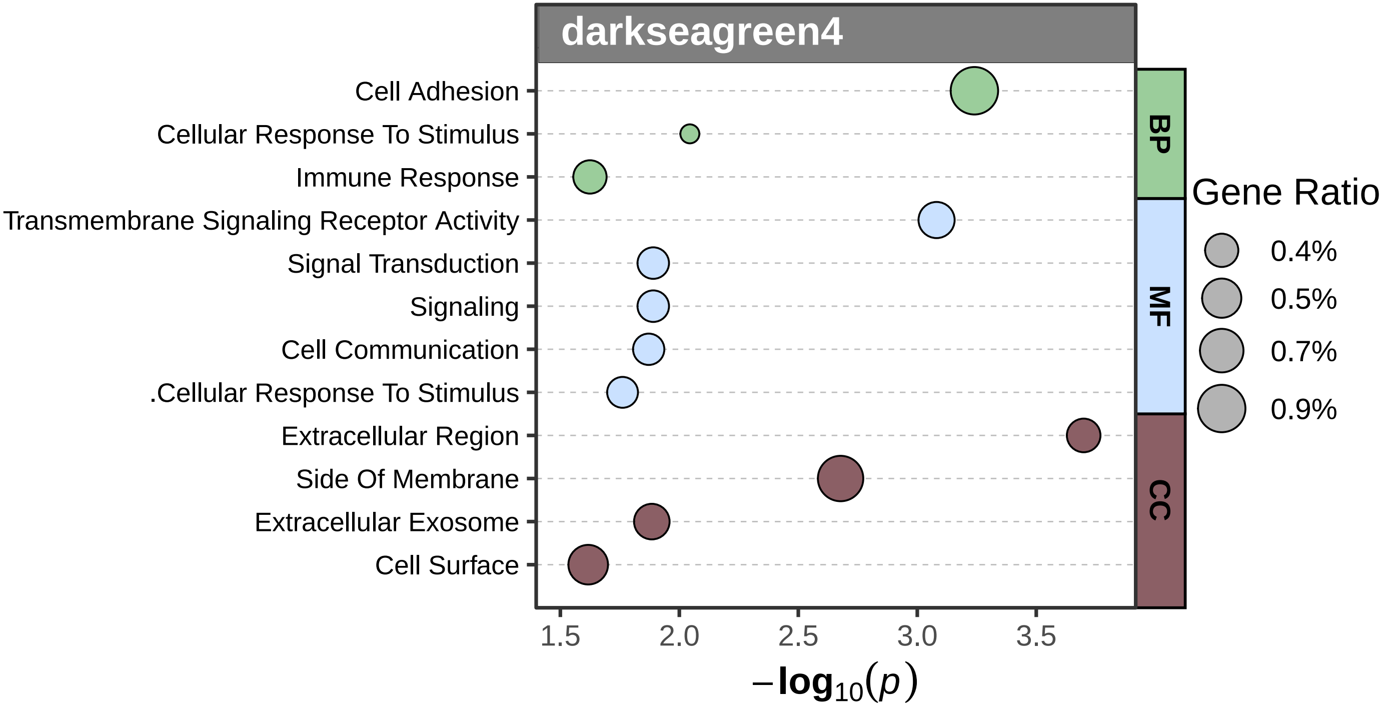
darkseagreen4
Cell Adhesion
Cellular Response To Stimulus
Immune Response
Transmembrane Signaling Receptor Activity
Signal Transduction
Signaling
Cell Communication
.Cellular Response To Stimulus
Extracellular Region
Side Of Membrane
Extracellular Exosome
Cell Surface
BP
MF
CC
Gene Ratio
0.4%
0.5%
0.7%
0.9%
1.5
2.0
2.5
3.0
3.5
$-\log_{10}(p)$

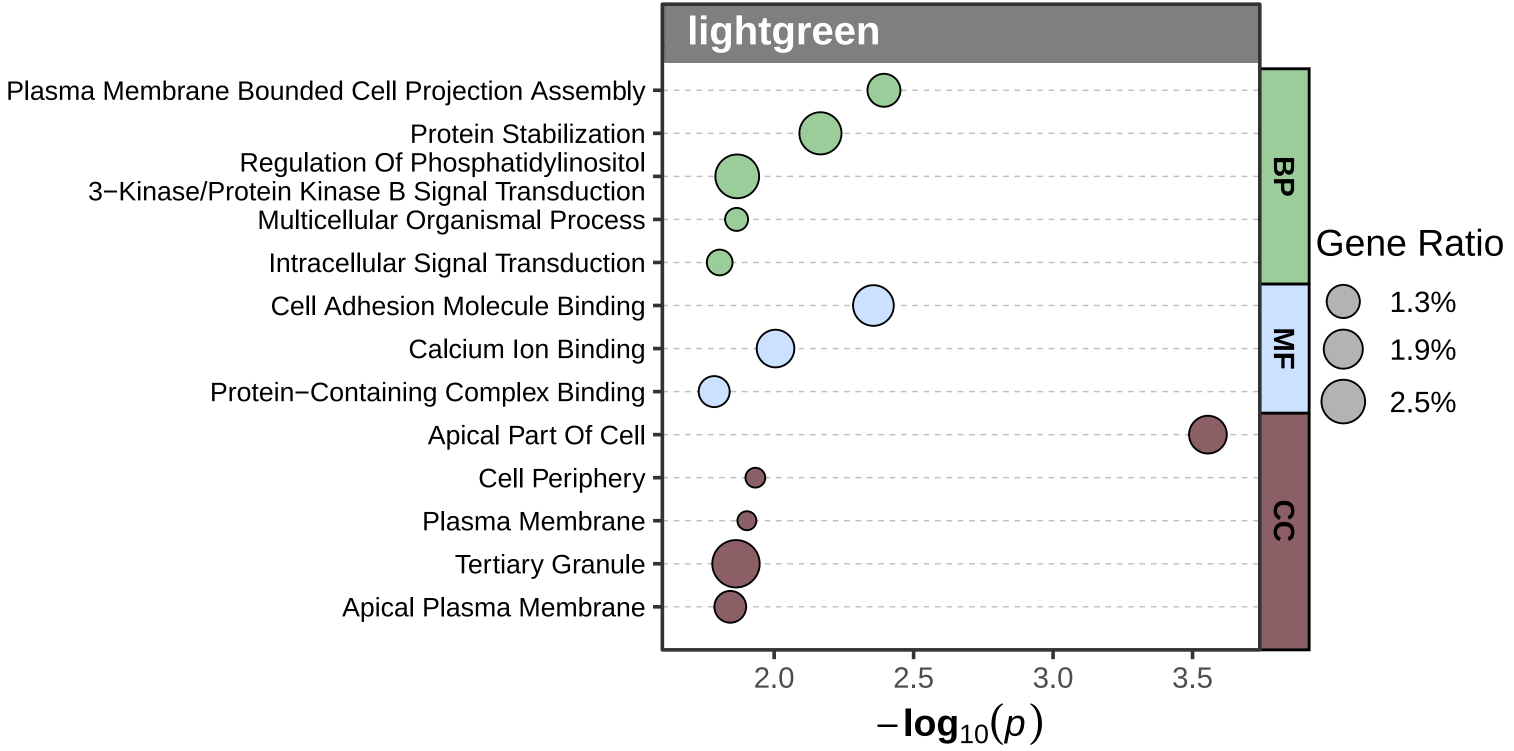

lightgreen
Plasma Membrane Bounded Cell Projection Assembly
Protein Stabilization
Regulation Of Phosphatidylinositol 3−Kinase/Protein Kinase B Signal Transduction
Multicellular Organismal Process
Intracellular Signal Transduction
Cell Adhesion Molecule Binding
Calcium Ion Binding
Protein−Containing Complex Binding
Apical Part Of Cell
Cell Periphery
Plasma Membrane
Tertiary Granule
Apical Plasma Membrane
BP
MF
CC
Gene Ratio
1.3%
1.9%
2.5%
2.0
2.5
3.0
3.5
−log10(p)

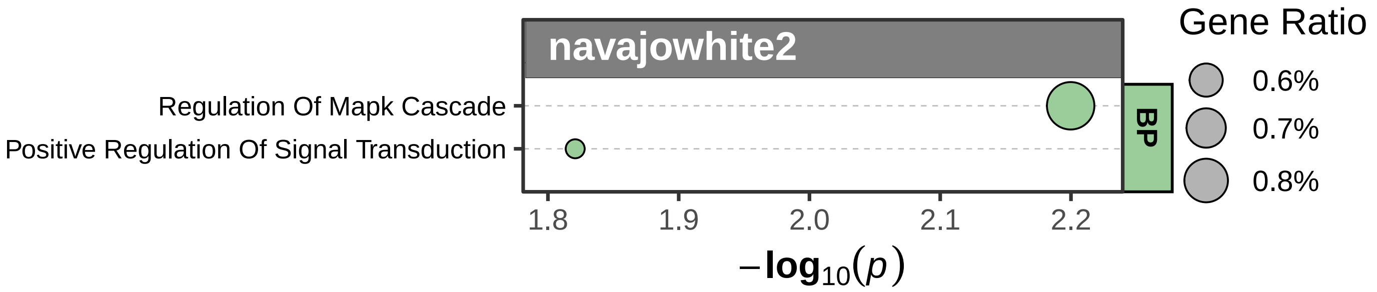
navajowhite2
Regulation Of Mapk Cascade
Positive Regulation Of Signal Transduction
BP
1.8
1.9
2.0
2.1
2.2
$-\log_{10}(p)$
Gene Ratio
0.6%
0.7%
0.8%

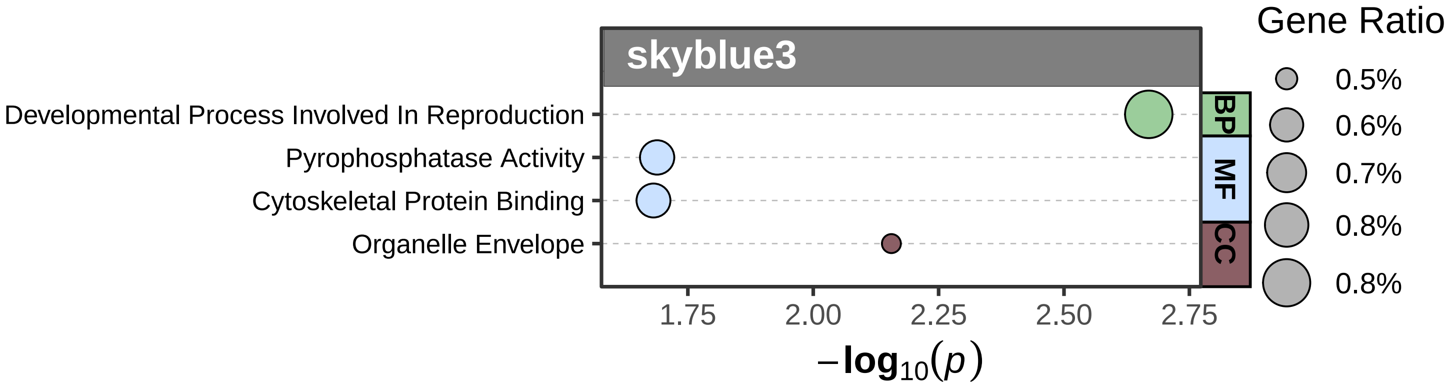

skyblue3
Developmental Process Involved In Reproduction
Pyrophosphatase Activity
Cytoskeletal Protein Binding
Organelle Envelope
BP
MF
CC
1.75
2.00
2.25
2.50
2.75
$-\log_{10}(p)$
Gene Ratio
0.5%
0.6%
0.7%
0.8%
0.8%

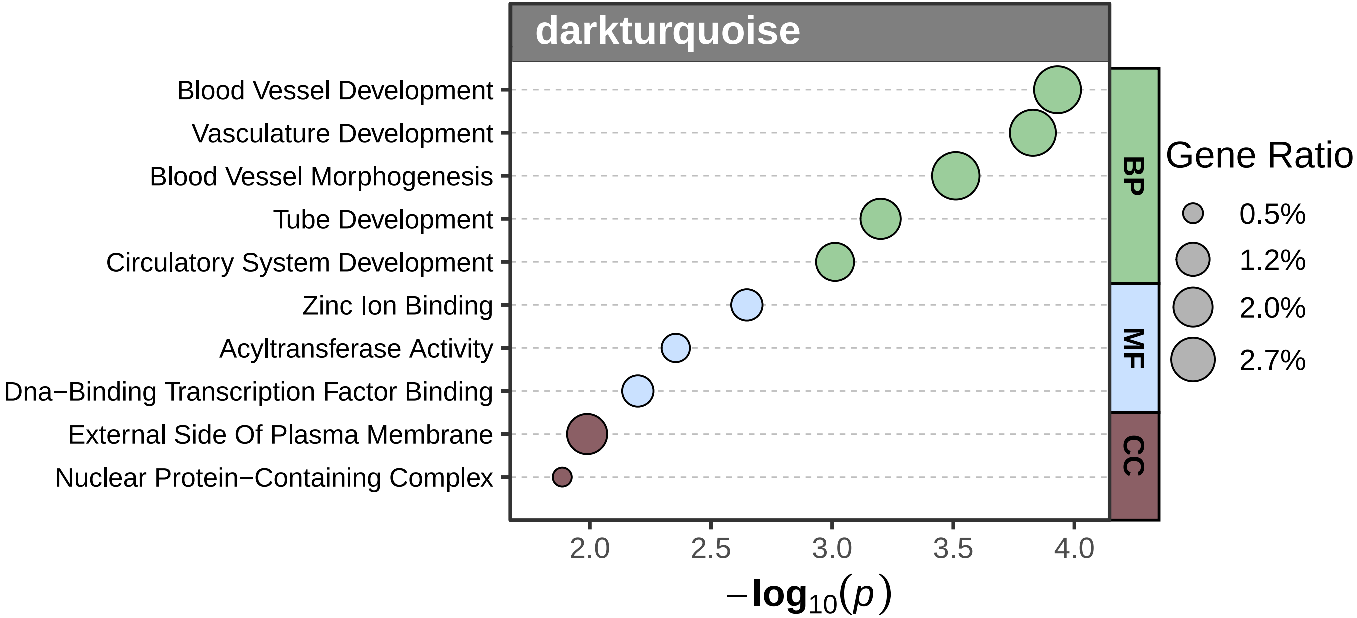
darkturquoise
Blood Vessel Development
Vasculature Development
Blood Vessel Morphogenesis
Tube Development
Circulatory System Development
Zinc Ion Binding
Acyltransferase Activity
Dna−Binding Transcription Factor Binding
External Side Of Plasma Membrane
Nuclear Protein−Containing Complex
BP
MF
CC
Gene Ratio
0.5%
1.2%
2.0%
2.7%
2.0
2.5
3.0
3.5
4.0
$-\log_{10}(p)$

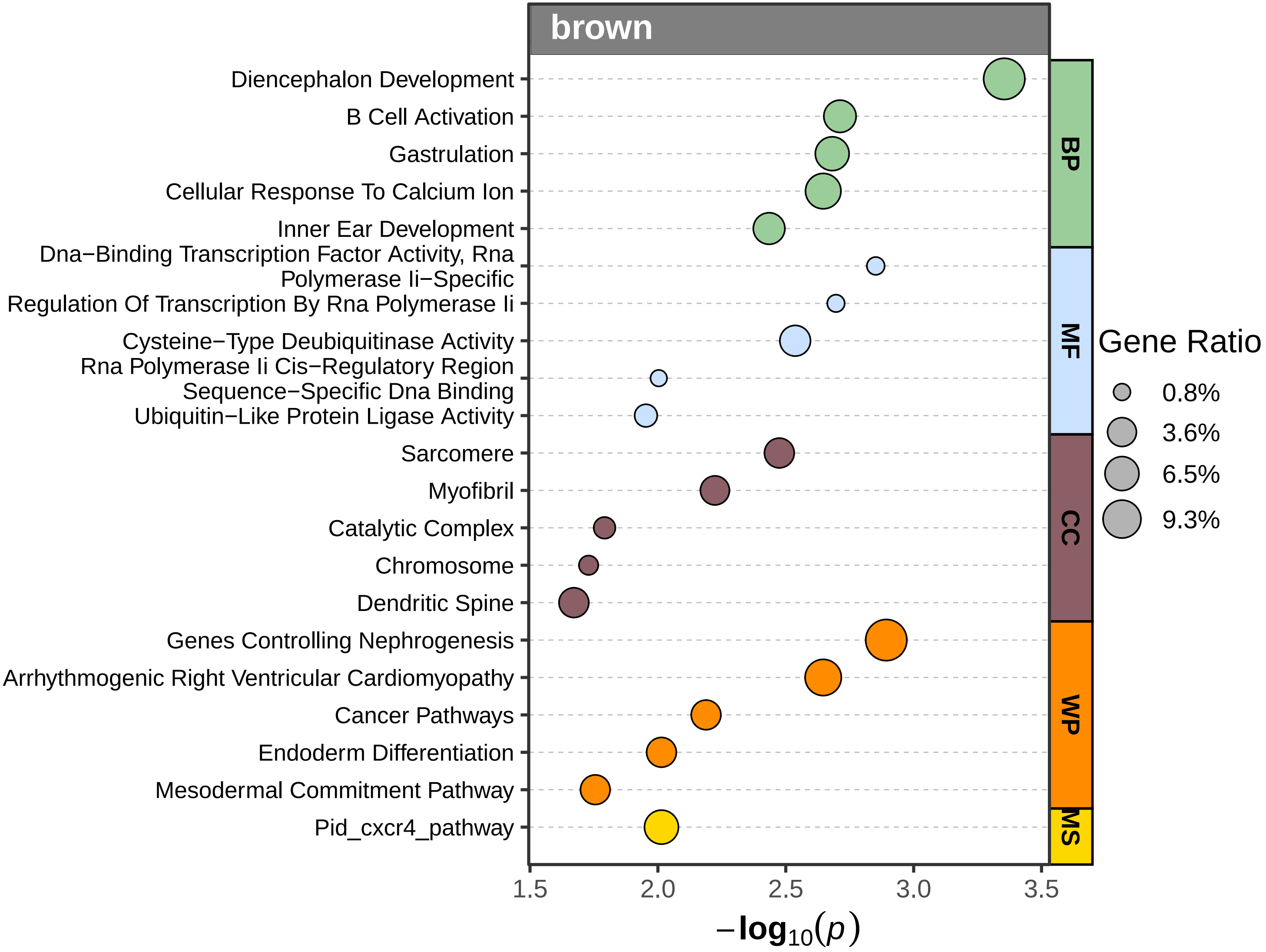

brown
Diencephalon Development
B Cell Activation
Gastrulation
Cellular Response To Calcium Ion
Inner Ear Development
Dna−Binding Transcription Factor Activity, Rna Polymerase Ii−Specific
Regulation Of Transcription By Rna Polymerase Ii
Cysteine−Type Deubiquitinase Activity
Rna Polymerase Ii Cis−Regulatory Region Sequence−Specific Dna Binding
Ubiquitin−Like Protein Ligase Activity
Sarcomere
Myofibril
Catalytic Complex
Chromosome
Dendritic Spine
Genes Controlling Nephrogenesis
Arrhythmogenic Right Ventricular Cardiomyopathy
Cancer Pathways
Endoderm Differentiation
Mesodermal Commitment Pathway
Pid_cxcr4_pathway
BP
MF
CC
WP
MS
1.5
2.0
2.5
3.0
3.5
$-\log_{10}(p)$
Gene Ratio
0.8%
3.6%
6.5%
9.3%

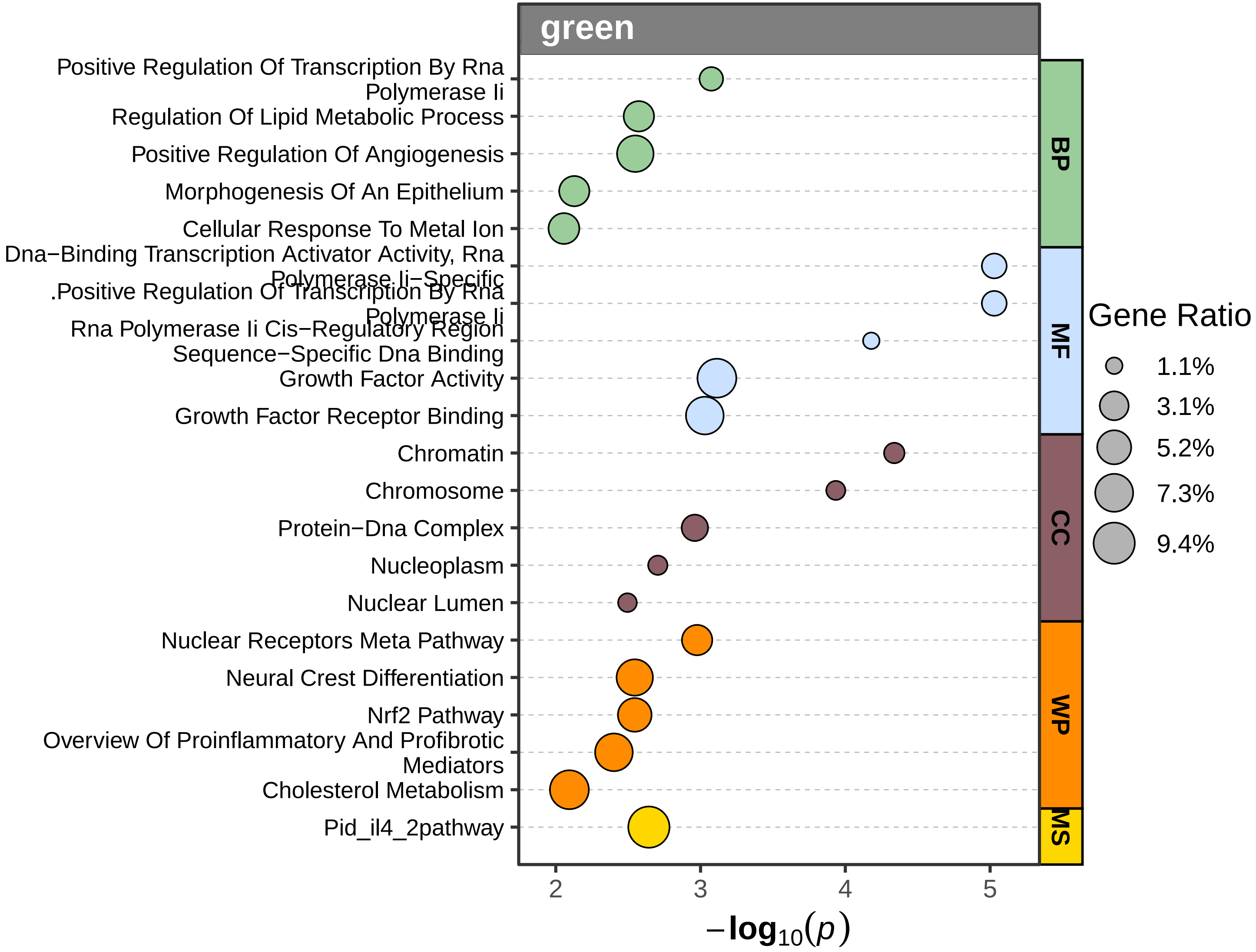
green
Positive Regulation Of Transcription By Rna Polymerase Ii
Regulation Of Lipid Metabolic Process
Positive Regulation Of Angiogenesis
Morphogenesis Of An Epithelium
Cellular Response To Metal Ion
Dna−Binding Transcription Activator Activity, Rna Polymerase Ii−Specific
.Positive Regulation Of Transcription By Rna Polymerase Ii
Rna Polymerase Ii Cis−Regulatory Region Sequence−Specific Dna Binding
Growth Factor Activity
Growth Factor Receptor Binding
Chromatin
Chromosome
Protein−Dna Complex
Nucleoplasm
Nuclear Lumen
Nuclear Receptors Meta Pathway
Neural Crest Differentiation
Nrf2 Pathway
Overview Of Proinflammatory And Profibrotic Mediators
Cholesterol Metabolism
Pid_il4_2pathway
BP
MF
CC
WP
MS
Gene Ratio
1.1%
3.1%
5.2%
7.3%
9.4%
2
3
4
5
$-\log_{10}(p)$

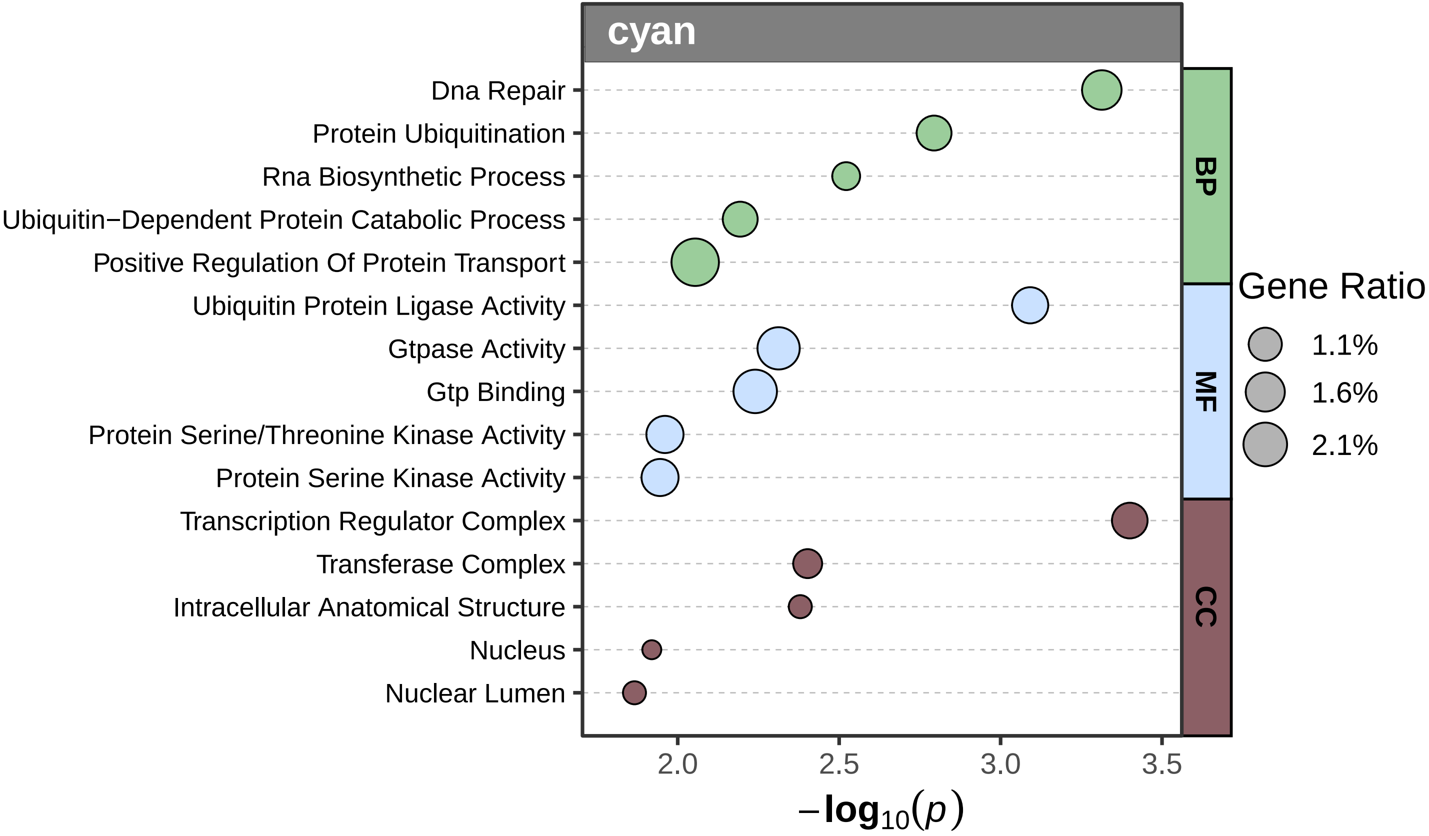

cyan
Dna Repair
Protein Ubiquitination
Rna Biosynthetic Process
Ubiquitin−Dependent Protein Catabolic Process
Positive Regulation Of Protein Transport
Ubiquitin Protein Ligase Activity
Gtpase Activity
Gtp Binding
Protein Serine/Threonine Kinase Activity
Protein Serine Kinase Activity
Transcription Regulator Complex
Transferase Complex
Intracellular Anatomical Structure
Nucleus
Nuclear Lumen
BP
MF
CC
Gene Ratio
1.1%
1.6%
2.1%
2.0
2.5
3.0
3.5
$-\log_{10}(p)$

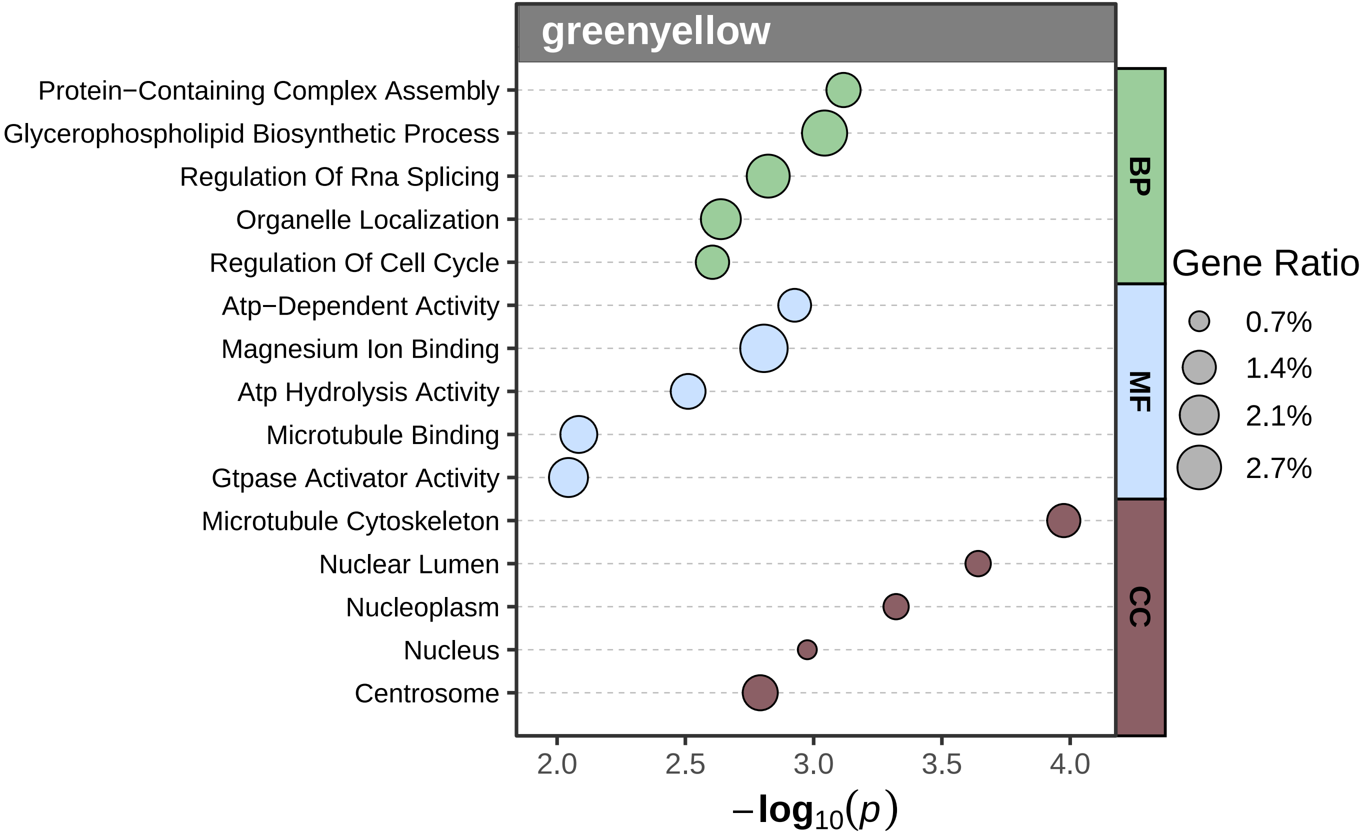
greenyellow
Protein−Containing Complex Assembly
Glycerophospholipid Biosynthetic Process
Regulation Of Rna Splicing
Organelle Localization
Regulation Of Cell Cycle
Atp−Dependent Activity
Magnesium Ion Binding
Atp Hydrolysis Activity
Microtubule Binding
Gtpase Activator Activity
Microtubule Cytoskeleton
Nuclear Lumen
Nucleoplasm
Nucleus
Centrosome
BP
MF
CC
2.0
2.5
3.0
3.5
4.0
$-\log_{10}(p)$
Gene Ratio
0.7%
1.4%
2.1%
2.7%

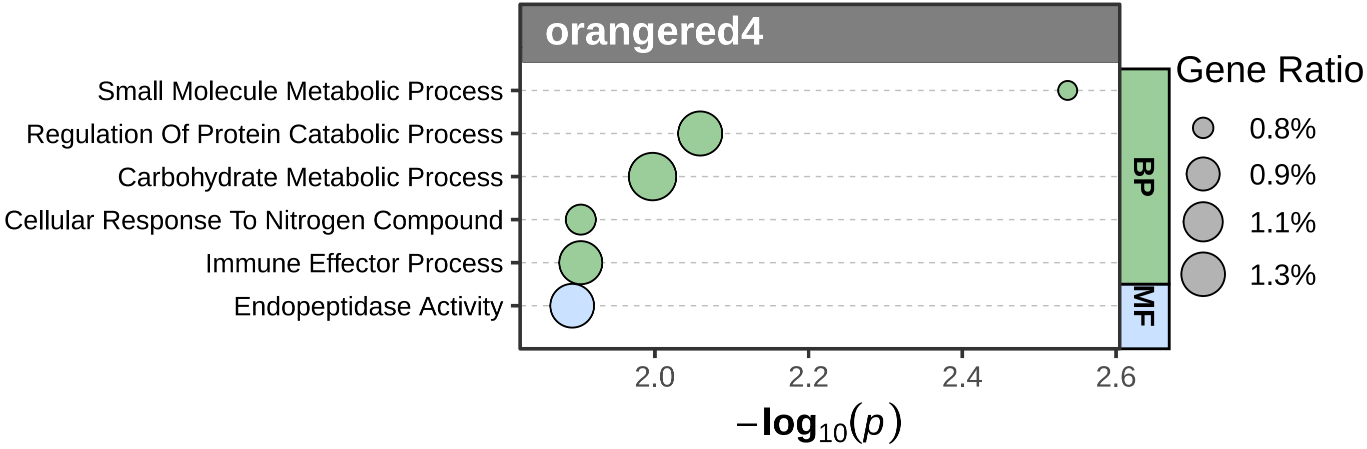

orangered4
Small Molecule Metabolic Process
Regulation Of Protein Catabolic Process
Carbohydrate Metabolic Process
Cellular Response To Nitrogen Compound
Immune Effector Process
Endopeptidase Activity
BP
MF
Gene Ratio
0.8%
0.9%
1.1%
1.3%
2.0
2.2
2.4
2.6
$-\log_{10}(p)$

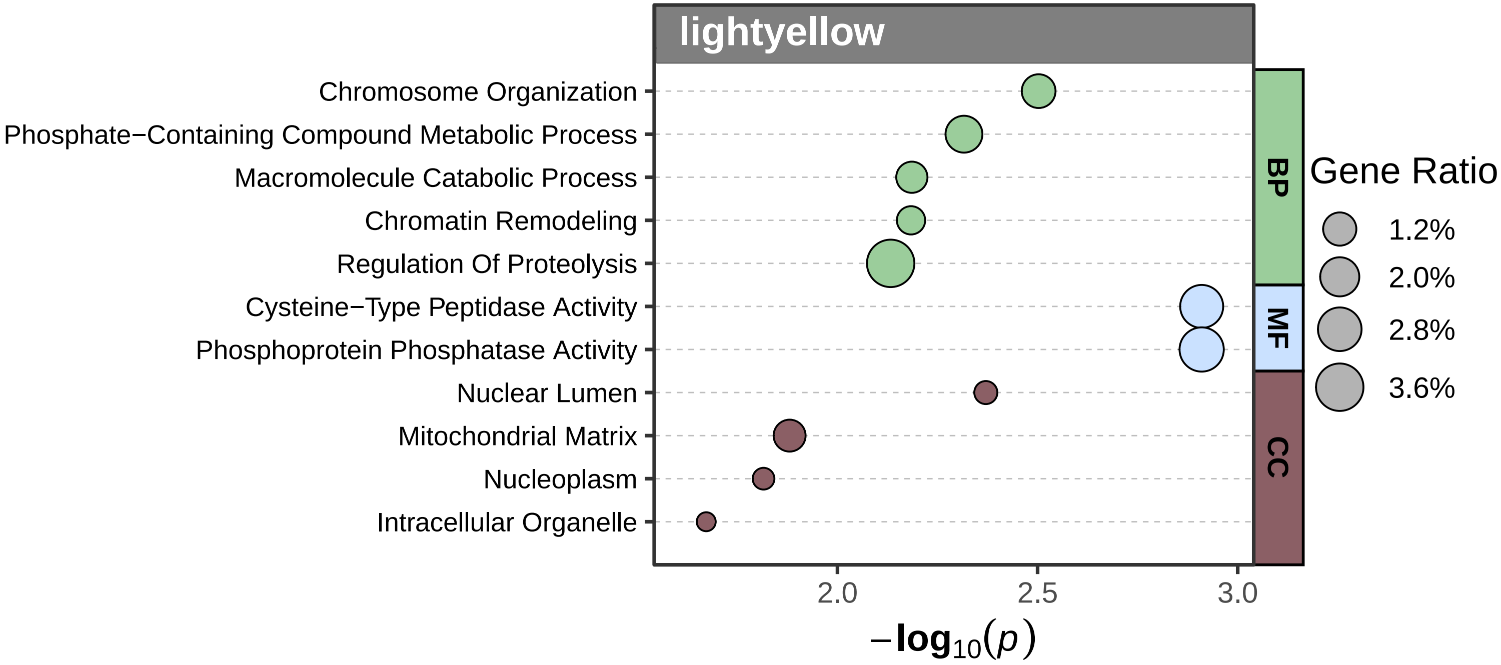

lightyellow
Chromosome Organization
Phosphate−Containing Compound Metabolic Process
Macromolecule Catabolic Process
Chromatin Remodeling
Regulation Of Proteolysis
Cysteine−Type Peptidase Activity
Phosphoprotein Phosphatase Activity
Nuclear Lumen
Mitochondrial Matrix
Nucleoplasm
Intracellular Organelle
BP
MF
CC
Gene Ratio
1.2%
2.0%
2.8%
3.6%
2.0
2.5
3.0
$-\log_{10}(p)$

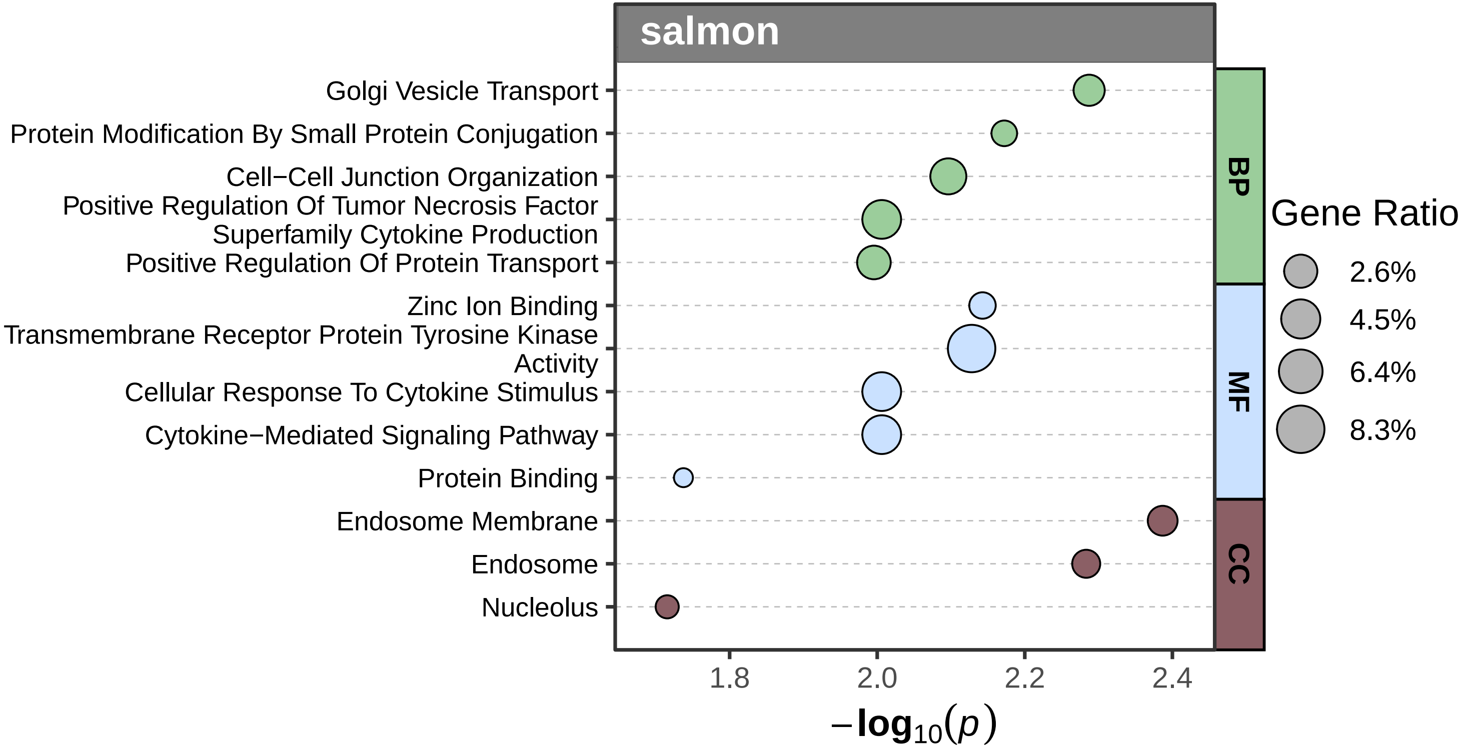
salmon
Golgi Vesicle Transport
Protein Modification By Small Protein Conjugation
Cell−Cell Junction Organization
Positive Regulation Of Tumor Necrosis Factor Superfamily Cytokine Production
Positive Regulation Of Protein Transport
Zinc Ion Binding
Transmembrane Receptor Protein Tyrosine Kinase Activity
Cellular Response To Cytokine Stimulus
Cytokine−Mediated Signaling Pathway
Protein Binding
Endosome Membrane
Endosome
Nucleolus
BP
MF
CC
Gene Ratio
2.6%
4.5%
6.4%
8.3%
1.8
2.0
2.2
2.4
$-\log_{10}(p)$

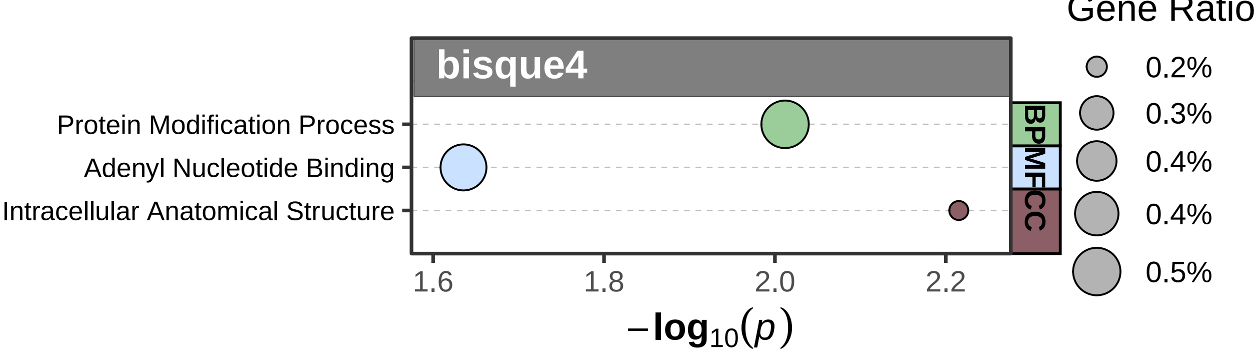

bisque4
Protein Modification Process
Adenyl Nucleotide Binding
Intracellular Anatomical Structure
BP
MF
CC
1.6
1.8
2.0
2.2
$-\log_{10}(p)$
Gene Ratio
0.2%
0.3%
0.4%
0.4%
0.5%

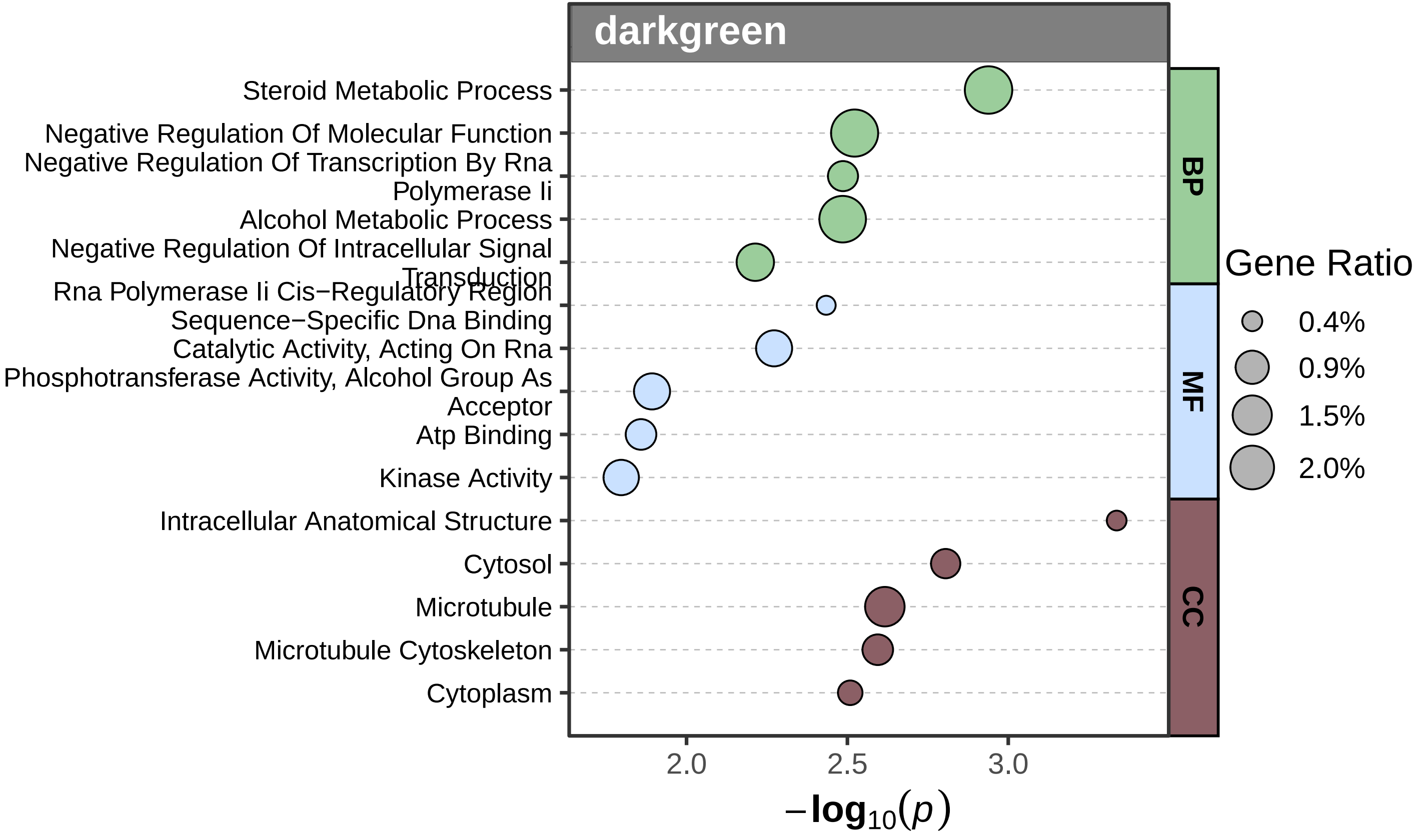

darkgreen
Steroid Metabolic Process
Negative Regulation Of Molecular Function
Negative Regulation Of Transcription By Rna Polymerase Ii
Alcohol Metabolic Process
Negative Regulation Of Intracellular Signal Transduction
Rna Polymerase Ii Cis−Regulatory Region Sequence−Specific Dna Binding
Catalytic Activity, Acting On Rna
Phosphotransferase Activity, Alcohol Group As Acceptor
Atp Binding
Kinase Activity
Intracellular Anatomical Structure
Cytosol
Microtubule
Microtubule Cytoskeleton
Cytoplasm
BP
MF
CC
Gene Ratio
0.4%
0.9%
1.5%
2.0%
2.0
2.5
3.0
$-\log_{10}(p)$

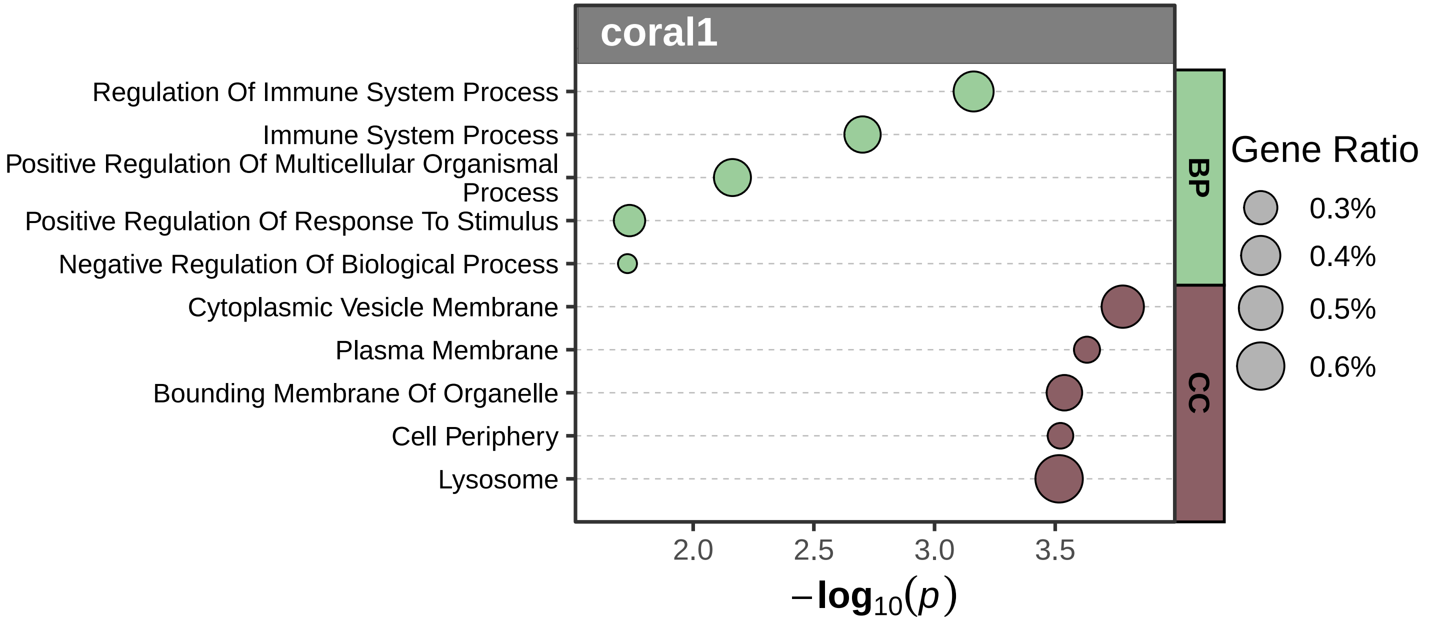

coral1
Regulation Of Immune System Process
Immune System Process
Positive Regulation Of Multicellular Organismal Process
Positive Regulation Of Response To Stimulus
Negative Regulation Of Biological Process
Cytoplasmic Vesicle Membrane
Plasma Membrane
Bounding Membrane Of Organelle
Cell Periphery
Lysosome
BP
CC
Gene Ratio
0.3%
0.4%
0.5%
0.6%
2.0
2.5
3.0
3.5
$-\log_{10}(p)$

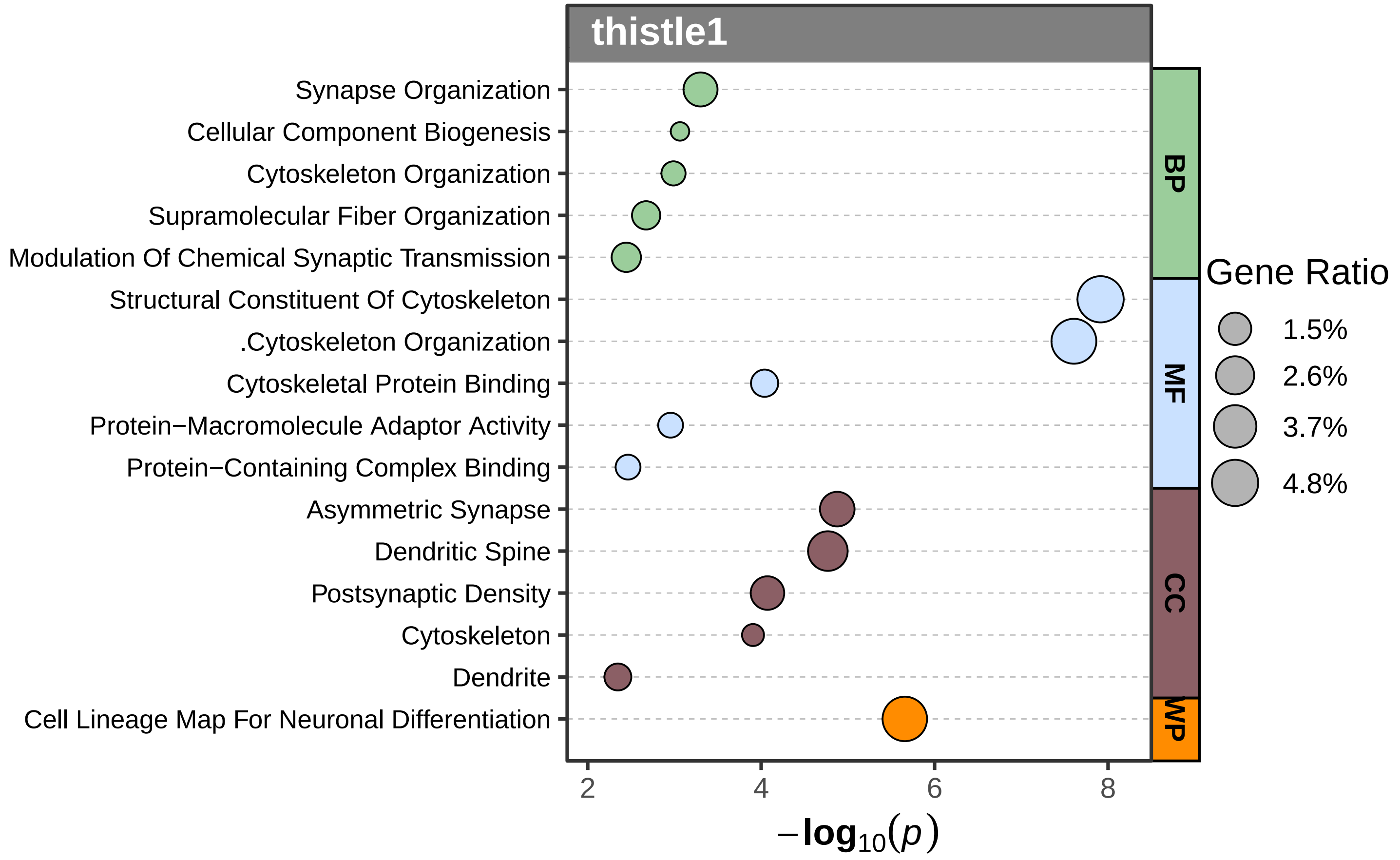
thistle1
Synapse Organization
Cellular Component Biogenesis
Cytoskeleton Organization
Supramolecular Fiber Organization
Modulation Of Chemical Synaptic Transmission
Structural Constituent Of Cytoskeleton
.Cytoskeleton Organization
Cytoskeletal Protein Binding
Protein−Macromolecule Adaptor Activity
Protein−Containing Complex Binding
Asymmetric Synapse
Dendritic Spine
Postsynaptic Density
Cytoskeleton
Dendrite
Cell Lineage Map For Neuronal Differentiation
BP
MF
CC
WP
2
4
6
8
$-\log_{10}(p)$
Gene Ratio
1.5%
2.6%
3.7%
4.8%

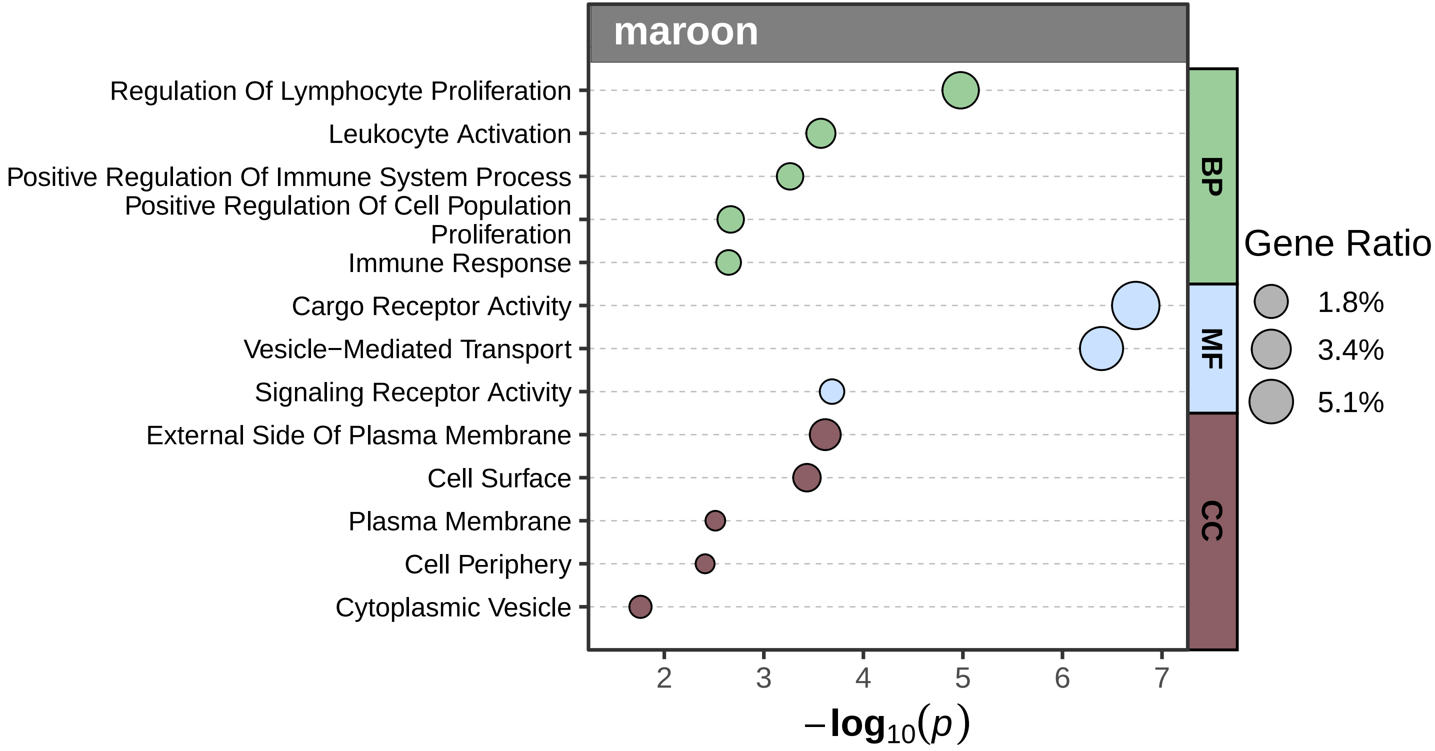

maroon
Regulation Of Lymphocyte Proliferation
Leukocyte Activation
Positive Regulation Of Immune System Process
Positive Regulation Of Cell Population Proliferation
Immune Response
Cargo Receptor Activity
Vesicle−Mediated Transport
Signaling Receptor Activity
External Side Of Plasma Membrane
Cell Surface
Plasma Membrane
Cell Periphery
Cytoplasmic Vesicle
BP
MF
CC
Gene Ratio
1.8%
3.4%
5.1%
2
3
4
5
6
7
$-\log_{10}(p)$

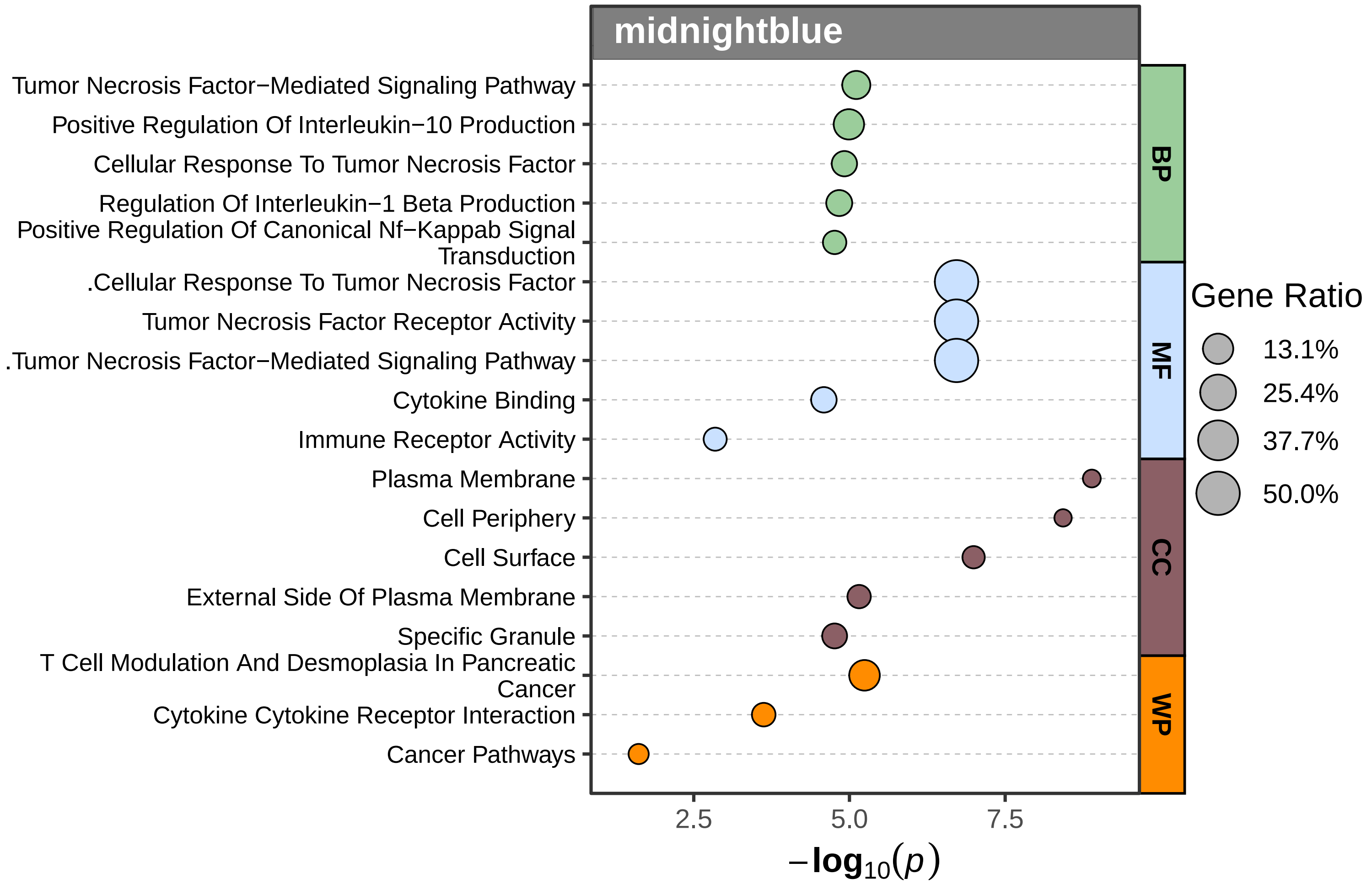
midnightblue
Tumor Necrosis Factor−Mediated Signaling Pathway
Positive Regulation Of Interleukin−10 Production
Cellular Response To Tumor Necrosis Factor
Regulation Of Interleukin−1 Beta Production
Positive Regulation Of Canonical Nf−Kappab Signal Transduction
.Cellular Response To Tumor Necrosis Factor
Tumor Necrosis Factor Receptor Activity
.Tumor Necrosis Factor−Mediated Signaling Pathway
Cytokine Binding
Immune Receptor Activity
Plasma Membrane
Cell Periphery
Cell Surface
External Side Of Plasma Membrane
Specific Granule
T Cell Modulation And Desmoplasia In Pancreatic Cancer
Cytokine Cytokine Receptor Interaction
Cancer Pathways
BP
MF
CC
WP
Gene Ratio
13.1%
25.4%
37.7%
50.0%
2.5
5.0
7.5
$-\log_{10}(p)$
